# Comprehensive Lipidomic Characterization of Human Blood by High-Throughput UHPSFC/MS using Bioinert Column and Modified MTBE Extraction

**DOI:** 10.64898/2025.12.16.694312

**Authors:** Petra Peroutková, Robert Jirásko, Anna Taylor, Michal Holčapek, Ondřej Peterka

**Affiliations:** University of Pardubice, Faculty of Chemical Technology, Department of Analytical Chemistry, Studentská 573, 532 10 Pardubice, Czech Republic

**Keywords:** Supercritical fluid chromatography, Mass spectrometry, Bioinert column, Lipidomics, Lipid extraction, Human blood

## Abstract

Supercritical fluid chromatography is traditionally employed for nonpolar and moderately polar analytes, while the analysis of ionic compounds remains a recognized limitation of the technique. Here, we introduce a novel ultrahigh-performance supercritical fluid chromatography–mass spectrometry (UHPSFC/MS) method using a bioinert column, enabling the separation of lipids with a broad polarity range from nonpolar to ionic species. The UHPSFC/MS method was optimized using 79 lipid species across 41 lipid subclasses, achieving a total run time of 7.5 min, including the column equilibration. The comparison of the separation with conventional and bioinert columns revealed a substantial improvement in peak shapes for ionic lipid classes, such as PS, LPS, PA, LPA, CerP, and SPBP. Additionally, we introduce a combination of the modified chloroform-free extraction followed by a hexane elimination step, which reduces concentrations of nonpolar lipids allowing the injection of more concentrated extracts and minimize ion source contamination. The optimized methodology was applied for the untargeted analysis of human plasma and erythrocyte-rich fraction to achieve highly confident identification of 657 lipid species across 37 lipid subclasses in human blood. The method followed the recommendations for validation of (bio)analytical methods, and its accuracy was confirmed by quantitative analysis of the reference material NIST SRM 1950, with the determined concentrations in agreement with the consensus values from ring trials. The current methodology represents a novel high-throughput and comprehensive quantitative lipidomic method for biological samples. Moreover, the findings provide the potential for the development of bioinert components for SFC platforms.

## INTRODUCTION

Lipidomics is a specialized and rapidly evolving sub-discipline of metabolomics dedicated to comprehensive, large-scale analysis that aims to identify and accurately quantify thousands of individual lipid species and also to elucidate their complex metabolic pathways and functional roles.^1,2^ Lipids are a structurally and functionally diverse group of hydrophobic or amphiphilic small biomolecules that are essential for life. Beyond their well-known roles as energy storage and the foundational structural components of cellular membranes, lipids are recognized as signaling molecules.^3,4^ Dysregulation of lipid homeostasis is directly involved in the pathogenesis of numerous serious diseases, including cardiovascular disease, diabetes, neurodegenerative disorders, and cancer.^5,6,7^ Consequently, the ability to perform high-throughput and high-resolution quantitative analyses of the lipidome is crucial for clinical applications.

The complexity of the lipidome, which potentially comprises tens of thousands of molecular species ranging from amol to nmol/mg protein, presents a significant analytical challenge.^8^ These molecules often share isobaric or isomeric compositions, making their separation and unambiguous identification difficult.^9^ Liquid chromatography coupled to mass spectrometry (LC-MS) is the gold standard in lipidomics, but it often requires long run times and a high consumption of organic solvents.^10,11^ Supercritical fluid chromatography (SFC) employs a supercritical fluid as the mobile phase, typically supercritical carbon dioxide (scCO_2_), whose density is similar to that of a liquid, providing good solvation power, while diffusion and viscosity are close to a gas.^12^ It enables fast and high efficiency separations with high flow rates and low back pressures, especially in combination with column containing sub-2 μm particles.^13,14^ However, the low polarity of the mobile phase (close to heptane) is limited for the analysis of polar lipids, requiring the use of polar modifiers (mobile phase B) consistent of polar organic solvent and volatile salts, acids, bases, and/or small amounts of water, which enable separation of nonpolar and polar lipids in one analysis.^15^

Several approaches are used in the lipidomic analysis, mainly separation techniques coupled to MS or direct infusion of the sample (without chromatographic separation) to MS, called as shotgun analysis. Shotgun represents a high-throughput quantitative approach for lipidomic analysis, but it does not allow separation of isobaric lipids at the MS1 level and carries the risk of in-source fragmentation. The most widespread approach, reversed phase (RP) LC/MS, provides detailed structural information, but faces limitations in automated data processing, requires longer analysis times, and poses challenges for accurate quantitation due to different elution of analytes and internal standards (IS), resulting in different ionization efficiency and matrix effects.^10,11^ The lipid class separation approach offers separation of isobaric lipids, co-elution of analytes and IS from the same lipid class, and highly automated data processing. On the other hand, this approach does not allow separation of some isomeric lipids and reporting structural information based on the fatty acyl/alkyl level. Moreover, the HILIC technique does not separate the nonpolar lipids eluting in the void volume and requires a longer equilibration time of the column. In comparison, SFC brings separation of nonpolar and polar lipids in one analysis, high chromatographic resolution, minimal organic solvent consumption, and short analysis times.^12^

However, some lipids may contain a chromatographically challenging ionic group, especially the phosphate moieties, which can interact with the metal surfaces of the instrument and column, resulting in the poor peak shape and loss of sensitivity. Bioinert materials, such as polyether ether ketone, titanium, Hastelloy, or MP35N, represent a crucial advancement for those analytes in omics analyses.^16,17,18^ The aim of the study was to develop a high-throughput, comprehensive, and accurate quantitative methodology using bioinert material for the lipidomic analysis of human blood, including highly automated data processing.

## EXPERIMENTAL SECTION

### Chemicals and standards

Methanol (MeOH), 2-propanol (IPA), hexane, butanol, acetic acid, and formic acid (all LC/MS gradient grade) and ammonium carbonate (≥ 30.0% NH_3_ basis) were purchased from Honeywell (Riedel-de Haën, Germany). Hydrochloric acid (35%, p.a.) was obtained from PENTA s.r.o. (Prague, Czech Republic) and ammonium acetate (≥99.99%) from Sigma-Aldrich (St. Louis, MO, USA). LiChrosolv solvents, chloroform stabilized with 2-methyl-2-butene, methyl *tert*-butyl ether (MTBE), and ammonium formate (LC-MS grade) were purchased from Merck (Darmstadt, Germany). Water (LC-MS grade) used for the mobile phase was purchased from Th. Geyer GmbH & Co. (Renningen, Germany). Carbon dioxide of 4.5 grade (99.995%) was obtained from Messer (Bad Soden, Germany). Deionized water used for extraction procedures was prepared by the Milli-Q Reference Water Purification System (Molsheim, France). Lipid standards and IS were purchased from Avanti Polar Lipids (Alabaster, AL, USA), Nu-Chek Prep (Elysian, MN, USA), Cayman Chemical (Ann Arbor, MI, USA), or Merck (Darmstadt, Germany). All stock solutions of lipid standards were prepared in MeOH/CHCl_3_ (1:1, *v/v*) and stored at -80 °C. Deuterated IS are typically supplied in chloroform solution, which can be used directly as a stock solution. Mixtures of standards (Std-Mix) and internal standards (IS-Mix) (**Tables S1-S4**) were prepared by combining aliquots of individual lipid stock solutions and subsequently diluted with MeOH/CHCl_3_ (1:1, v/v) mixture to obtain the final desired concentrations.

### Biological samples

All samples were stored at −80 °C prior to lipidomic extraction. All volunteers in the study were over 18 years of age and blood samples were collected from each participant after an overnight fast. The pooled plasma sample was employed for optimization of the extraction procedure, the untargeted analysis, and validation, while the pooled erythrocyte-rich fraction was used only for the untargeted analysis. This study was performed in accordance with the Declaration of Helsinki, and all collected data were pseudonymized. Informed consent was obtained from all volunteers, and the ethical committee approved the collection of blood samples. The NIST SRM 1950 human plasma served as a reference material for quantitative analysis and comparison with literature values.

### Sample preparation

For the MTBE liquid–liquid extraction, 25 µL of plasma, or alternatively a 25 µL of erythrocyte-rich fraction, and 20 µL of IS-Mix (**Table S3**) were pipetted into a 4 mL glass vial. To this, 0.7 mL of methanol and 2.3 mL of MTBE were added, and the vial was placed in an ultrasonic bath for 15 min at 30°C. After the mixture was cooled to ambient temperature, 0.6 mL of 250 mM ammonium carbonate (adjusted to pH 5 with acetic acid) was added. The mixture was subsequently stirred at 560 rpm (KS 130 shaker, IKA, Staufen, Germany) at ambient temperature for 5 min. Next, the mixture was centrifuged (Hettich EBA 20) for 5 min at 6,000 rpm (3,462×g). The MTBE (upper) layer was then carefully transferred into an 8 mL glass vial using a glass pipette to form the initial fraction. For re-extraction, 2.3 mL of MTBE was added to the remaining aqueous layer, and the mixture was stirred at 560 rpm at ambient temperature for 5 min, centrifuged for 5 min at 6,000 rpm (3,462×g), and the resulting MTBE layer was transferred and combined with the previous fraction. The combined MTBE phase was evaporated under a gentle stream of nitrogen at 35°C. Finally, 1 mL of hexane and 1 mL of the mixture of MeOH/H_2_O (98:2, *v/v*) were added to the residue, vortexed for 30 seconds, and centrifuged for 5 min at 6,000 rpm (3,462×g). The hexane (upper) layer was transferred to waste, and the remaining methanol phase was evaporated under the gentle stream of nitrogen at 35°C. Before analysis, the residue was dissolved in 500 µL of the CHCl_3_/MeOH mixture (1:1, v/v) and 10 times diluted extract for positive ion mode and extract without dilution for negative ion mode were injected. An overview of the extraction procedure is visualized in **Figure 1**.

**Figure 1.**
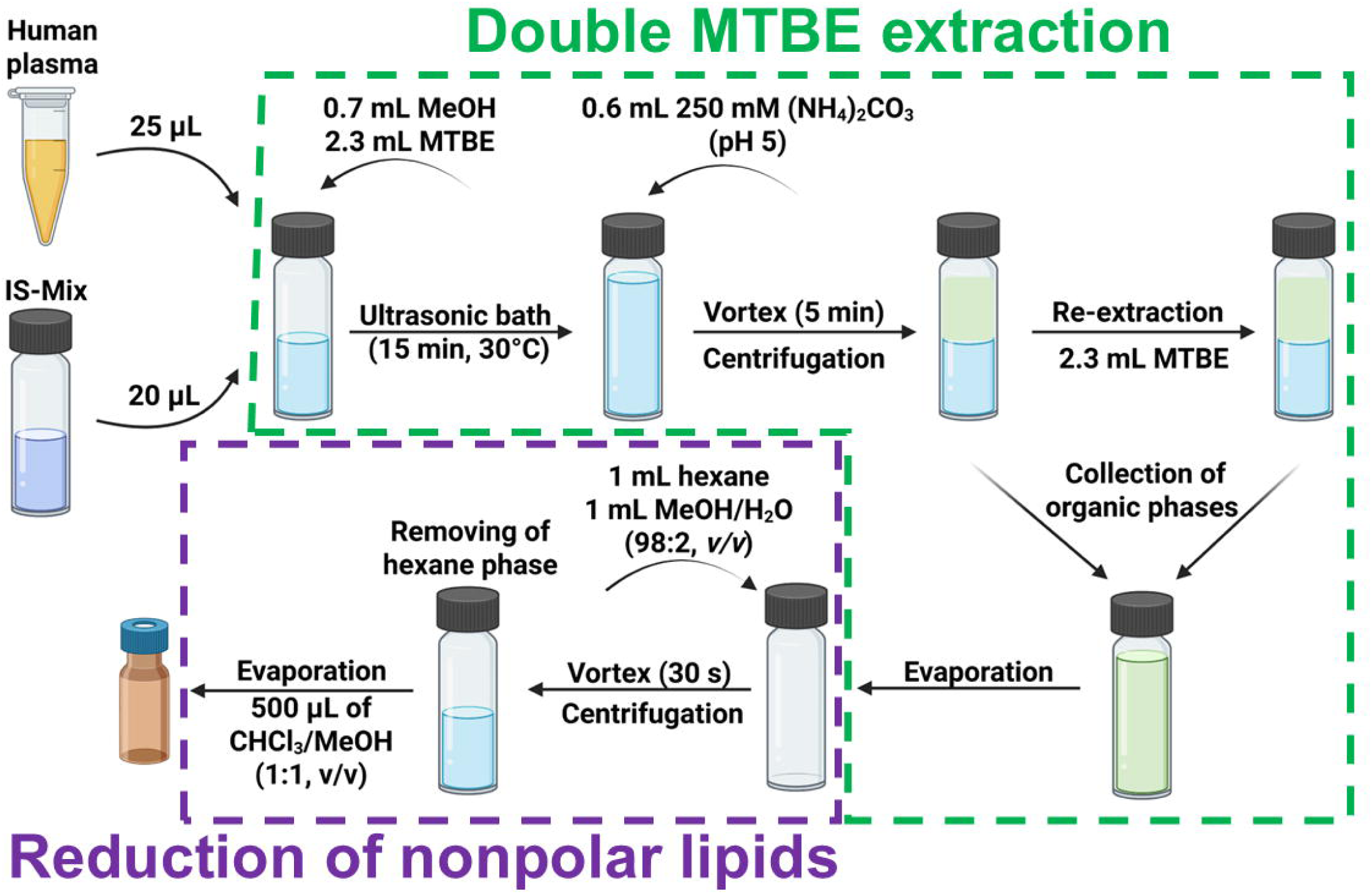
Overview of the double MTBE extraction procedure and the subsequent reduction of nonpolar lipid concentration using a hexane extraction step for human plasma. Figure was created using BioRender.

### UHPSFC/MS conditions

UHPSFC analysis was performed on an Acquity UPC2 instrument from Waters (Milford, MA, USA) using the Acquity Premier BEH HILIC column (100 × 2.1 mm, 1.7 µm). The following linear gradient was employed using scCO_2_ (mobile phase A) and MeOH containing 30 mM ammonium acetate in methanol with 1% water used as a modifier (mobile phase B): 0 min, 1% B; 5 min, 55% B; 7 min, 55% B; 7.1 min, 1% B; 7.5 min, 1% B. The total run time of the method was 7.5 min, including the column equilibration. The column temperature was set to 60 °C, the flow rate to 0.9 mL/min, the automatic back-pressure regulator to 1800 psi, and the injection volume was 1 μL. The needle was washed after each injection with IPA/hexane/H_2_O (4:2:1, *v/v/v*).

The UHPSFC was coupled to the hybrid quadrupole–time of flight (QTOF) mass spectrometer Synapt G2-Si (Waters). The make-up solvent had the identical composition as the modifier with the flow rate of 0.1 mL/min. The following parameters were set for MS measurements: sensitivity mode, electrospray ionization (ESI) in positive or negative ion mode, the mass range of *m/z* 200–1600, the capillary voltage of 3 kV for positive and -2 kV for negative ion mode, the sampling cone of 20 V, the source offset of 90 V, the source temperature 150 °C, the cone gas flow of 50 L/h, the desolvation gas flow of 1000 L/h, and the nebulizer gas flow of 4 bar. The desolvation temperature was set to 500 °C for positive ion mode and 600 °C for negative ion mode. The analysis was done in the continuum mode with the scan time of 0.5 s. The peptide leucine enkephalin was used as the lock mass. The reporting checklist^19^ of the method is attached in **ESM 2**.

### Data processing

The raw data from UHPSFC/MS analysis were processed for noise reduction, *m/z* correction using the lock mass value, and conversion from continuum to centroid mode by the Accurate Mass Measure tool in MassLynx 4.1. The peak areas used for method optimization were exported by TargetLynx based on the predefined mass window tolerance of ±20 mDa and the retention time tolerance of ±0.2 min. The manual identification was performed in the full scan mode by matching theoretical and experimental values within a defined mass tolerance window (±10 ppm) using an in-house lipid database. Moreover, retention times of individual lipid classes were confirmed by standards, together with the dependencies of retention time on the carbon number or the number of double bonds within a lipid class, were used for highly confident identification. The lipid concentrations in NIST SRM 1950 human plasma were calculated using LipidQuant 2.1 software,^20^ and type I and type II isotopic corrections were automatically applied. The input data for individual lipid classes were generated by MarkerLynx, including experimental *m/z* values with MS signal responses. The response factors for cholesterol esters (CE) were calculated as the ratio of the calibration curve slope of CE 16:0 D7 and individual standards. Visualizations were prepared in the R software environment (ver. 4.4.1).^21^

### Method validation

The calibration curves were prepared based on measurements of 14 concentration levels in triplicates and evaluated for positive and negative ion modes separately. The limit of detection (LOD) and limit of quantification (LOQ) values were determined by direct experimental measurements, rather than being theoretically estimated from the calibration curve parameters. Accuracy, precision, extraction recovery, and matrix effects were evaluated using human plasma spiked by IS-Mix based on 3 concentration levels. Low, medium, and high concentration levels corresponding to 10, 20, and 40 μL of the IS-Mix, respectively, and all parameters were evaluated based on at least four independent replicates. Intra- and inter-day accuracy and precision were assessed using six independent samples. Instrument precision was investigated using IS without a matrix. The selectivity of IS was examined using randomly selected plasma samples from three males and three females, and also for pooled plasma. The carry-over was investigated using solvent blanks injected after pooled plasma spiked with the highest concentration levels of calibration curves (50 μL of IS-Mix). Extraction recovery was determined by the comparison of the signal intensity of IS spiked before and after the extraction at three concentration levels. Matrix effects were assessed by comparing the signal intensity of IS spiked after the extraction with responses of neat IS at three concentration levels.

## RESULTS AND DISCUSSION

### Optimization of UHPSFC/MS method

Although SFC is primarily associated with the separation of nonpolar and less polar analytes, a modifier improves the separation of polar analytes.^15^ The composition of the modifier is essential for chromatographic separation, but the gradient elution is limited by the pressure limit of the instrument and by the need to maintain supercritical or subcritical conditions of the mobile phase.^12,13^ The initial conditions of the method were inspired by the method described by Wolrab et al.,^22^ which led to the use of a BEH stationary phase and the mobile phase consisting of 30 mM ammonium acetate in methanol with 1% water. The gradient slope was recalculated under the conditions of the current column. For the evaluation of peak shapes enhancement using a bioinert column (ACQUITY Premier BEH HILIC, 100 × 2.1 mm; 1.7 µm), a conventional column (ACQUITY BEH HILIC, 100 × 2.1 mm, 1.7 µm) under comparable conditions was used for the analysis of the same Std-Mix (**Table S1**). A significant improvement was observed for phosphatidylserines (PS), lysophosphatidylserines (LPS), phosphatidic acids (PA), lysophosphatidic acids (LPA), ceramide-phosphate (CerP), and sphingoid base-phosphate (SPBP), visualized in **Figure 2**, while the results for other lipid classes were comparable. Due to the lack of commercially available bioinert SFC columns, the bioinert column primarily intended for HILIC was used. Afterwards, the flow rate of the mobile phase and the slope of the gradient were optimized based on 79 lipid species from 41 lipid subclasses (**Table S2**), resulting in the separation method with the total run time of 7.5 min including the column equilibration. Moreover, the method resolves the isomeric lipids (**Figure S1**), such as glucosylceramides (GlcCer) *vs.* galactosylceramides (GalCer) and phosphatidylglycerols (PG) *vs.* bis(monoacylglycerol)phosphates (BMP).

**Figure 2.**
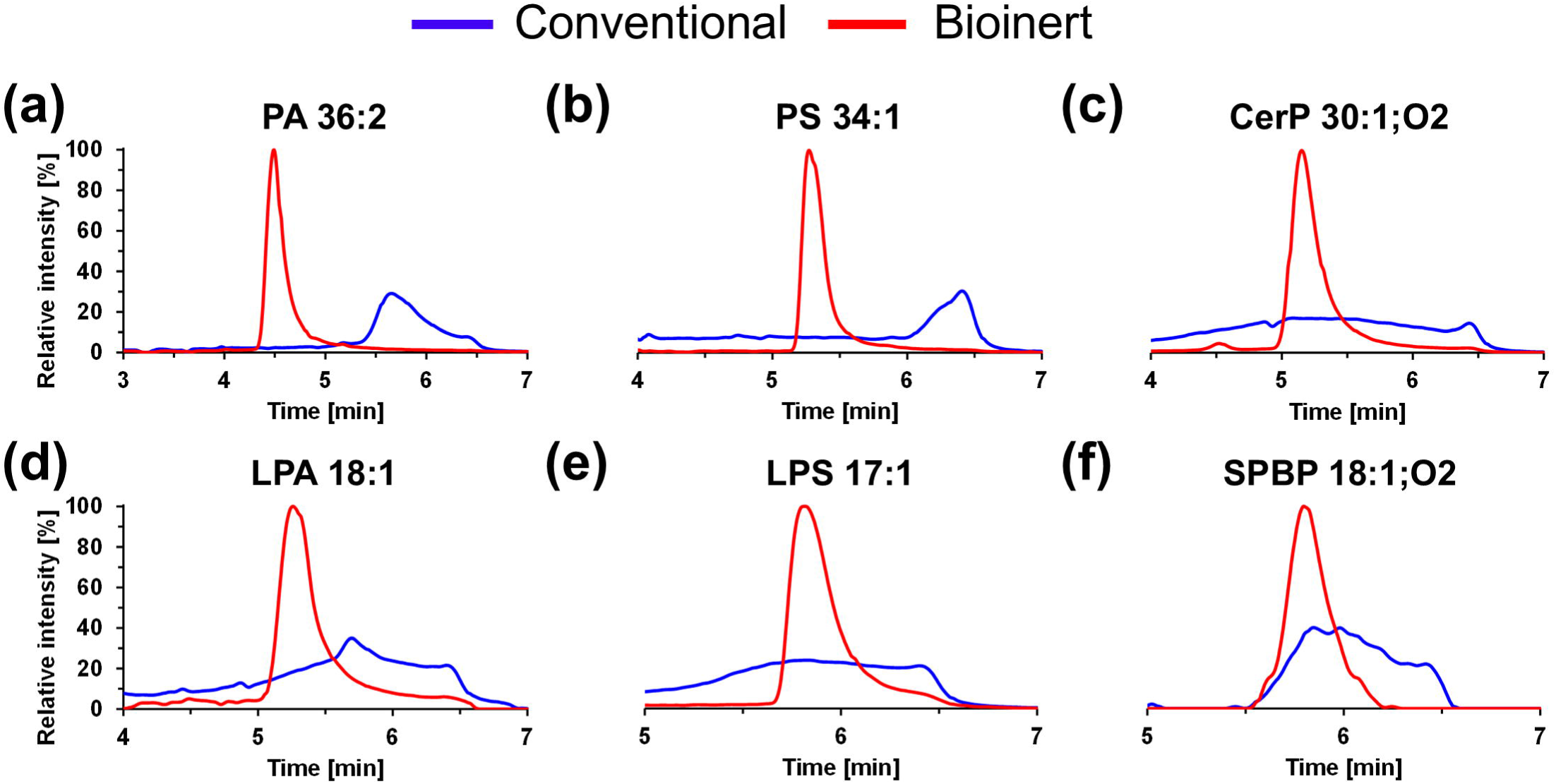
Comparison of the bioinert (ACQUITY Premier BEH HILIC, 100 × 2.1 mm; 1.7 µm) and the conventional (ACQUITY BEH HILIC, 100 × 2.1 mm; 1.7 µm) columns based on the peak shapes, illustrated for the lipid standards of **(a)** PA 36:2, **(b)** PS 34:1, **(c)** CerP 30:1;O2, **(d)** LPA 18:1, **(e)** LPS 17:1, and **(f)** SPBP 18:1;O2.

Subsequently, the flow rate of the makeup solvent (the same composition as the modifier) and the MS parameters (source and desolvation temperatures) were optimized based on the signal intensities of the lipid standards (**Table S2**) to achieve high ionization efficiency. Mostly two standards from one lipid class were included to evaluate the potential effect of the fatty acyl chain length, but both follow the same trends. The makeup flow rates of 0.1, 0.2, 0.25, 0.3, and 0.4 mL/min were examined, and 0.1 mL/min provided the best results for most investigated lipid classes (**Figure S2**). However, the makeup flow rate had a greater effect in the negative ion mode. The source (100, 125, and 150 °C) and desolvation (400, 500, and 600 °C) temperatures were optimized separately for positive and negative ion modes, and both temperatures significantly affected ionization efficiency (**Figures S3 and S4**). Higher temperatures resulted in increased ionization efficiency for most of the investigated lipid classes, but higher temperatures also promoted the formation of [M−H₂O+H]⁺ adduct at the expense of other adducts for some lipid classes in the positive ion mode, such as monoacylglycerols (MG), diacylglycerols (DG), free sterols (ST), ceramides (Cer), hexosylceramides (HexCer), and CerP. Moreover, higher temperature increased in-source fragmentation of CE in the positive ion mode. To minimize these undesirable effects, source and desolvation temperatures of 100 °C and 400 °C, respectively, would be optimal. However, these settings significantly decrease the ionization efficiency for other lipid classes, therefore 150 °C for source temperature and different desolvation temperatures for positive (500 °C) and negative (600 °C) ion modes were selected as the best compromise for most lipid classes. A higher number of simultaneously formed adducts were observed in the positive ion mode compared to the negative mode, where ionization predominantly yielded one preferential adduct (**Figures S3 and S4**). However, both [M-H]^-^ and [M+CH_3_COO]^-^ ions were observed for some lipid classes. Comparable intensities of both [M–H]⁻ and [M+CH₃COO]⁻ ions were observed exclusively for NAE, while for Cer, HexCer, Hex2Cer, and Hex3Cer these adducts were also detected but with a markedly lower response. Base peak intensity chromatograms of Std-Mix measured by the optimized method in both polarities are shown in **Figure 3**.

**Figure 3.**
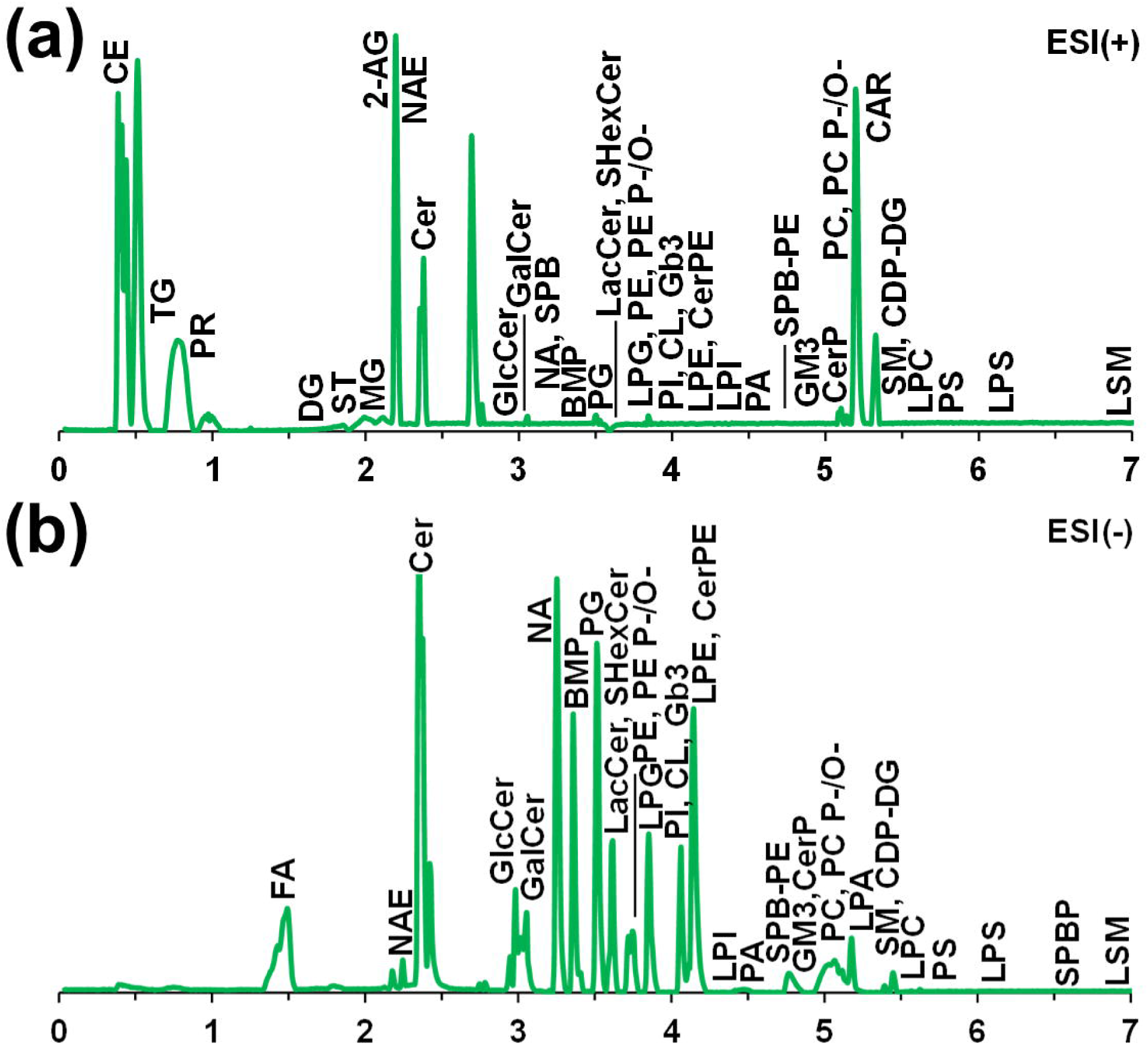
Base peak chromatograms of lipid standard mixture acquired in **(a)** positive and **(b)** negative ion modes.

The performance of the developed UHPSFC/MS method was compared with previously published methods in terms of their chromatographic conditions (**Table S5**). Compared to other intra-laboratory^22,23^ and inter-laboratory^24,25,26^ UHPSFC/MS methods, current method separates more lipid classes in a shorter total run time. Although the total run time of most methods is 8–12 min, they require higher mobile phase flow rates due to the dimensions of SFC columns (i.d. 3.0 × 100 mm) and higher flow rates of the makeup solvent, resulting in approximately two times higher solvent consumption. On the other hand, a smaller column diameter (i.d. 2.1 × 100 mm) adversely affected the peak shape of the first-eluting CE and triacylglycerols (TG), while the peak shape of other lipid classes remained unaffected. Compared to HILIC/MS methods representing an alternative LC approach, the UHPSFC/MS method enables significantly shorter run times. Although the total analysis time of some HILIC/MS methods is around 10 min,^22,27^ it is usually 20 min and longer.^23,25,28^ Moreover, the UHPSFC/MS method separates a greater number of lipid classes and allows the simultaneous analysis of both nonpolar and polar lipids. Although the flow rate is lower in HILIC/MS, the overall consumption of organic solvents is higher due to the longer run time and the relatively high organic solvent content in the mobile phase(s). The trend has been confirmed by Tomiyasu et al. in their direct comparison of both techniques.^25^

### Optimization of extraction procedure

The sample preparation is a critical step in lipidomic analysis, as improper handling can lead to sample degradation and the formation of various artifacts. Based on our previous study,^22^ a modified double Folch extraction procedure was selected as the initial extraction method. Because this extraction method had been evaluated mainly for neutral lipids, an additional optimization, including various additives and adjustments to the pH of the aqueous phase, was performed. In addition to the originally used 250 mM ammonium carbonate (pH close to 9 without adjustment), aqueous phase without additives or mainly additives compatible with MS were examined. The replacement of ammonium carbonate with ammonium acetate or ammonium formate, followed by pH adjustment using acetic acid, formic acid, and HCl, was investigated, because acidic environment is more suitable for the extraction of ionic lipids, such as LPA, PA, CerP, and SPBP.^29^ The significantly lowest extraction recoveries were observed for the extraction without any additives, especially for polar and ionic lipids. The use of 250 mM ammonium carbonate without pH adjustment provided results comparable to other variants for most lipid classes, but limitations were observed for LPS, LPA, LPI, and SPBP. For other variants containing 250 mM of additives adjusted to an acidic pH, the extraction recoveries were comparable for most lipid classes. Although adjusting the pH from basic to acidic is uncommon, the 250 mM ammonium carbonate solution adjusted to pH 5 with acetic acid or pH 4 with formic acid yielded the best extraction recoveries. For further optimization, 250 mM ammonium carbonate adjusted to pH 5 with acetic acid was selected due to slightly better results. The effect of individual combinations of additives based on extraction recoveries is visualized in **Figure S5**.

Over the years, several extraction protocols have been developed, mainly using single-phase (BuMe^30^) and two-phase extraction (Folch,^31^ Bligh-Dyer,^32^ and Matyash^33^) methods, but each approach provides some limitations. Single-phase extractions lead to high matrix effects, while two-phase extractions can result in the distribution of lipids between the organic and aqueous phases depending on their structure, particularly affecting highly polar and ionic lipids.^34^ These commonly used extraction methods were evaluated based on extraction recoveries of 79 lipid species from 41 lipid subclasses, by comparing the signal intensities of IS spiked into the plasma matrix before and after extraction. Based on previous optimization and to unify the extraction conditions, the re-extraction step and 250 mM ammonium carbonate at pH 5 were included in two-phase extractions. A detailed description of other extraction methods are provided in **ESM 1**. However, the Folch extraction without the re-extraction step (single Folch) was also performed to investigate the improvement of the re-extraction step, which showed higher extraction recoveries for all investigated lipid classes. The extraction recoveries for the single Folch extraction were mainly between 70–80%, but there were lipid classes with recoveries lower than 50%, such as phosphatidylethanolamines (PE), lysophosphatidylethanolamines (LPE), LPS, lysophosphatidylinositols (LPI), lysosphingomyelins (LSM), SPBP, and sphingosyl phosphoethanolamine (SPB-PE). In comparison, the inclusion of the re-extraction step increased extraction recoveries to 85–108% and by approximately 20% for the problematic lipid classes. The double Bligh–Dyer extraction provided comparable results to the double Folch procedure, except for LPS, lysophosphatidylglycerols (LPG), LPA, LPI, LSM, and SPBP, which showed lower values for the double Bligh–Dyer method. The protein precipitation approach demonstrated consistent extraction recoveries across all lipid classes, with recoveries mainly between 80–90%, except for the lowest values (50% and 53%, respectively) observed for SPBP and LPA. However, the best results were observed for the MTBE extraction procedure using MTBE as an alternative to chloroform-based extractions. The extraction recoveries were mainly between 90–100% for all investigated lipid classes, except for LPS and LSM, which showed recoveries of 55% and 57%, respectively. The comparison of extraction recoveries for individual extraction methods is illustrated in **Figure 4**.

**Figure 4.**
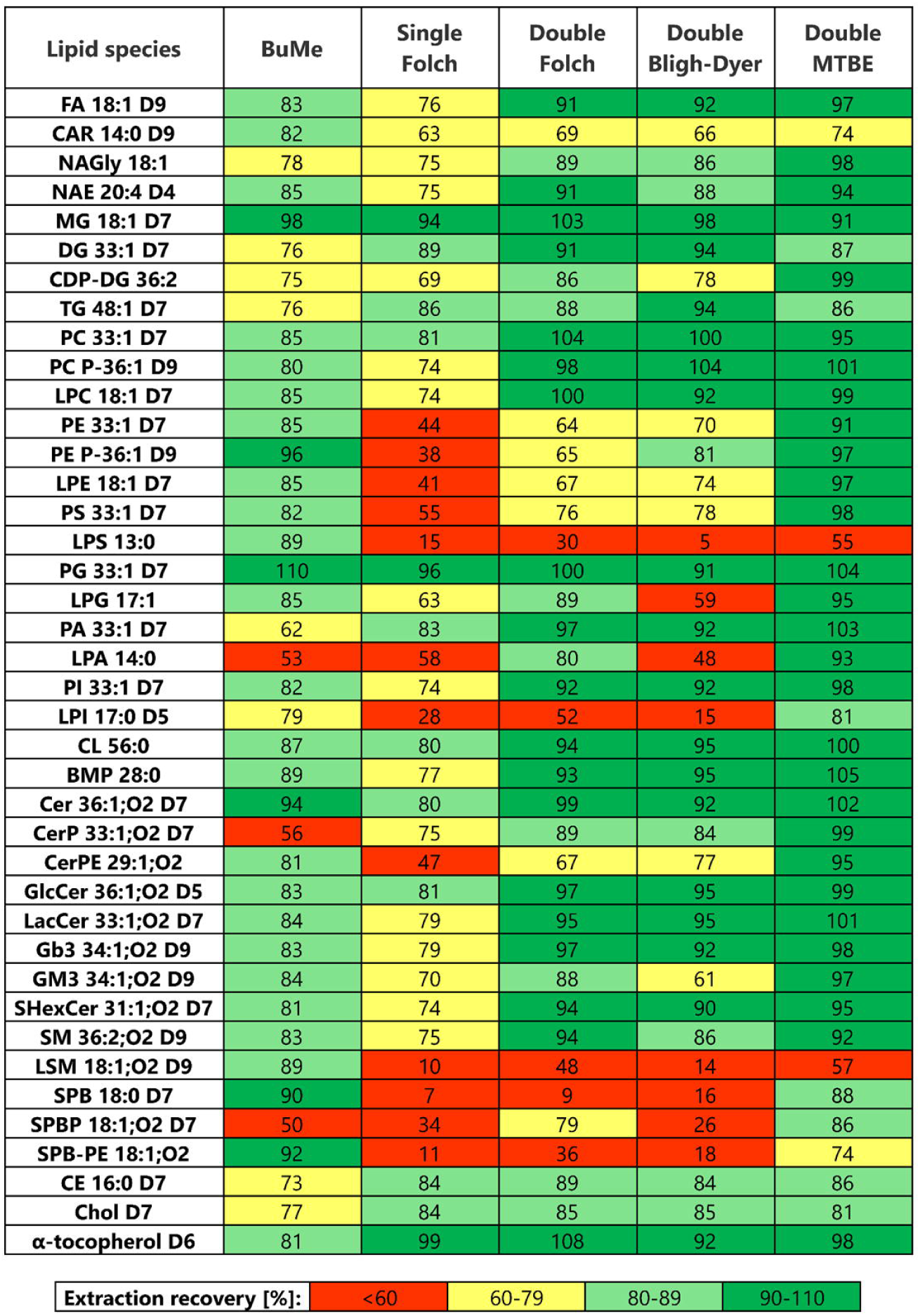
Comparison of extraction recoveries for single-phase extraction (BuMe protein precipitation) and two-phase extractions (Folch, Bligh–Dyer, and MTBE). The double extraction procedure included a re-extraction step. Extraction recovery was calculated by comparing the signal intensity of IS spiked before and after the extraction. Values represent the average from three independent experiments.

Plasma contains high concentrations of nonpolar lipids, such as TG and CE, which leads to a higher dilution factor in the positive ion mode and increased contamination of the MS ion source. The protocol was modified to obtain high extraction recovery for polar and ionic lipids, while nonpolar lipids were reduced by hexane extraction step. The two-phase extraction, including MeOH/H_2_O (98/2, *v/v*) and hexane phases, was inspired by our previous studies using hexane extraction for the lipid fractionation^35^ and the selective extraction of nonpolar lipids^36^. Compared to previous protocols, the volume of hexane was reduced from 2 to 1 mL to allow only a reduction of their concentrations rather than complete removal, so that the analysis of these classes could be maintained. However, this hexane elimination step is not necessary for matrices containing lower concentrations of nonpolar lipids. The extraction recoveries of the final extraction method are described in the method validation.

### Identification of lipids in human blood

The optimized methodology was used for the untargeted analysis of human blood, separately for the human plasma and the erythrocyte-rich fraction. The individual lipid species were identified based on the combination of several parameters, such as mass accuracy, confirmation of retention times of individual lipid classes by standards, and retention dependencies. Although lipids from the same lipid classes elute in one chromatographic peak, there is a partial separation within the lipid class based on the number of double bonds and the carbon number.^37^ The retention time is increased with the higher number of double bonds for all lipid classes and with the higher carbon number for non- and less-polar lipid classes (sterol esters (SE), TG, DG, MG, Cer, and fatty acids (FA)), while the higher carbon number for polar lipids results in lower retention (**Figure S6**). Lower retention was also observed for sphingolipids compared to hydroxylated sphingolipids from the same lipid class, providing two independent retention dependencies. However, the changes in retention times caused by the number of double bonds and the carbon number in high-speed UHPSFC separations are smaller than in HILIC^28^ and substantially differ to RP-UHPLC^38^.

The semi-automated identification was performed using LipidQuant 2.1 software,^20^ which utilizes an in-house database of lipids, but each feature was manually verified, resulting in a list of identifications (**Table S6**). However, the co-elution of high-abundance phosphatidylcholines (PC) and low-abundance LPA does not allow identification of LPA due to in-source fragmentation. Although PA can also be produced by in-source fragmentation of phospholipids, the current method separates PA from all phospholipid classes (**Figure S7A**), resulting in the identification of PA in real samples. The separation of individual sphingolipid classes (**Figure S7B**) also helps to eliminate misidentifications caused by in-source fragmentation, particularly due to the loss of sugar unit(s). Although Hex2Cer and SHexCer elute close together, there are small differences in retention times for individual lipid species with the same fatty acyl composition. Furthermore, the current method enables the separation of isomeric lipid classes, such as GlcCer *vs.* GalCer, PG *vs.* BMP, or 1,2-DG *vs.* 1,3-DG (**Figure S1**), where the retention orders of individual isomers were confirmed by standards. Base peak chromatograms in positive and negative ion modes for plasma and the erythrocyte-rich fraction are shown in **Figure 5**. The shorthand nomenclature for lipids was used according to Liebisch et al.^39^ and used abbreviations are summarized in **ESM 1**.

**Figure 5.**
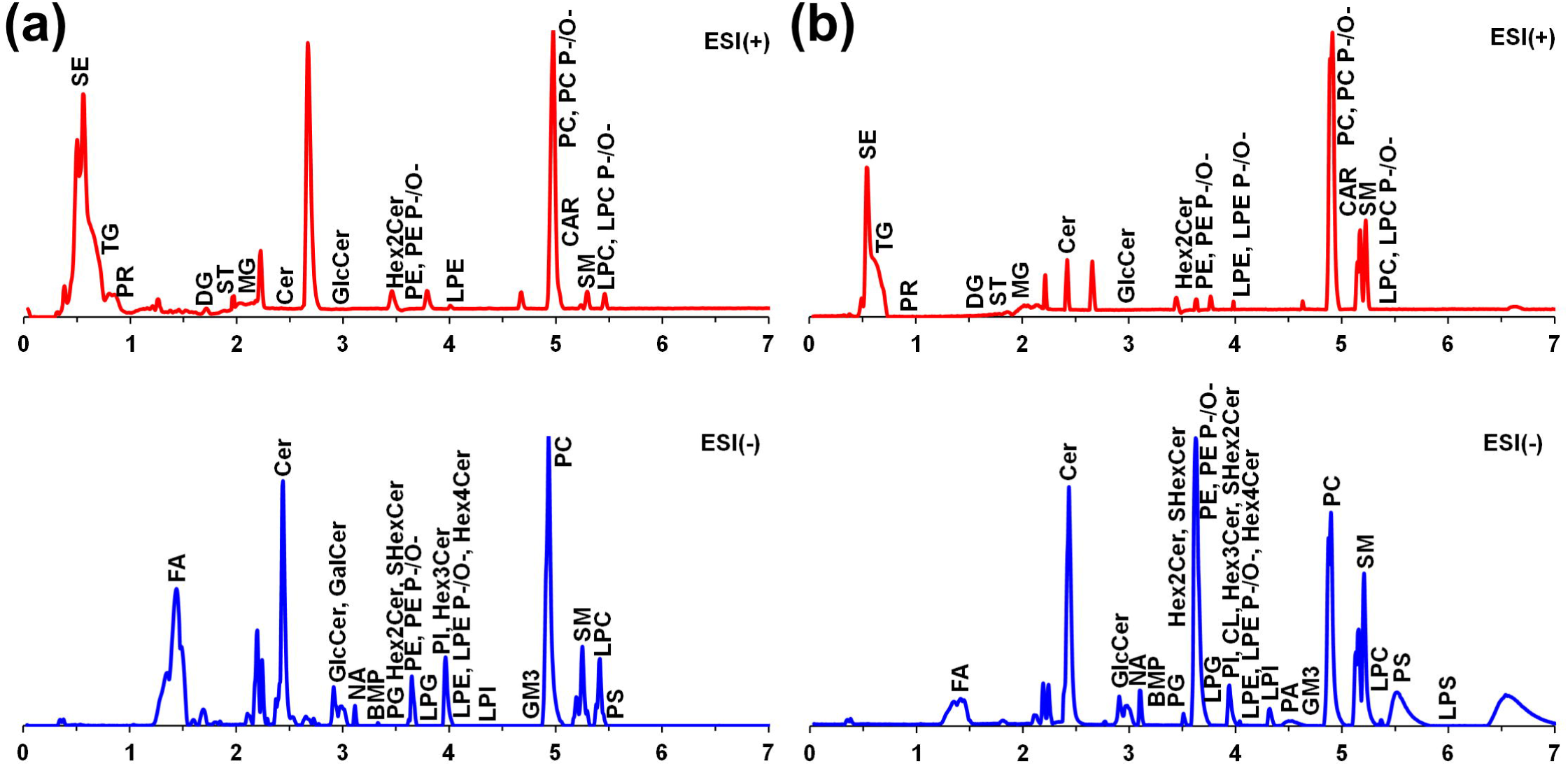
Base peak chromatograms of lipid extracts from **(a)** human plasma and **(b)** erythrocyte-rich fraction acquired in both positive (red) and negative (blue) ion modes.

In total, 657 lipid species from 37 lipid subclasses were identified in the human blood (**Table S6**), whereas 477 lipid species from 32 lipid subclasses in plasma and 575 lipid species from 34 lipid subclasses in the erythrocyte-rich fraction were detected. The negative ion mode yielded more lipid identifications than the positive ion mode (351 *vs.* 228 for plasma and 473 *vs.* 224 for the erythrocyte-rich fraction), with 102 lipid species in plasma and 122 in the erythrocyte-rich fraction detected in both polarities (**Figure S8**). The comparison of identified lipid species demonstrates partial differences between both matrices (**Figure 6**), with 395 lipid species identified in both matrices, while 82 lipid species were observed only in plasma and 180 only in the erythrocyte-rich fraction (**Figure S8**). More phospholipids and sphingolipids were detected in the erythrocyte-rich fraction, especially PE, PS, PA, PG, trihexosylceramide (Hex3Cer), and tetrahexosylceramide (Hex4Cer), while nonpolar lipids such as TG, DG, and SE are more abundant in plasma.

**Figure 6:**
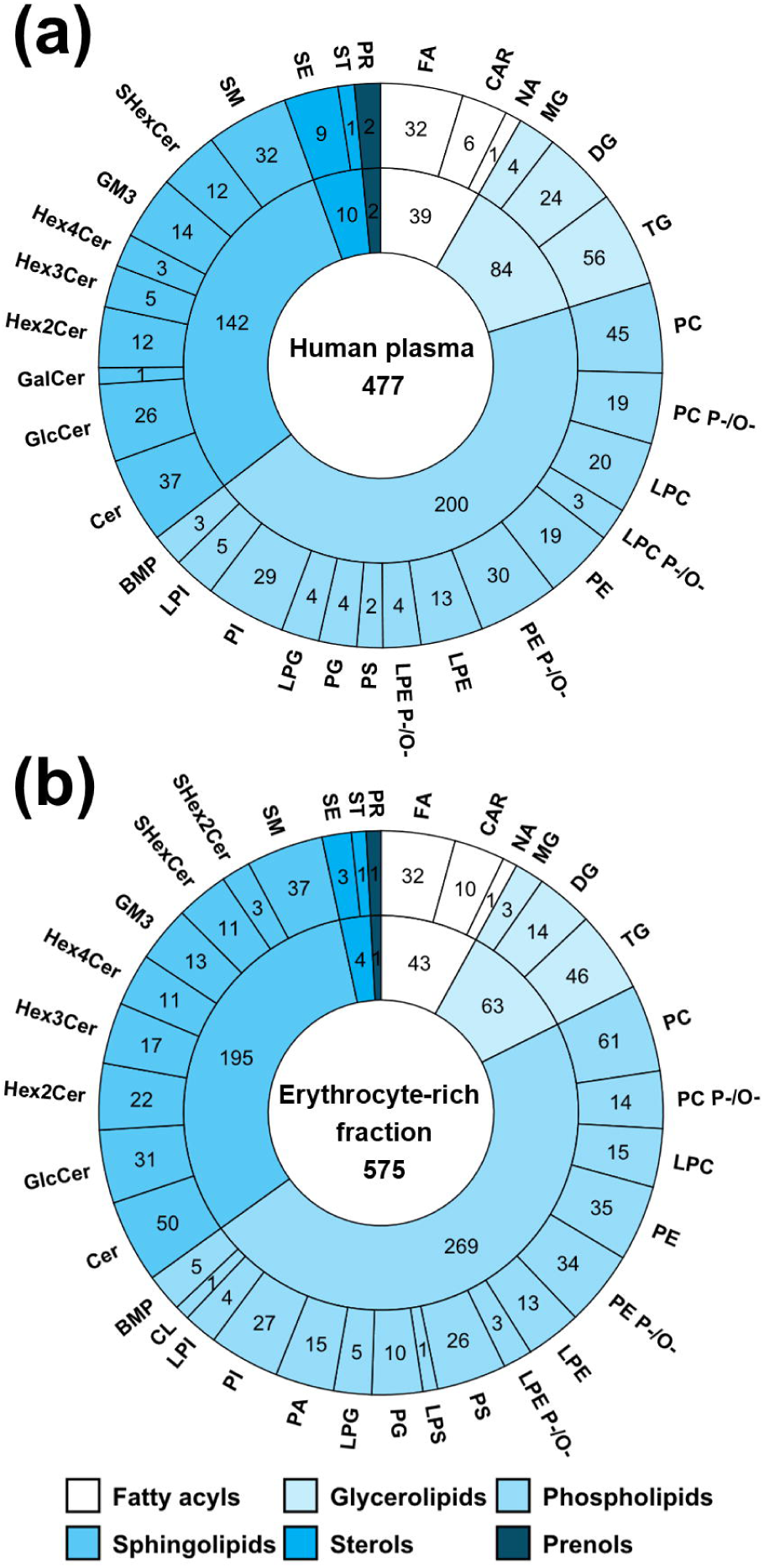
Comparison of numbers of identified lipid species from individual lipid classes in **(a)** human plasma and **(b)** erythrocyte-rich fraction.

The intra-laboratory comparison using the same UHPSFC/MS (QTOF) platform clearly showed the benefits of the current methodology, which led to a significantly higher number of identified lipid species in the plasma sample (477 *vs.* 126 and 227, respectively)^22,23^ and the erythrocyte-rich fraction (575 *vs.* 173)^23^ during a shorter total run time (7.5 *vs.* 8 and 10 min, respectively)^22,23^. However, the previous UHPSFC/MS methods performed analysis only in the positive ion mode and Wolrab et al.^22^ reported quantitative results. The intra-laboratory comparison with HILIC/MS (QTOF) methods showed that the current UHPSFC/MS methodology provided a significantly higher number of lipid identifications in both plasma (477 *vs.* 132, 52, and 239, respectively)^22,23,28^ and erythrocyte-rich fraction (575 *vs.* 64) in shorter run time. The inter-laboratory comparison using various UHPSFC/MS methods confirmed the improvement of the current methodology for lipidomic analysis. Although the previous methods were also performed in both ion modes, the current method enabled the identification of a higher number of lipid species in human plasma using the QTOF platforms (477 *vs.* 165 and 168, respectively)^24,26^ and the QQQ platforms (477 *vs.* 307 and 182, respectively)^24,25^. A similar trend was also observed in the inter-laboratory comparison with the HILIC/MS methods using the QQQ platform in both ion modes (477 *vs.* 152)^25^ and the Orbitrap platform in the positive ion mode (477 *vs.* 341)^27^. However, the inter-laboratory methods^25,26,27^ reported quantitative results. Detailed intra- and inter-laboratory comparison based on the number of identified lipid species is summarized in **Table S7**.

### Method validation for human plasma

The methodology was validated before the quantitative analysis following recommendations for (bio)analytical methods.^22^ The validation was carried out for the individual IS, and two IS were used for almost every lipid class. In total, 69 IS from 37 lipid subclasses, including deuterated and exogenous lipid species with shorter, longer, or odd carbon fatty acyls, were used (**Table S3**). The calibration curves provided linear regression coefficients greater than 0.99 for all investigated IS and the calibration ranges, LOD, and LOQ for individual IS are summarized for both positive and negative ion modes in **Table S8**. Carnitines (CAR), MG, DG, TG, CE, and ST were investigated only in the positive ion mode, while FA, *N*-acyl ethanolamines (NAE; endocannabinoids), PS, PG, LPG, phosphatidylinositols (PI), LPI, cardiolipin (CL), BMP, CerPE, Hex3Cer, gangliosides (GM3), and sulfatides (SHexCer) in the negative ion mode due to their ionization efficiencies. PC, lysophosphatidylcholines (LPC), PE, LPE, Cer, HexCer, Hex2Cer, sphingomyelins (SM), and prenols (PR) were evaluated in both ion modes. The method was further validated for several other parameters, including accuracy, precision, selectivity, instrument precision, and matrix effect (**Tables S9-S10**). All IS were considered suitable for the quantitative analysis of human plasma, with the exception of DG 36:2 D5 in the positive ion mode and SM 36:2 D9 in the negative ion, for which the second exogenous IS was used due to poor selectivity. A detailed description of individual parameters is provided in **ESM 1**.

The extraction recoveries were investigated using matrices of four males and four females at three concentration levels and the comparison of the signal intensity of IS spiked before and after extraction demonstrated high extraction yields. However, the hexane elimination step was intended to reduce the extraction recovery of high-abundance nonpolar lipids, yielding average recoveries of 13% for TG and 9% for CE. This elimination step also influenced the extraction recovery of DG, ST, and PR, resulting in average recoveries of 51%, 60%, and 75%, respectively. Although the extraction recoveries of the non-polar classes are low, all other validation criteria were satisfied, allowing their quantitative analysis. An average extraction recovery of 80% was observed for LPI, which was attributed to its distribution between the organic and aqueous phases. For all other lipid classes, the extraction recoveries were within ±15%.

### Quantitative analysis of NIST SRM 1950

Finally, the optimized and validated methodology was used for the quantitative analysis of the reference material NIST SRM 1950 human plasma to verify its accuracy of the current method. Although the current method enables the baseline separation of isomers, such as GlcCer vs. GalCer or 1,2-DG vs. 1,3-DG, our data processing for quantitative analysis is based on combined mass spectra of the entire lipid class.^20^ Consequently, only the sum of isomers is determined and reported as HexCer or DG without further specification. In total, 239 lipid species from 25 lipid subclasses were quantified in human plasma. The positive ion mode contributed to the quantification of 111 lipid species from 14 subclasses and the negative ion mode to 181 lipid species from 17 subclasses (**Figure S8D**). The concentrations of lipid species determined in both polarities (LPC, PC, PE, Cer, and SM) were compared (**Figure S9**) and all lipid classes provided high correlation (R^2^ = 0.97). Moreover, the response factors were established for SE (**Table S11**) to eliminate large differences in responses depending on the composition of fatty acyls, such as the number of double bonds and the length of the acyl chains.^40,41^ The response factors were calculated as the ratio of calibration curve slope of CE 16:0 D7 to that of individual standards.

The determined concentrations were compared with the literature values from previous ring trials (**Figure 7**).^42,43,44,45^ The strongest correlations were found with the results of Bowden et al.^42^ (R² = 0.90) and Ghorasaini et al.^44^ (R² = 0.85). Moderate correlations were observed with the studies by Quehenberger et al.^43^ and Mandal et al.^45^ (R² = 0.66 and 0.71, respectively). However, the deviations were particularly observed for PE, PE P-/O-, and LPE in Quehenberger et al. and for FA in Mandal et al., which are also among the other ring trials (**Figure S10**). After the exclusion of these outlying classes, the methods also showed high correlations, with R^2^ values of 0.82 and 0.86, respectively. On the other hand, the determined concentrations of PI using the current method were approximately two times higher compared to the ring trials. The determined concentrations of lipid species by the current method and the detailed comparison with ring trials are listed in **Table S12**. Minor differences in concentrations can be caused by different platforms, sample preparation, but particularly by various analytical approaches and IS composition. The current method utilizes a lipid class separation approach that ensures co-elution of IS with analytes compared to the lipid species separation approach preferred in ring trials. Moreover, the current method employs structurally similar deuterated IS and individual deuterated IS for PC P-/O- and PE P-/O-, unlike other methods, which typically use IS of PC and PE for quantitation of these subclasses. The combination of highly automated data processing, deuterated IS for each lipid class, type I and type II isotopic corrections, and response factors for SE ensures that the method meets the analytical requirements for high-throughput and accurate lipidomic quantitation.

**Figure 7.**
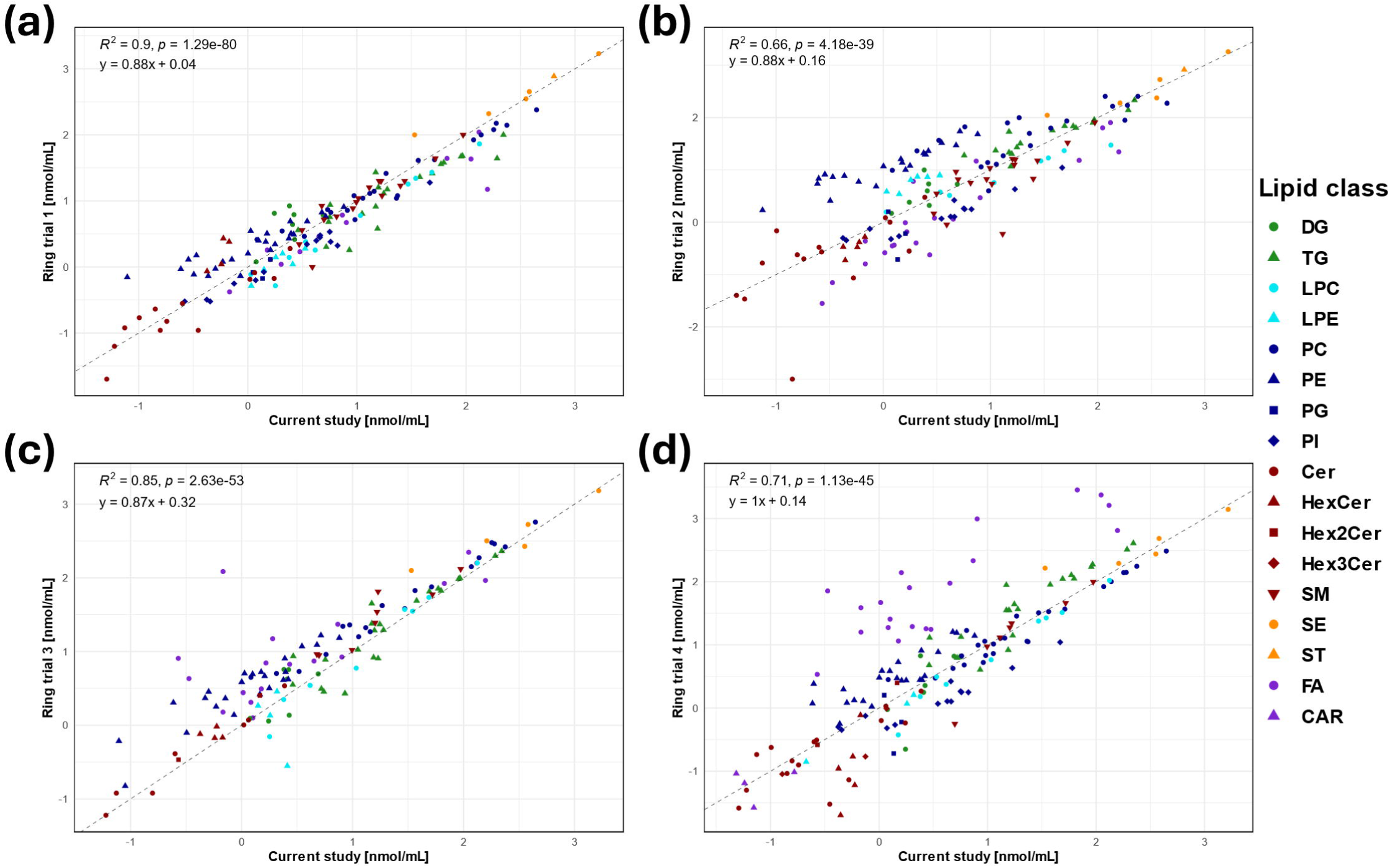
Comparison of determined concentrations in NIST SRM 1950 human plasma using the current method and four inter-laboratory ring trials using correlation graphs: **(a)** Bowden et al. (Ring trial 1),^42^ **(b)** Quehenberger et al. (Ring trial 2),^43^ **(c)** Ghorasaini et al. (Ring trial 3),^44^ and **(d)** Mandal et al. (Ring trial 4).^45^

## CONCLUSIONS

In this work, we introduced a novel high-throughput UHPSFC/MS method employing a bioinert column, whose use in SFC has not been previously reported. Combined with a modified MTBE extraction procedure, this methodology enabled the highly confident identification of 657 lipid species in human blood and quantification of 239 lipid species in human plasma. The innovative use of the bioinert column in the UHPSFC/MS method shows significant improvement in the analysis of chromatographically challenging ionic lipid classes, such as PS, LPS, PA, LPA, CerP, and SPBP. Although the bioinert LC column was applied for the SFC method, we demonstrate the potential for developing bioinert components specifically designed for SFC, which are not yet commercially available. The new UHPSFC/MS method also significantly decreases the consumption of organic solvents due to lower flow rates of the mobile phases and the makeup solvent, resulting in an overall reduction of about 50% compared to the previous method. The combination of the UHPSFC/MS and the modified extraction also led to minimized ion source contamination, allowing the measurement of approximately 3–5 times more samples between the deep MS cleaning steps compared to the previous method. Moreover, the implementation of a chloroform-free extraction and the reduction of organic solvent consumption contribute to the principles of green chemistry.

## ASSOCIATED CONTENT

Supporting Information. The Supporting Information is available free of charge on the ACS Publications website at DOI: XXX.

Electronic Supplementary Material (ESM) 1: Abbreviations, Description of additional extraction methods, Method optimization, Extraction optimization, Chromatograms, Venn diagrams, and Correlation graphs. (Word file)

Electronic Supplementary Material (ESM) 2: Reporting Checklist (PDF)

Electronic Supplementary Material (ESM) 3: Composition of Std-Mix and IS-Mix, Comparision with other methods, List of identification, Validation results, Response factors, and Quantitation results. (Excel file)

## Supporting information

Reporting Checklist

## ACKNOWLEDGMENTS

This work was supported by the project NU21-03-00499 sponsored by the Czech Health Research Council and the ERC Adv grant No. 101095860 (European Research Council). We would like to thank prof. Bohuslav Melichar from Palacký University and University Hospital Olomouc, Czech Republic for the arrangement of ethical approval for biological samples.

## COMPLIANCE WITH ETHICAL STANDARDS

All volunteers signed an informed consent, and the ethics committee approved the blood collection.

---

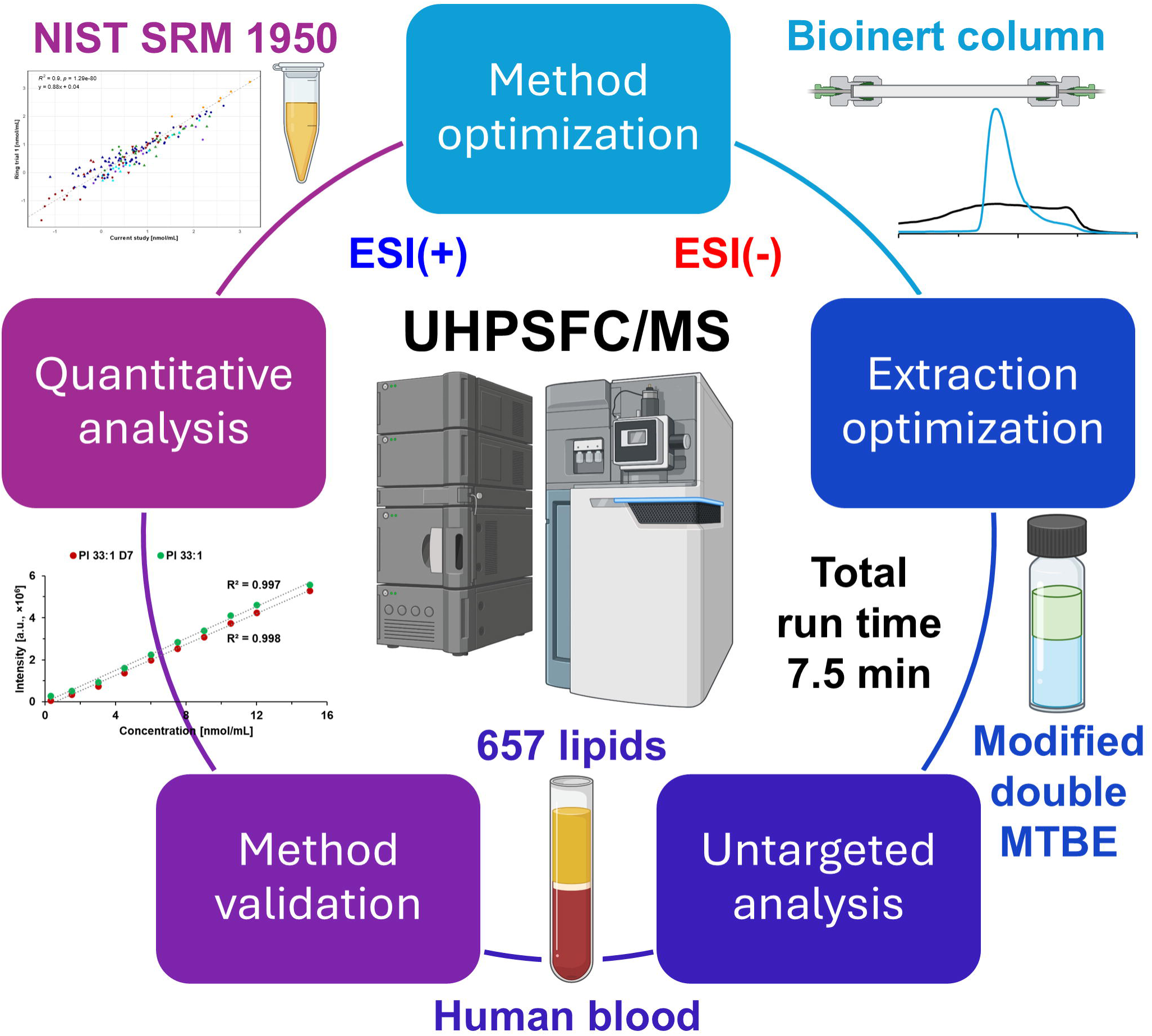

