## Supplementary Figures for "Comprehensive Lipidomic Characterization of Human Blood by High-Throughput UHPSFC/MS using Bioinert Column and Modified MTBE Extraction"

**Electronic Supplementary Material 1**

**Table of content:**

[**Figure S10:** Correlation graphs for NIST SRM 1950 human plasma comparing results from individual ring trials: Bowden et al. (Ring trial 1) [4], Quehenberger et al. (Ring trial 2) [5], Ghorasaini et al. (Ring trial 3) [6], and Mandal et al. (Ring trial 4) [7].	34](#_Toc216471273)

Abbreviations
**2-AG** – 2-Arachidonoylglycerol

**BMP** – Bis(monoacylglycerol)phosphate

**BuMe** – Butanol/Methanol

**CAR** – Carnitine

**CDP-DG** – Cytidine diphosphate diacylglycerol

**CE** – Cholesterol ester

**Cer** – Ceramide

**CerP** – Ceramide-phosphate

**CerPE** – Ceramide phosphoethanolamine

**Chol** – Cholesterol

**CL** – Cardiolipin

**DG** – Diacylglycerol

**ESI** – Electrospray ionization

**FA** –Fatty acid

**GalCer** – Galactosylceramide

**Gb3** – Globotriaosylceramide

**GlcCer** – Glucosylceramide

**GM3** – Ganglioside

**HexCer** – Hexosylceramide

**Hex2Cer** – Dihexosylceramide

**Hex3Cer** – Trihexosylceramide

**Hex4Cer** – Tetrahexosylceramide

**HILIC** – Hydrophilic interaction liquid chromatography

**IPA** – 2-propanol

**IS-Mix** – Mixtures of internal standards

**LacCer** – Lactosylceramide

**LC-MS** – Liquid chromatography coupled with mass spectrometry

**LOD** – Limit of detection

**LOQ** – Limit of quantification

**LPA** – Lysophosphatidic acid

**LPC** – Lysophosphatidylcholine

**LPE** – Lysophosphatidylethanolamine

**LPG** – Lysophosphatidylglycerol

**LPI** – Lysophosphatidylinositol

**LPS** – Lysophosphatidylserine

**LSM** – Lysosphingomyelin

**MeOH** – Methanol

**MG** – Monoacylglycerol

**MS** – Mass spectrometry

**MTBE** – Methyl tert-butyl ether

**NA** - *N*-acyl amines

**NAE** – *N*-acyl ethanolamines (Endocannabinoids)

**NAGly** – *N*-arachidonoyl glycine

**-O** – Alkyl bond

**-P** – Plasmalogen

**PA** – Phosphatidic acid

**PC** – Phosphatidylcholine

**PE** – Phosphatidylethanolamine

**PG** – Phosphatidylglycerol

**PI** – Phosphatidylinositol

**PR** – Prenol

**PS** – Phosphatidylserine

**QTOF** – Quadrupole–time of flight

**RP** – Reversed phase

**scCO2** – Supercritical carbon dioxide

**SE** – Sterol ester

**SFC** – Supercritical fluid chromatography

**SHex2Cer** – Dihexosyl sulfatide

**SHexCer** – Sulfatide

**SM** – Sphingomyelin

**SPB** – Sphingoid base

**SPB-PE** – Sphingosyl phosphoethanolamine

**SPBP** – Sphingoid base-phosphate

**Std-Mix** – Mixtures of standards

**ST** – Free sterol

**TG** – Triacylglycerol

**UHPSFC/MS** – Ultrahigh-performance supercritical fluid chromatography–mass spectrometry

### Extraction protocols

**Folch extraction [1].** Initially, 25 µL of plasma and 20 µL of IS-Mix were pipetted into a 4 mL glass vial. To this, 1 mL of methanol and 2 mL of CHCl_3_ were added, and the vial was placed in an ultrasonic bath for 15 min at 30 °C. After the mixture was cooled to ambient temperature, 0.6 mL of 250 mM ammonium carbonate (adjusted to pH 5 with acetic acid) was added. Subsequently, the mixture was stirred (560 rpm, IKA KS 130 shaker) at ambient temperature for 5 min. Next, the mixture was centrifuged for 5 min at 6,000 rpm (3,462×g). The organic layer (lower) was then carefully transferred to a new 4 mL glass vial using a glass pipette. The organic fraction was evaporated under a gentle stream of nitrogen at 35 °C. Before analysis, the residue was dissolved in 500 µL of the CHCl_3_/MeOH mixture (1:1, *v/v*) and carefully vortexed for 30 seconds.

**Double Folch extraction [1].** At first, 25 µL of plasma and 20 µL of IS-Mix were pipetted into a 4 mL glass vial. To this, 1 mL of methanol and 2 mL of CHCl_3_ were added, and the vial was placed in an ultrasonic bath for 15 min at 30 °C. After the mixture was cooled to ambient temperature, 0.6 mL of 250 mM ammonium carbonate (adjusted to pH 5 with acetic acid) was added. The mixture was subsequently stirred (560 rpm, IKA KS 130 shaker) at ambient temperature for 5 min. The mixture was centrifuged for 5 min at 6,000 rpm (3,462×g). The CHCl_3_ layer (lower) was then carefully transferred into an 8 mL glass vial using a glass pipette to form the initial fraction. For re-extraction, 2 mL of CHCl_3_ were added to the remaining aqueous layer and the mixture was stirred (560 rpm, IKA KS 130 shaker) at ambient temperature for 5 min and centrifuged for 5 min at 6,000 rpm (3,462×g). The resulting CHCl_3_ layer was transferred and combined with the previous fraction. The combined CHCl_3_ phase was evaporated under a gentle stream of nitrogen at 35 °C. Before analysis, the residue was dissolved in 500 µL of the CHCl_3_/MeOH mixture (1:1, *v/v*) and carefully vortexed for 30 seconds.

**Double Bligh-Dyer extraction [2].** At the beginning, 25 µL of plasma and 20 µL of IS-Mix were pipetted into an 8 mL glass vial. To this, 2 mL of methanol and 1 mL of CHCl_3_ were added, and the vial was placed in an ultrasonic bath for 15 min at 30 °C. After the mixture was cooled to ambient temperature, 0.8 mL of 250 mM ammonium carbonate (adjusted to pH 5 with acetic acid) was added. The mixture was placed in an ultrasonic bath for 5 min at 30 °C for homogenization. The mixture was centrifuged for 5 min at 6,000 rpm (3,462×g) and left to settle for 15 min. The CHCl_3_ layer (lower) was then carefully transferred into a new 8 mL glass vial using a glass pipette to form the initial fraction. For re-extraction, 1 mL of CHCl_3_ was added to the remaining aqueous layer and the mixture was placed in an ultrasonic bath for 5 min at 30°C, centrifuged for 5 min at 6,000 rpm (3,462×g), and left to settle for 15 min. The resulting CHCl_3_ layer was transferred and combined with the previous fraction. The combined CHCl_3_ phase was evaporated under a gentle stream of nitrogen at 35 °C. Before analysis, the residue was dissolved in 500 µL of the CHCl_3_/MeOH mixture (1:1, *v/v*) and carefully vortexed for 30 seconds.

**Protein precipitation using BuMe [3].** Initially, 25 µL of plasma and 20 µL of IS-Mix were pipetted into a 4 mL glass vial. To this, 0.25 mL of a butanol/methanol mixture (1:1, *v/v*) was added, and the vial was placed in an ultrasonic bath for 15 min at 30 °C. After the mixture was cooled to ambient temperature, 0.5 mL of the butanol/methanol mixture (1:1, *v/v*) was added to eliminate the risk of trapping analytes in the filter. The mixture was subsequently stirred (560 rpm, IKA KS 130 shaker) at ambient temperature for 5 min. Next, the mixture was centrifuged for 5 min at 6,000 rpm (3,462×g). The samples were filtered through a 0.2 µm cellulose filter (OlimPeak, Teknokroma, Spain). The remaining phase was evaporated under a gentle stream of nitrogen at 35 °C. Before analysis, the residue was dissolved in 500 µL of the CHCl_3_/MeOH mixture (1:1, *v/v*) and carefully vortexed for 30 seconds.

### Method validation

The lower LOD and LOQ in the negative ion mode were affected by the different sample dilutions (200 times for positive and 20 times for negative ion modes equivalent to plasma), but lipids containing choline in the structure (PC, LPC, and SM) provided higher ionization efficiency in positive ion mode. Instrument precision was below 10% for all internal standards evaluated at three concentration levels, ensuring the reproducibility of measurements. For accurate quantitation, precision and accuracy provide the key parameters. Precision and accuracy are key parameters for reliable quantitation and the method was evaluated by intra- and inter-day precision and accuracy using six independent samples. Both parameters were below 20% for almost all IS, except for PE in the positive ion mode due to lower sensitivity, and therefore, PE quantification was performed in the negative ion mode. Higher values were observed also for TG and CE in positive ion mode, but they were still below 20%. Selectivity, a critical parameter for the evaluation of IS, was examined in six different plasma samples, with all IS meeting the criteria except for DG 36:2 D5 in the positive ion mode and SM 36:2 D9 in the negative ion mode, which was considered unsuitable for the quantitative analysis of human plasma by this method and the second exogenous IS was used for the quantitative analysis. Although liquid–liquid extraction was employed, matrix-related ion suppression and enhancement were still present. The values were mainly within ±20% or very close to this interval, but a higher ion enhancement was observed for PC, LPC, and LPE in positive ion mode, and for CerPE, PI, and LPI in negative ion mode. Importantly, these effects were consistent, and higher matrix effect values were not considered an exclusion criterion.

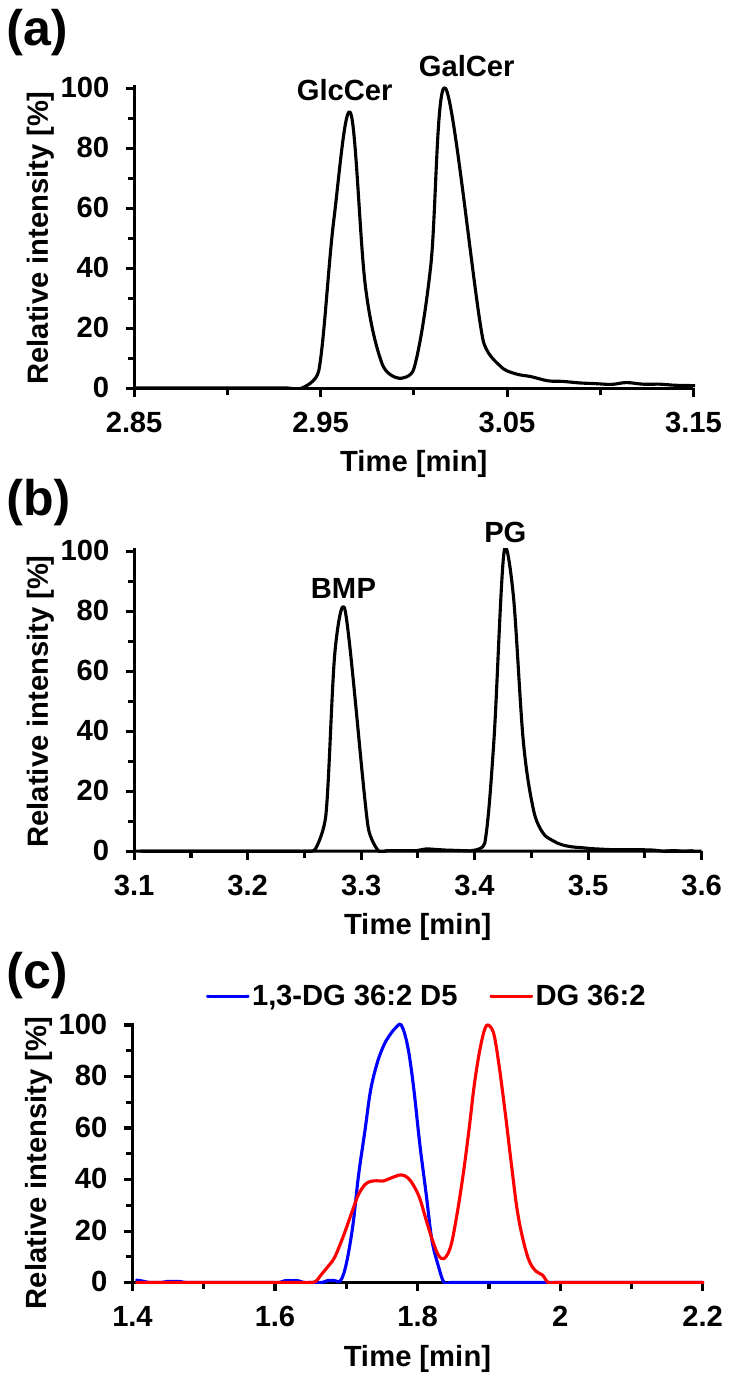

**Figure S1**: Separation of lipid isomers: **(a)** GlcCer *vs.* GalCer, **(b)** BMP *vs.* PG, and **(c)** 1,3-DG *vs.* 1,2-DG.

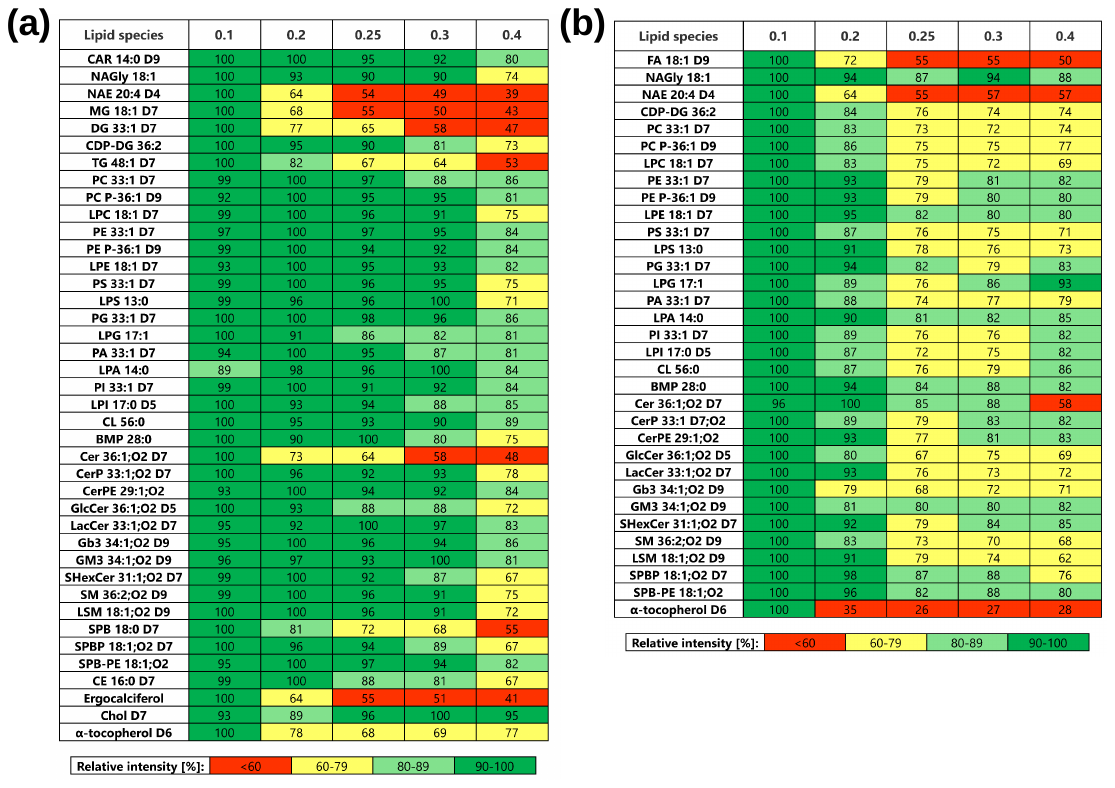

**Figure S2**: Optimization of makeup flow rate based on response of individual lipid species in **(a)** positive and **(b)** negative ion modes.
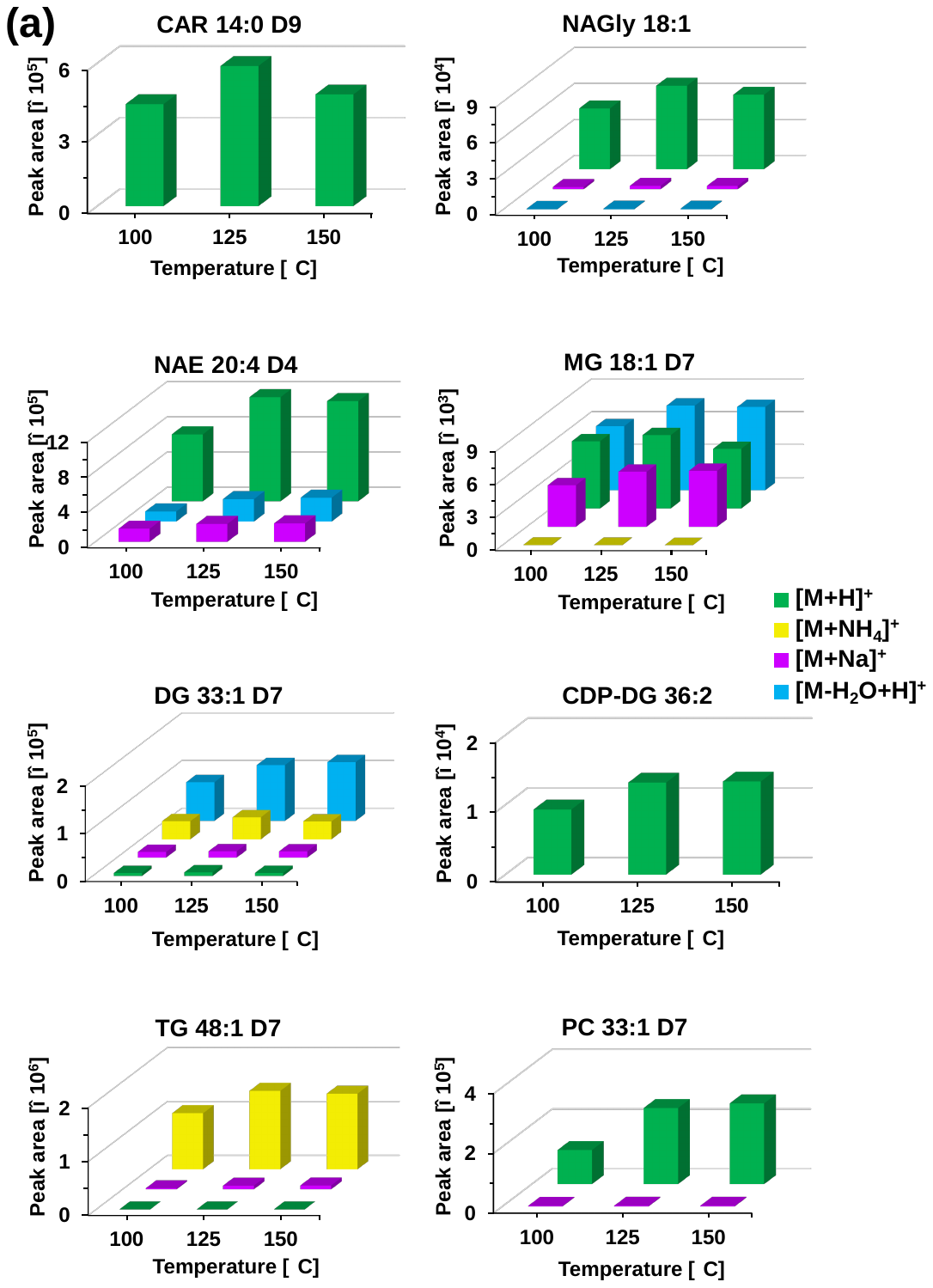

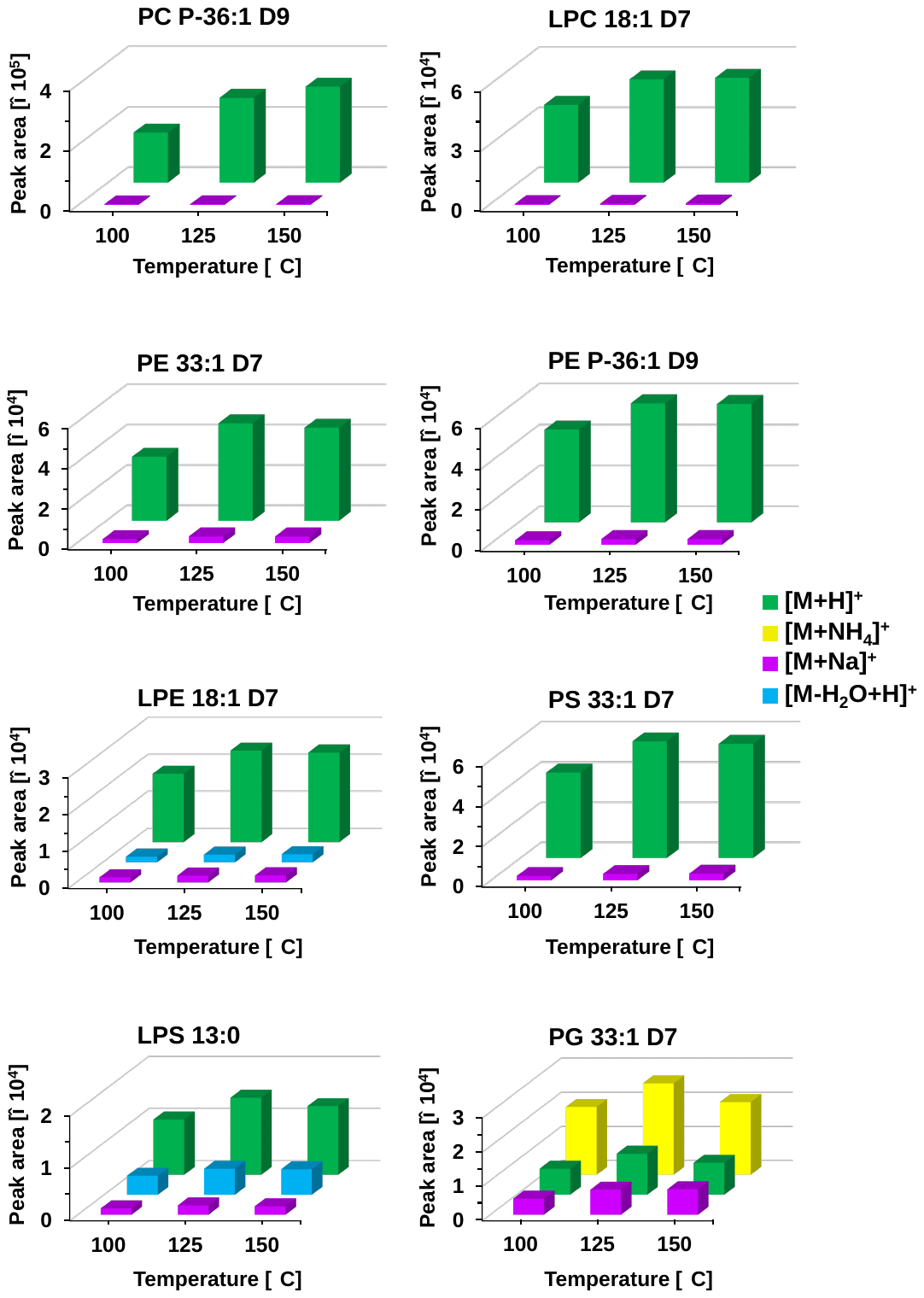

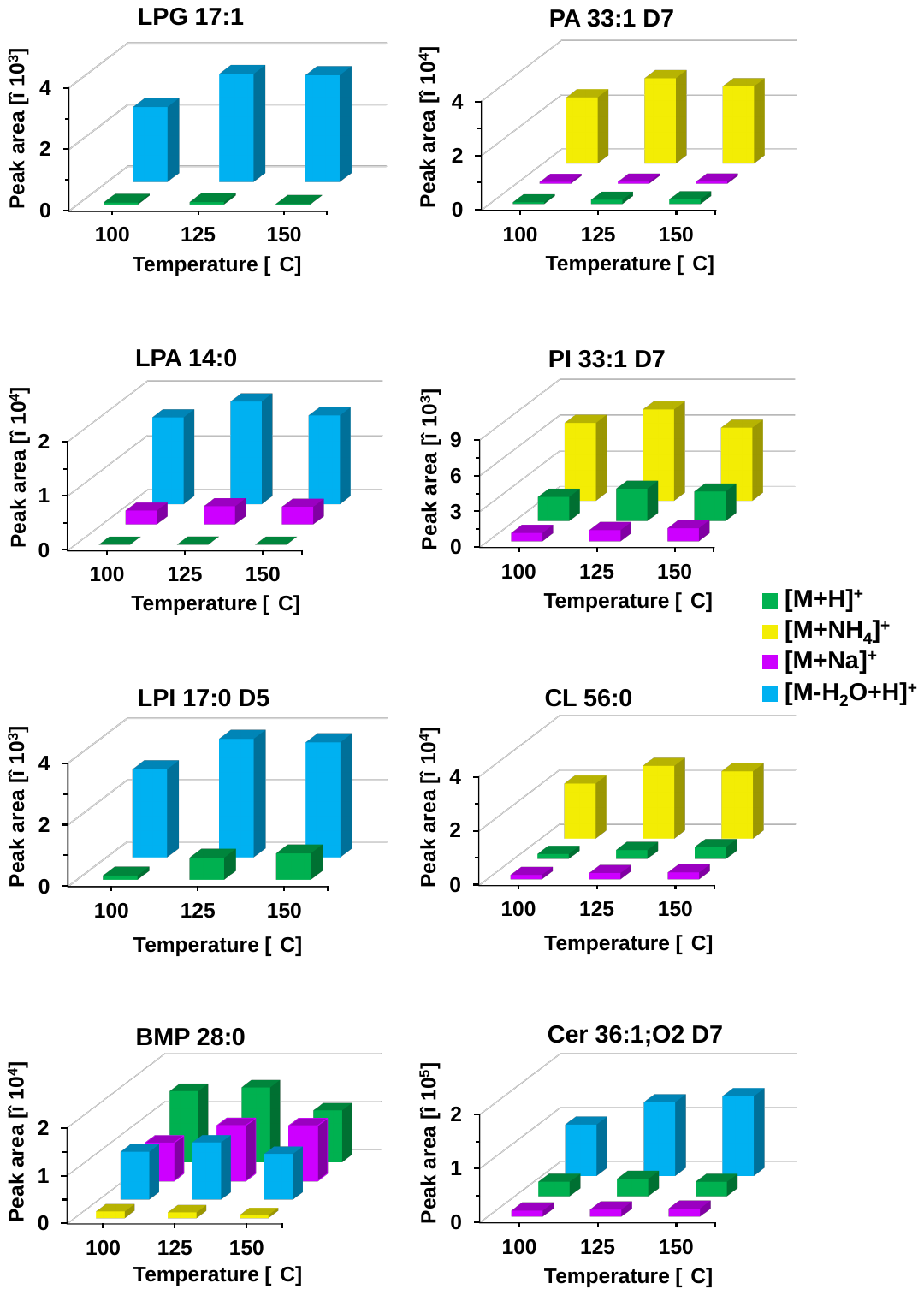

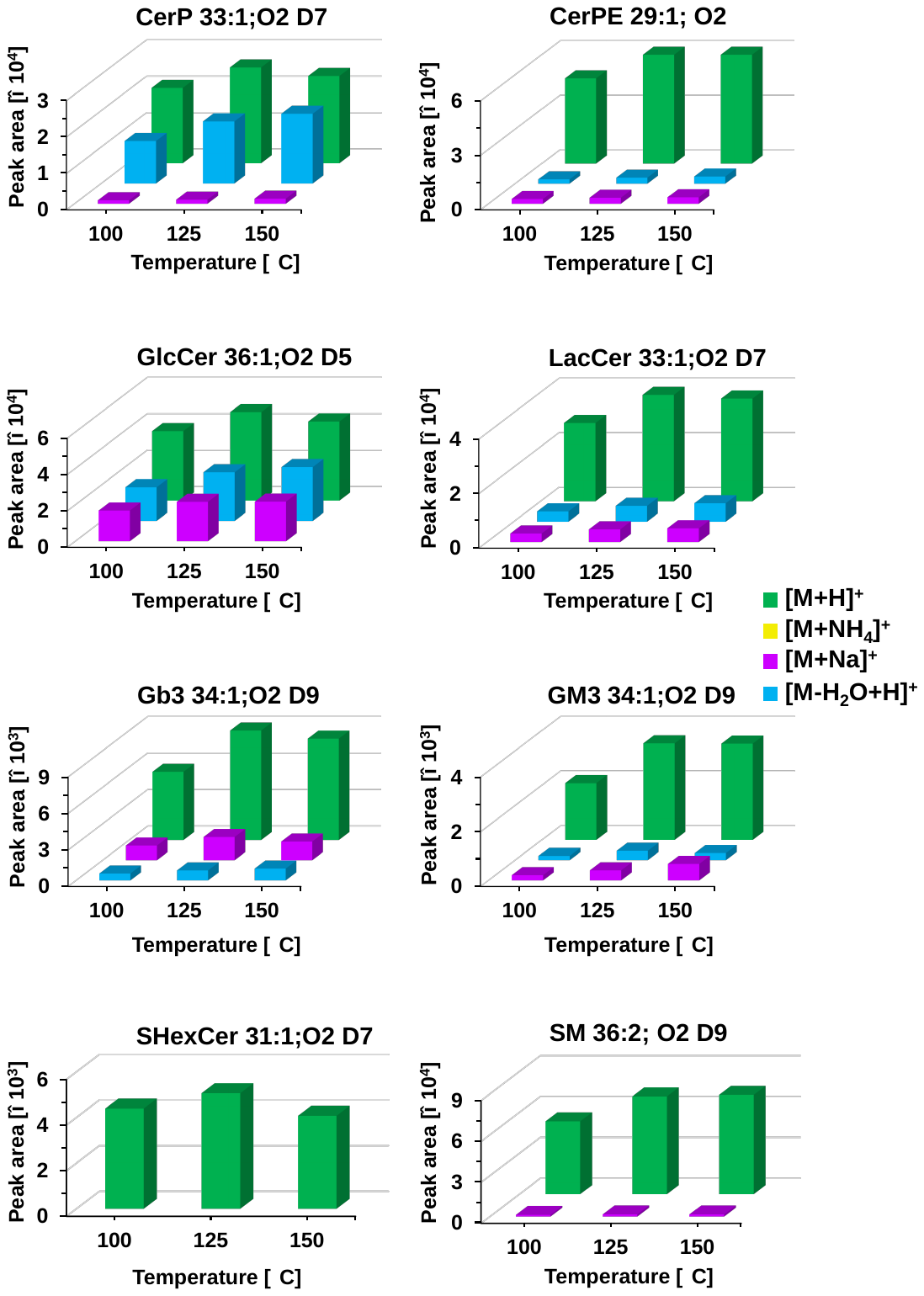

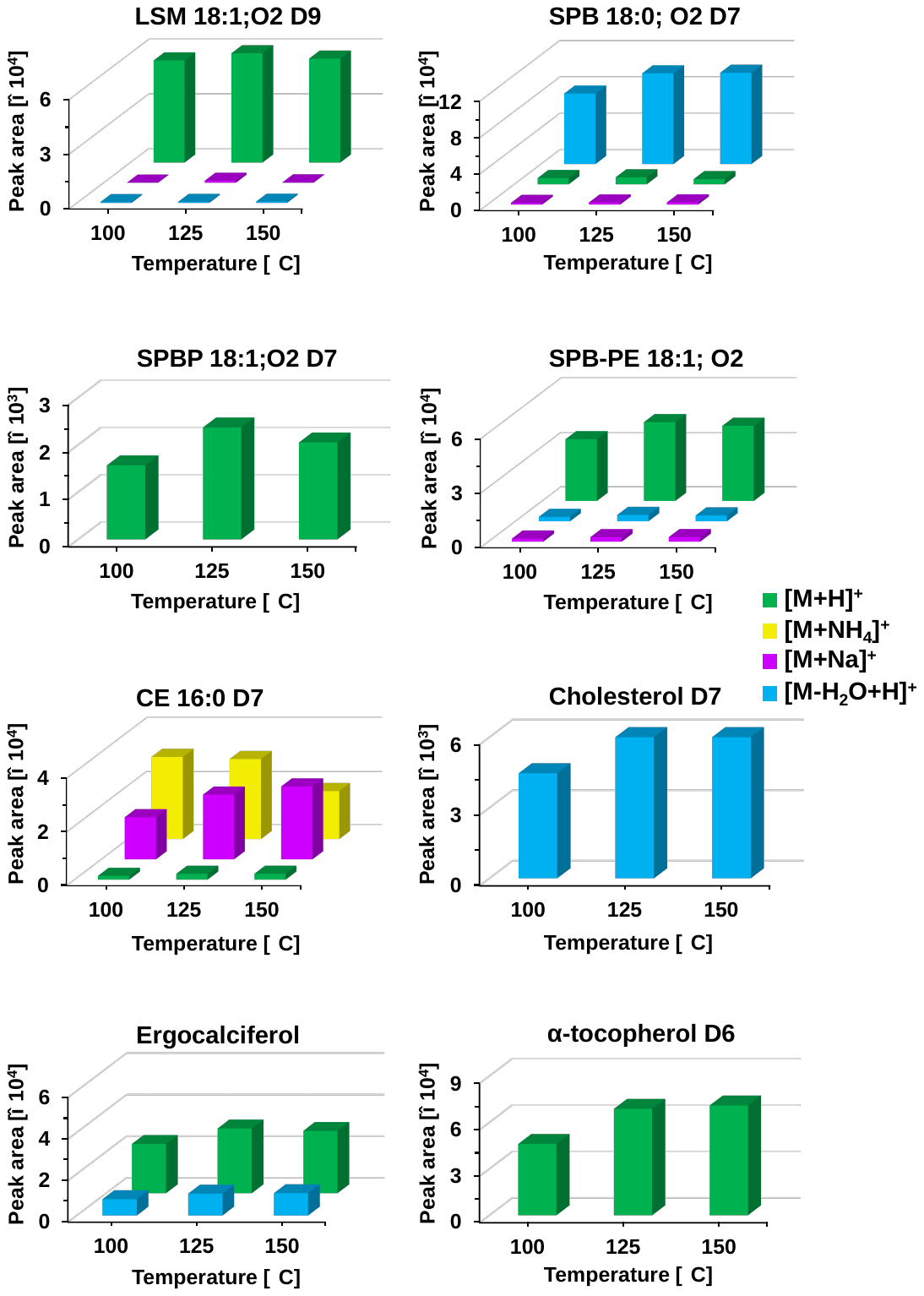

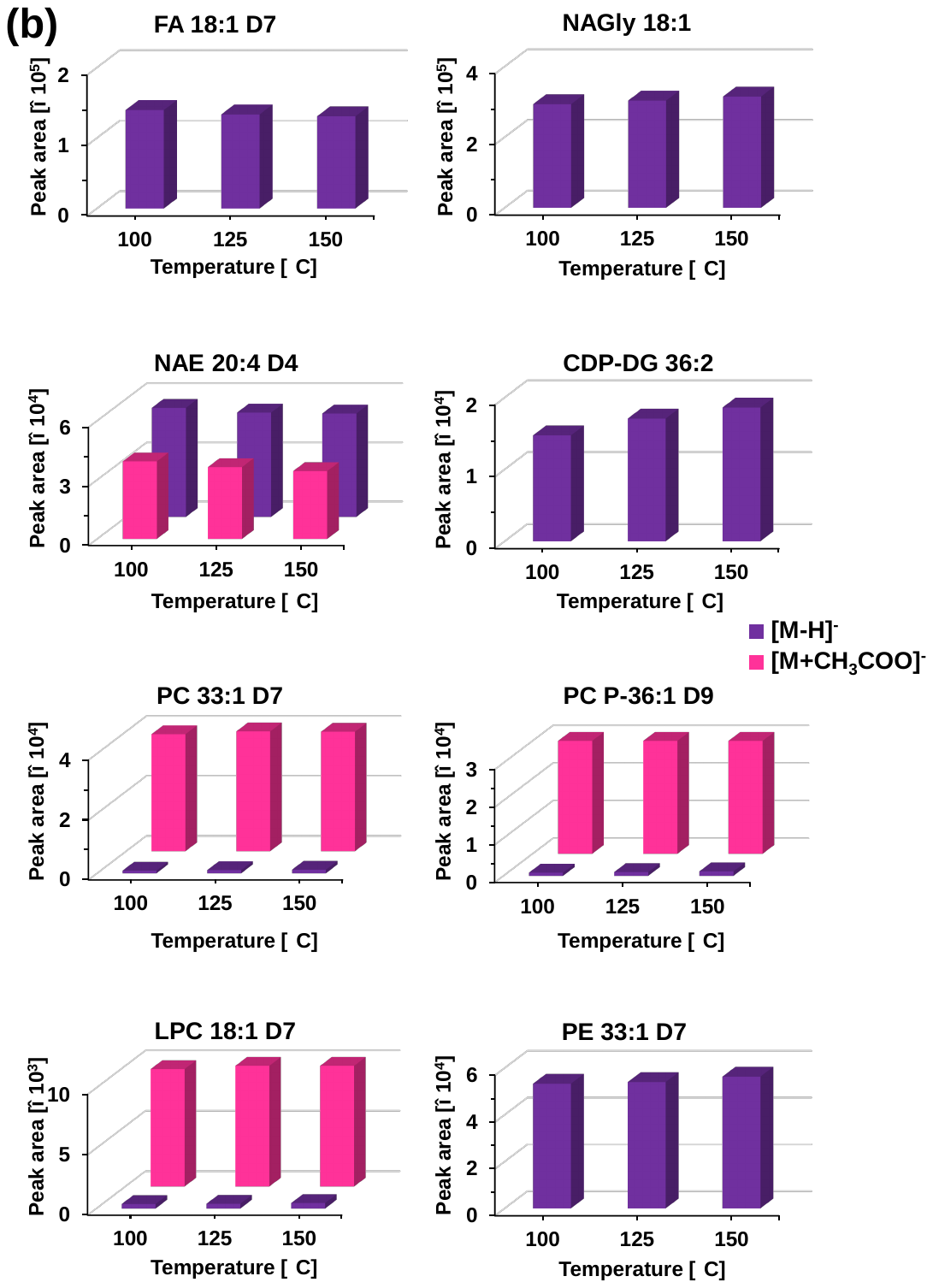

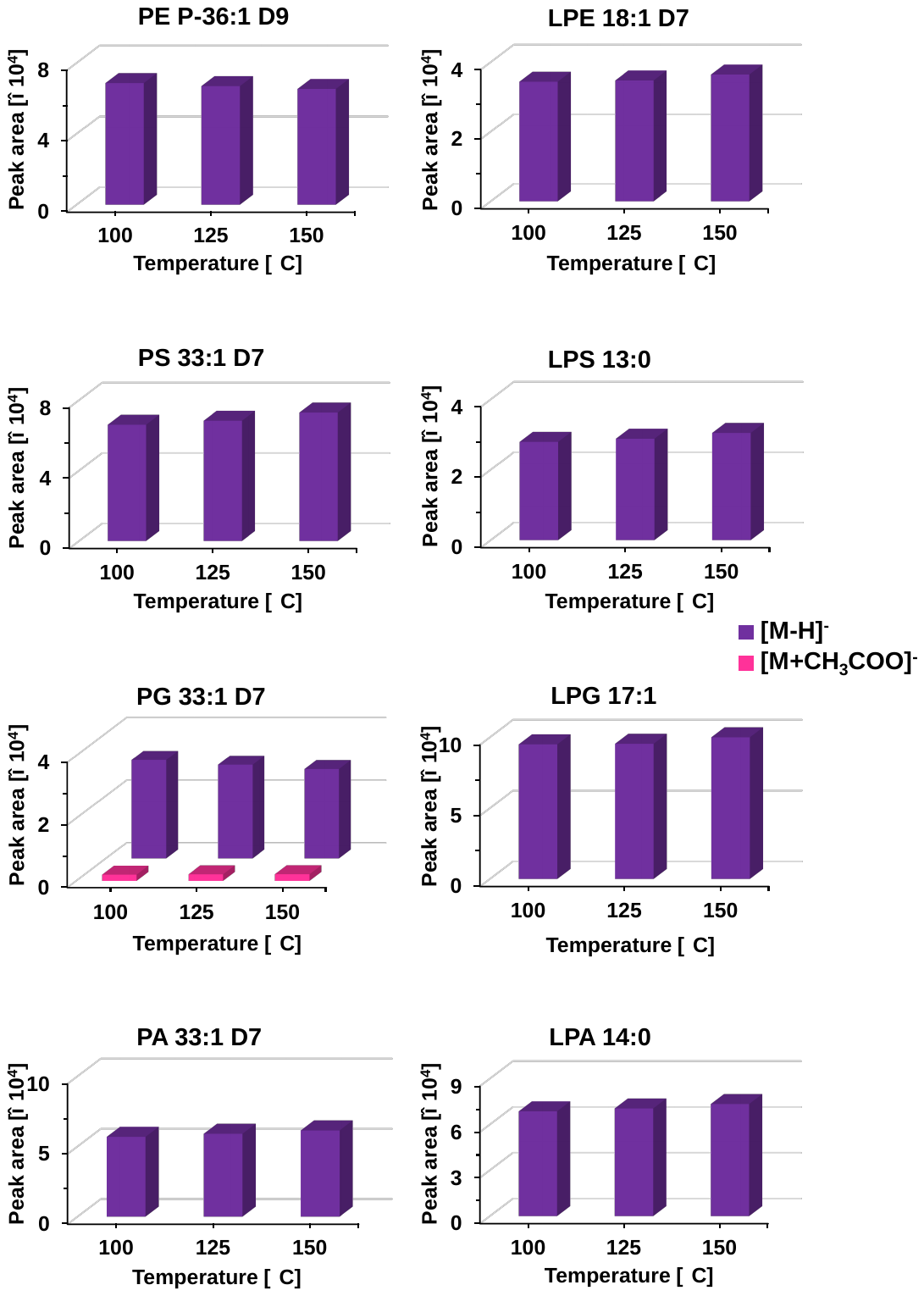

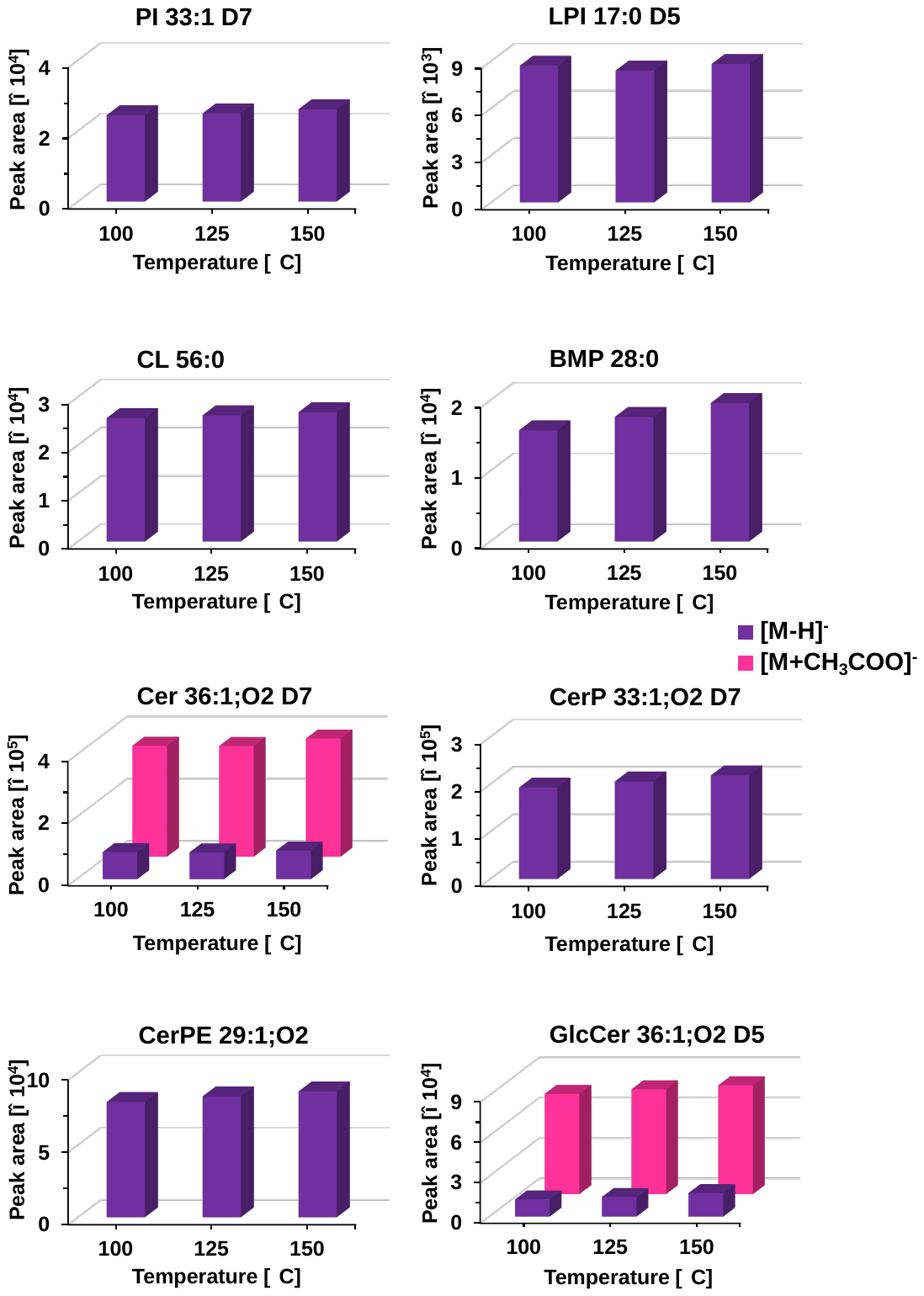

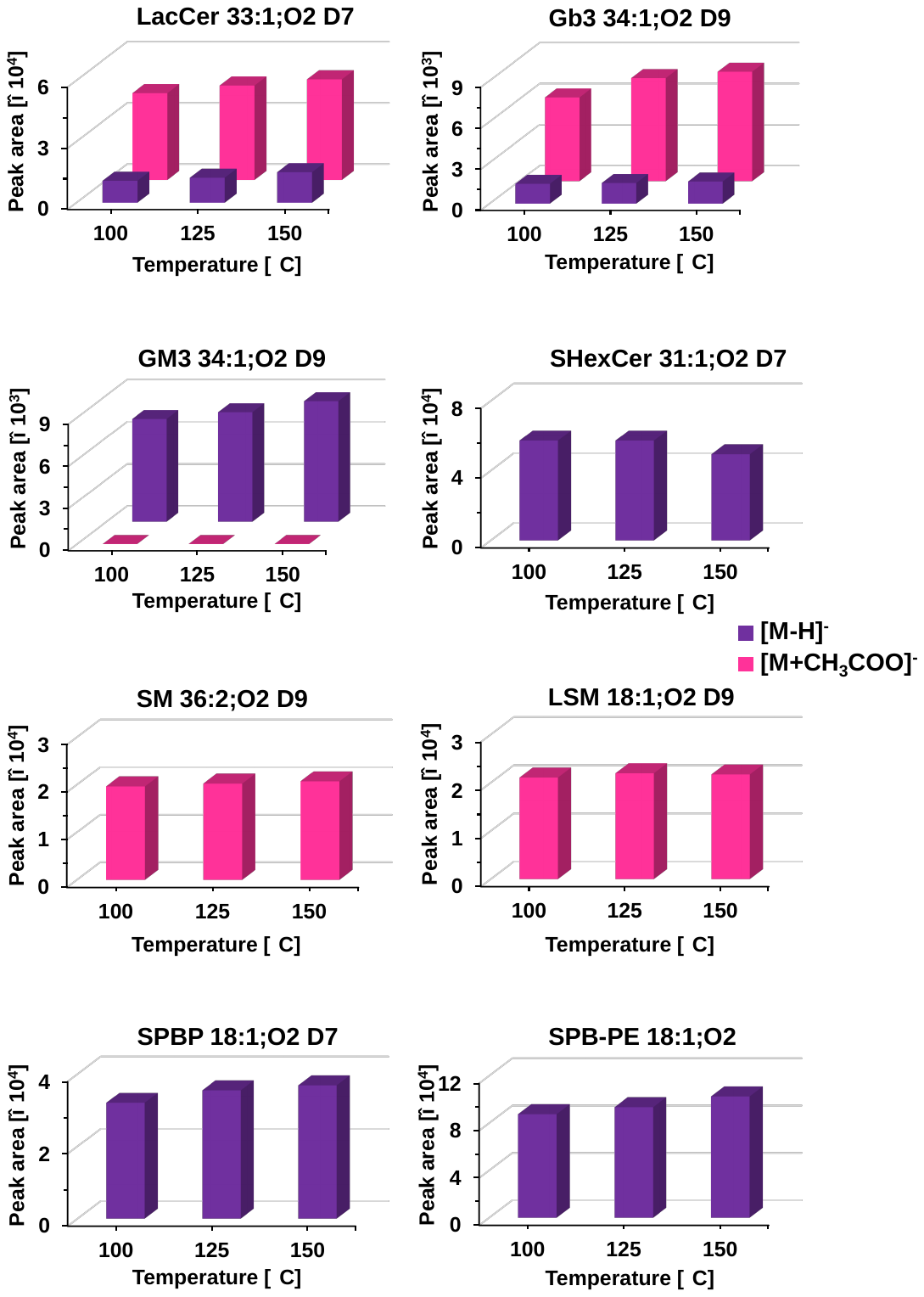

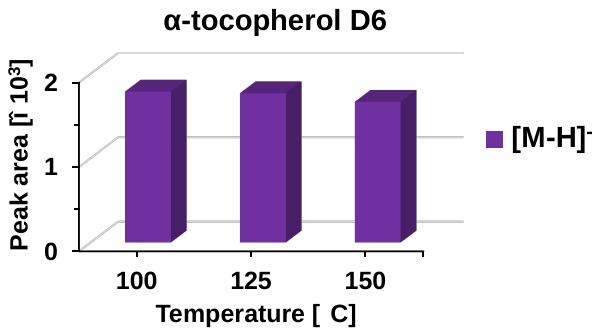

**Figure S3**: Optimization of source temperature in **(a)** positive and **(b)** negative ion modes. Annotation: Green [M+H]^+^, yellow [M+NH_4_]^+^, light purple [M+Na]^+^, blue [M-H_2_O+H]^+^, dark purple [M-H]^-^, and pink [M+CH_3_COO]^-^.

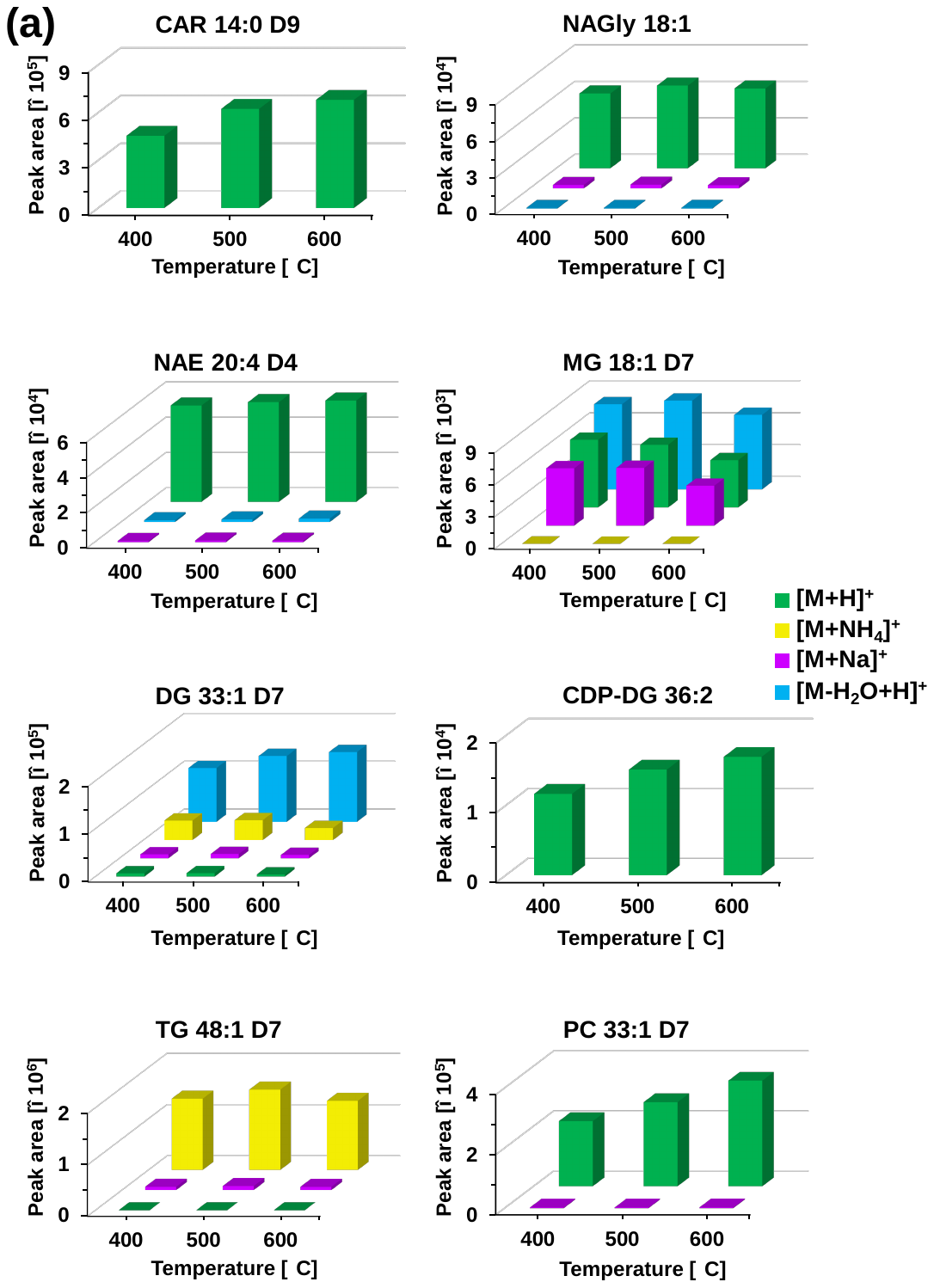

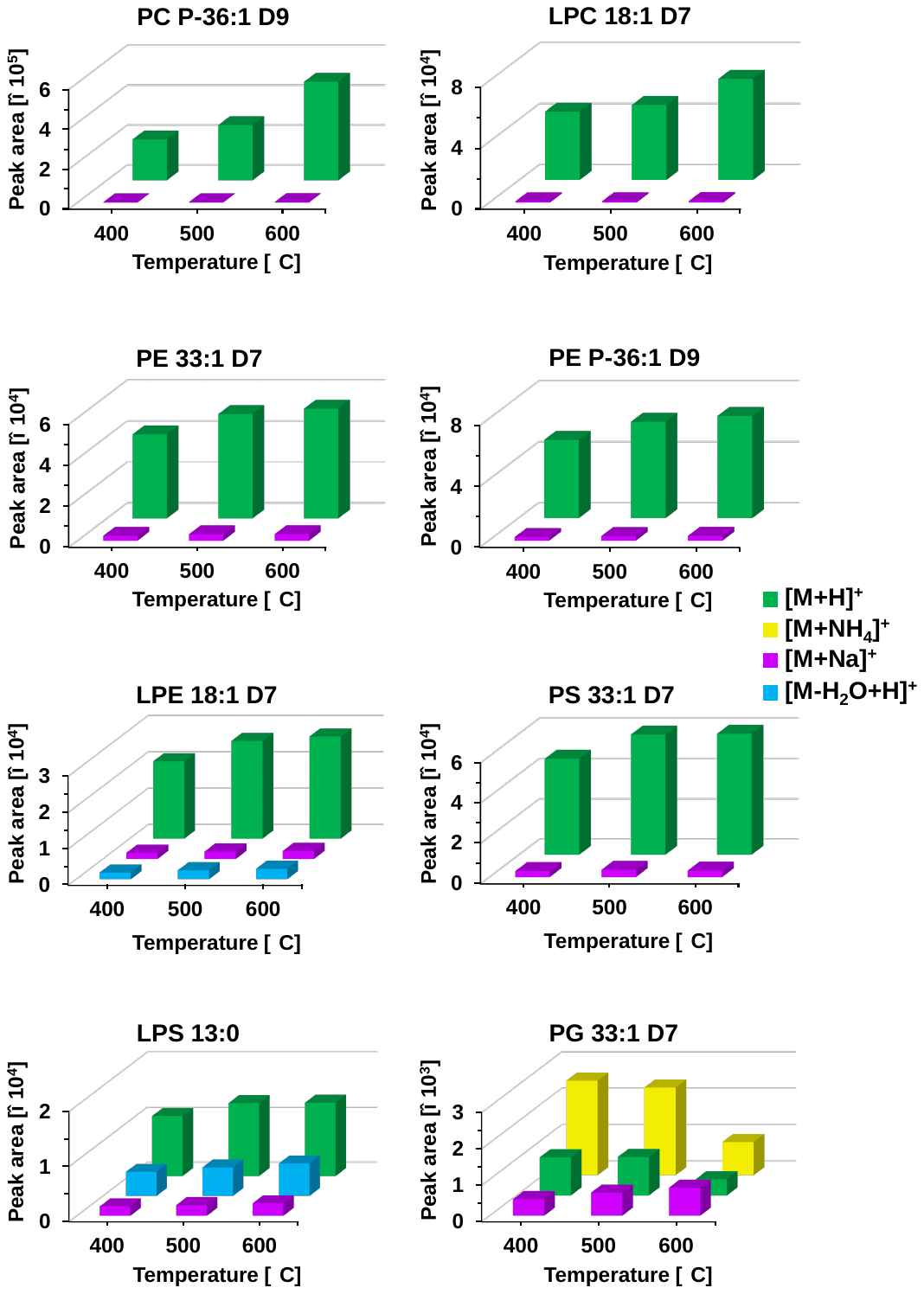

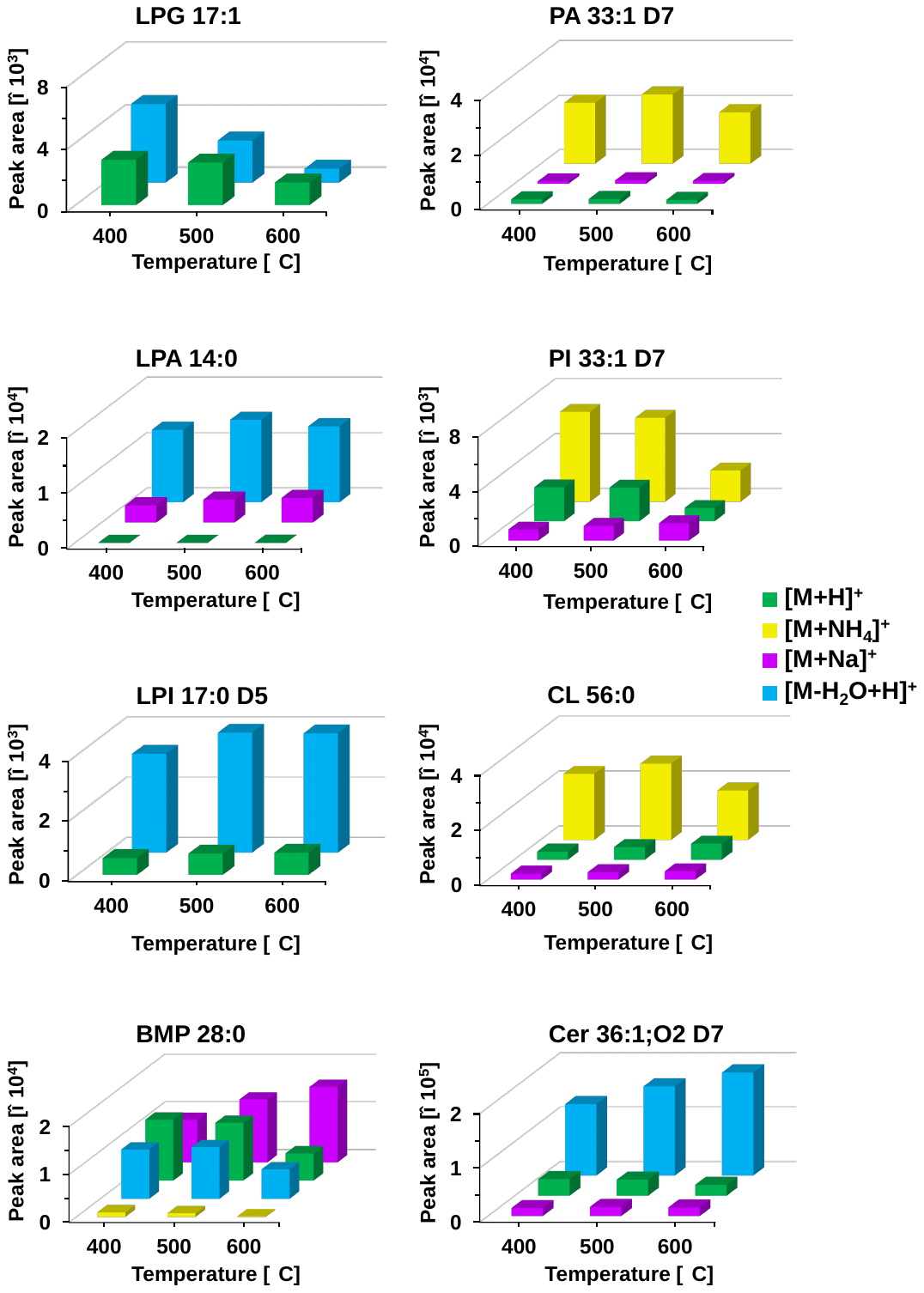

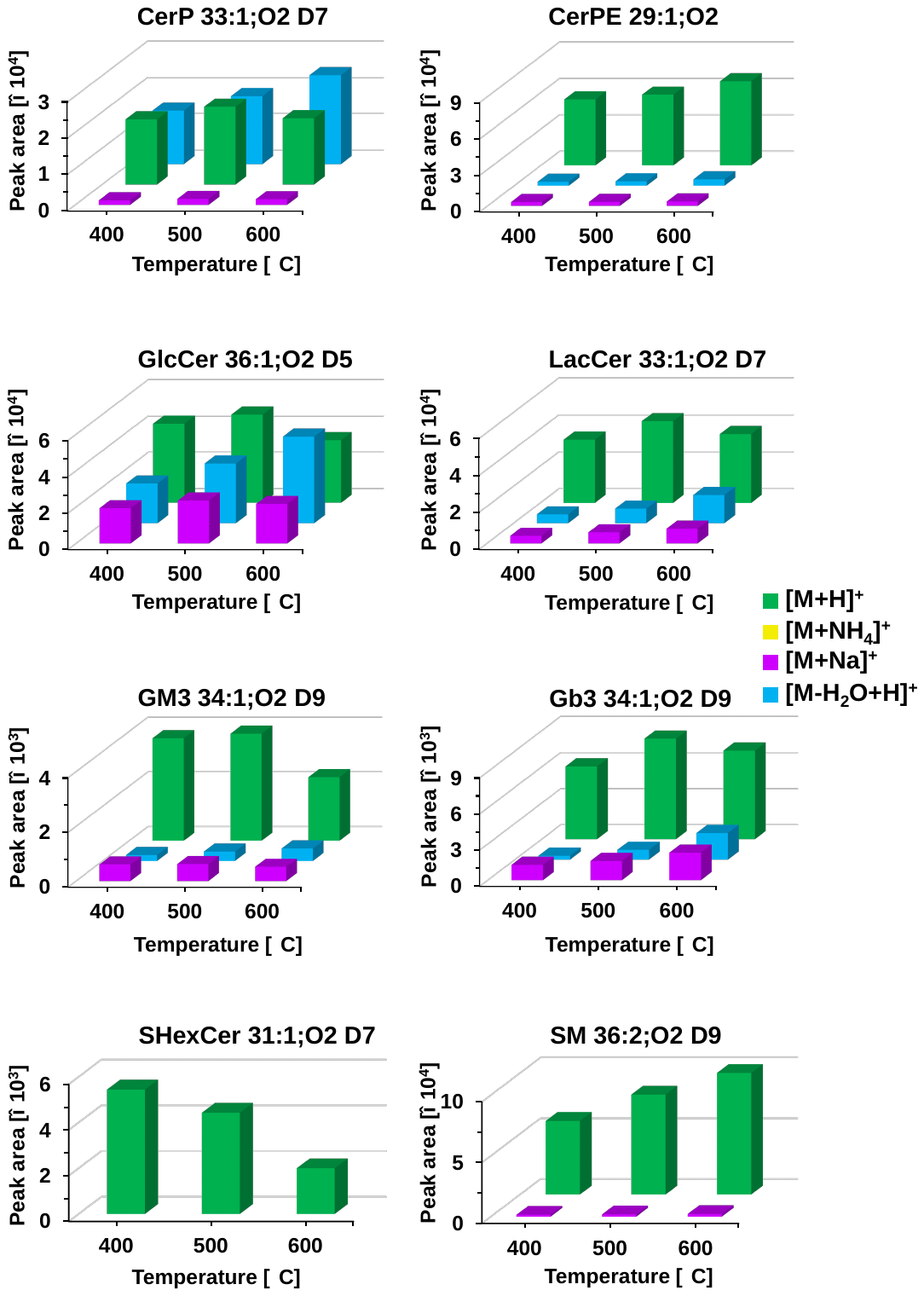

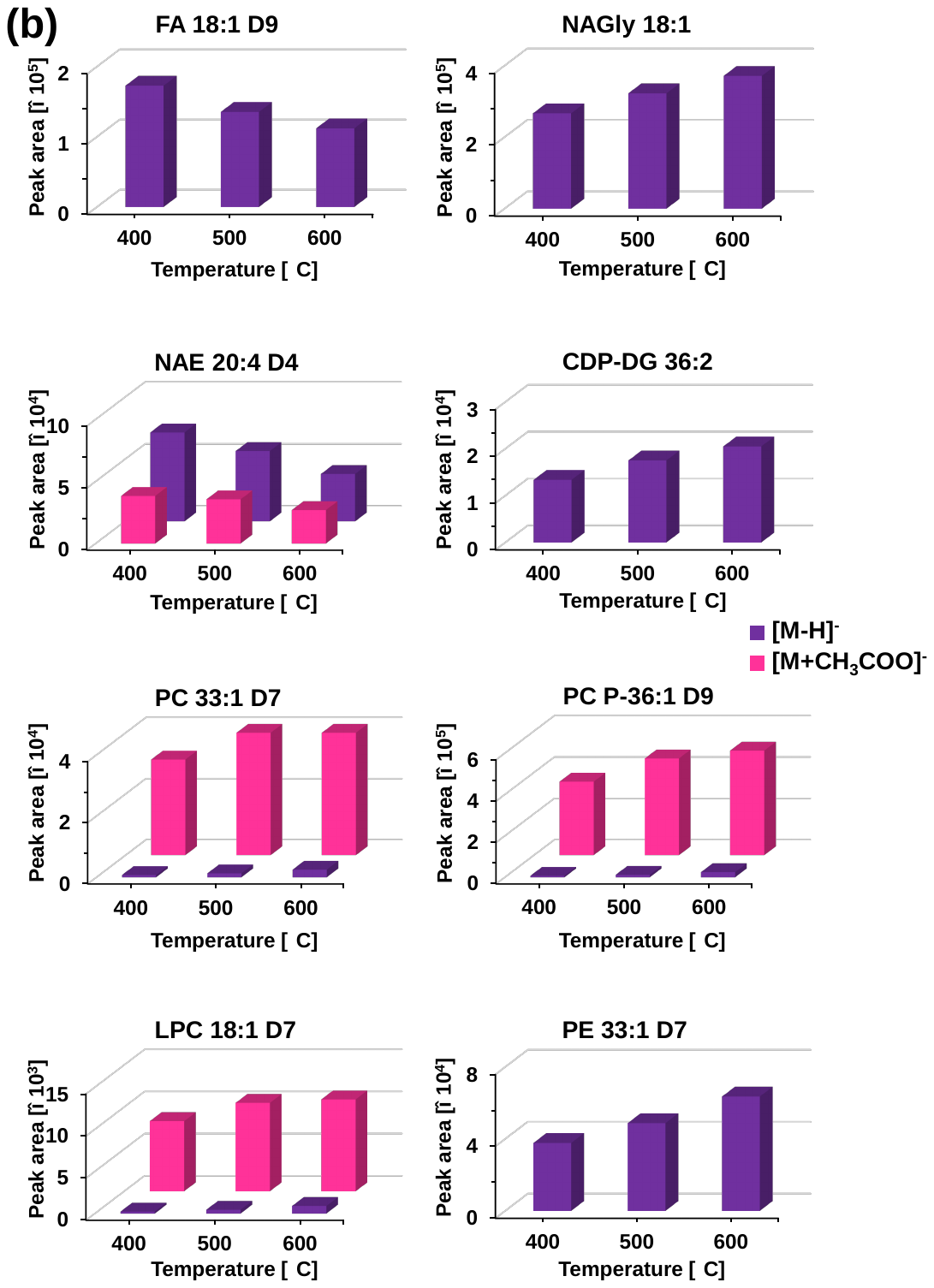

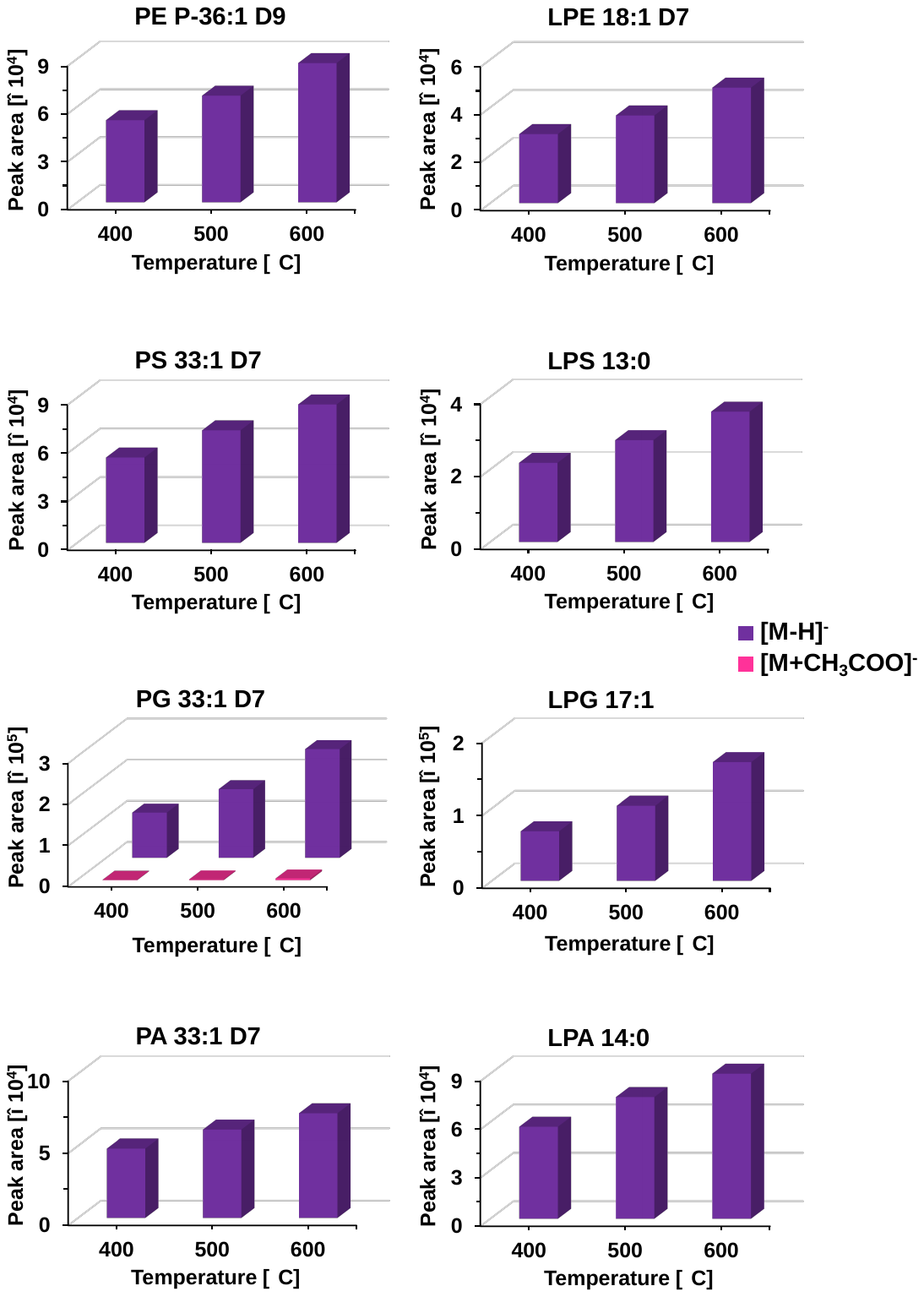

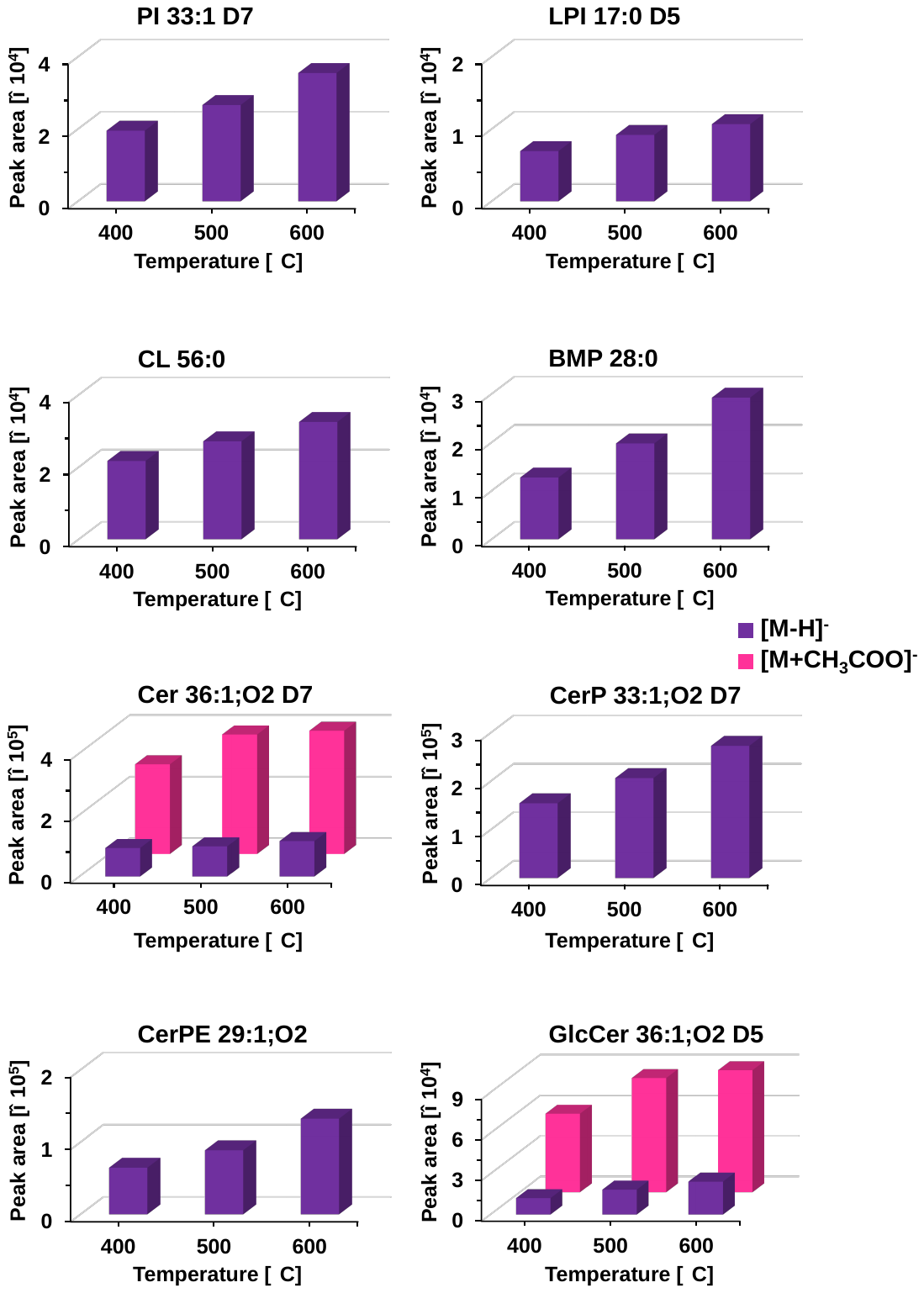

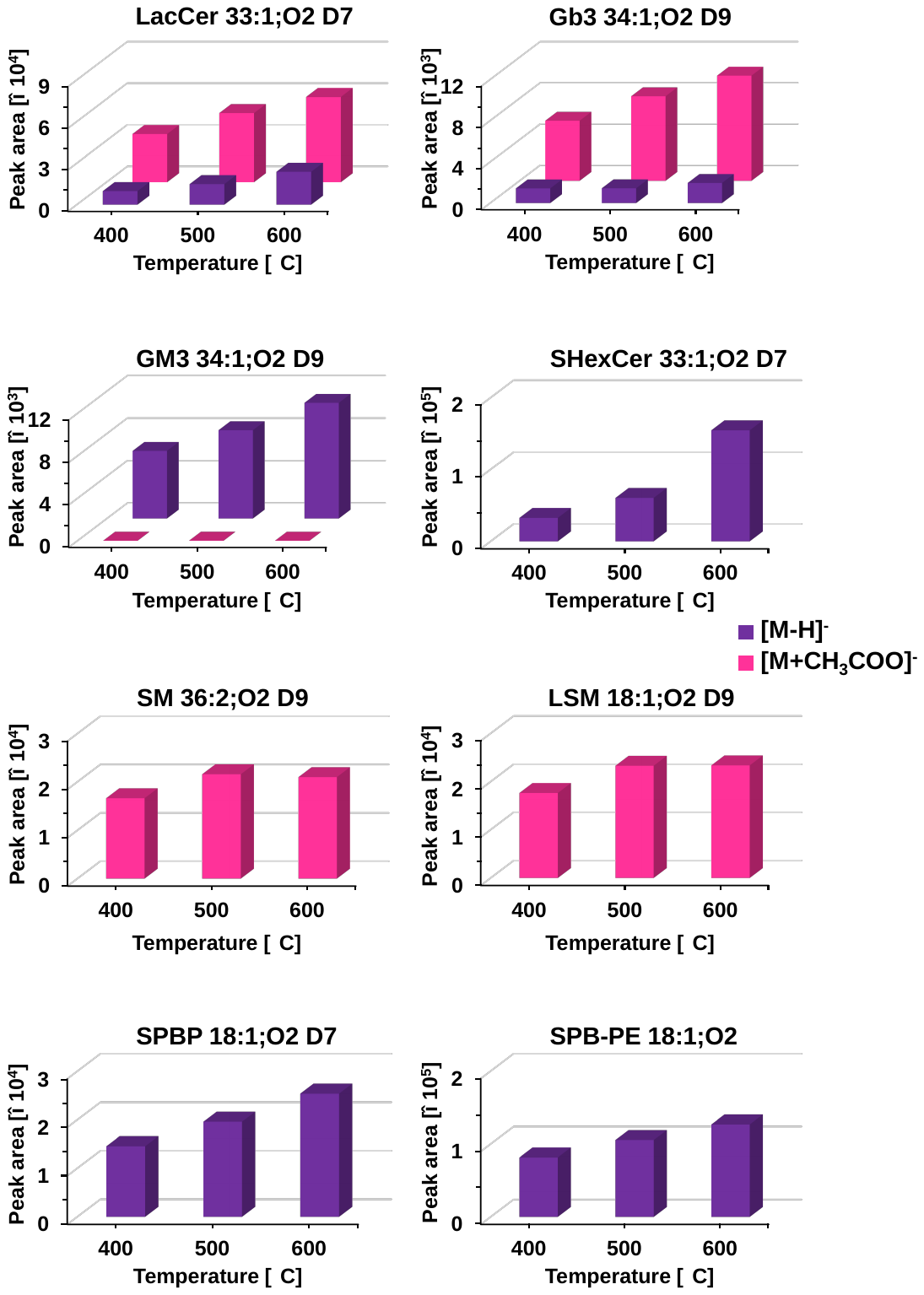

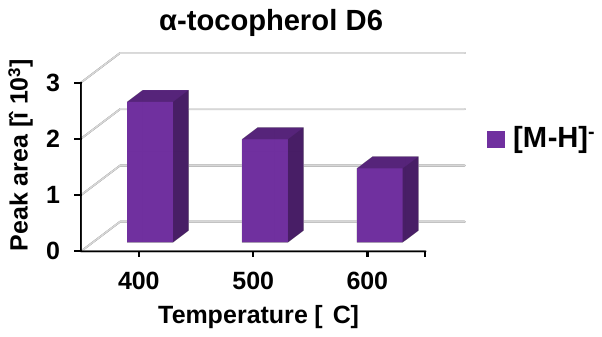

**Figure S4**: Optimization of desolvation temperature in **(a)** positive and **(b)** negative ion modes. Annotation: Green [M+H]^+^, yellow [M+NH_4_]^+^, light purple [M+Na]^+^, blue [M-H_2_O+H]^+^, dark purple [M-H]^-^, and pink [M+CH_3_COO]^-^.

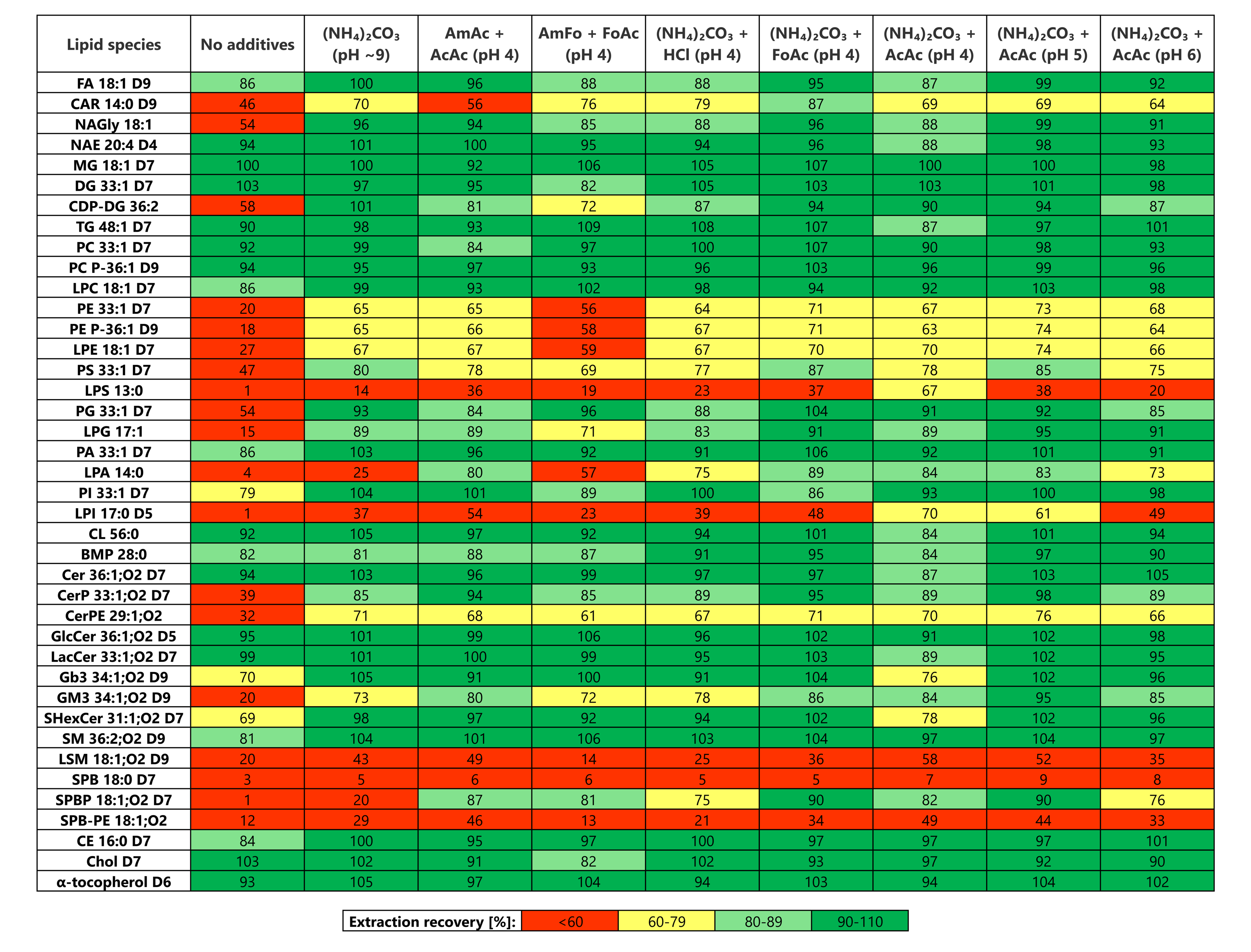

**Figure S5**: Optimization of the additive composition in the aqueous phase for double Folch extraction evaluated based on extraction recoveries of individual lipid species. Abbreviations: ammonium acetate (AmAc), acetic acid (AcAc), ammonium formate (AmFo), and formic acid (FoAc).

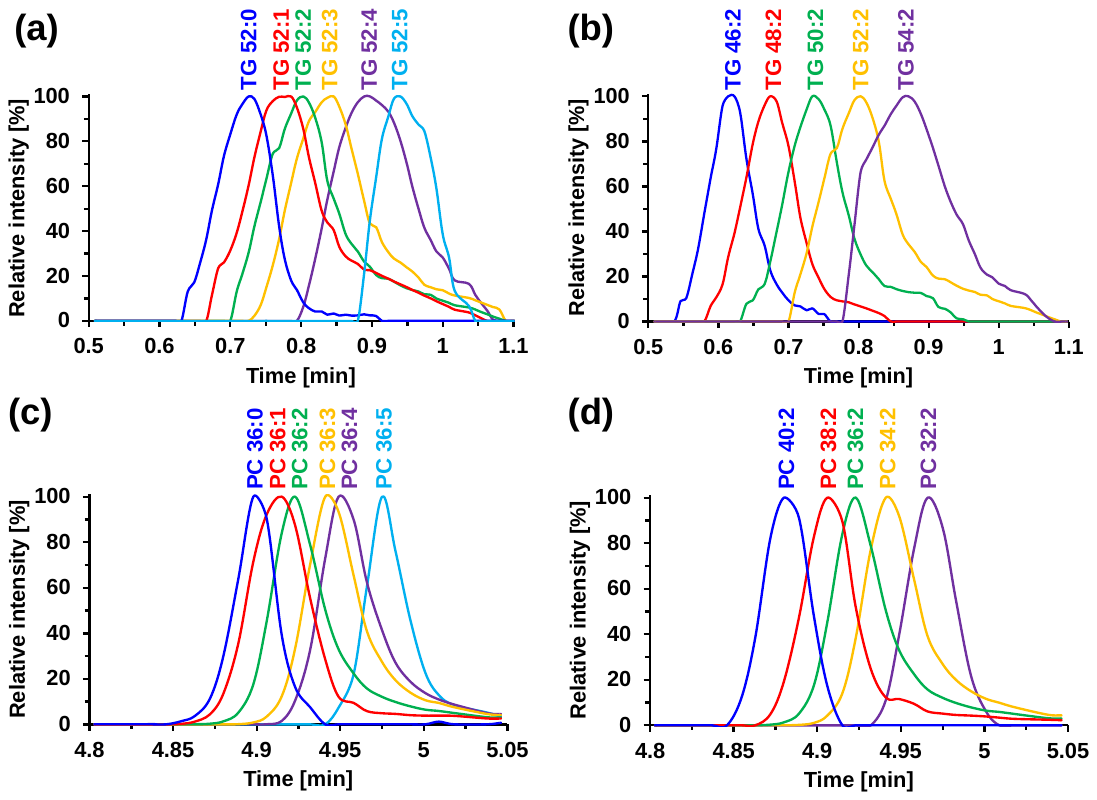

**Figure S6**: Dependence of retention times on **(a)** the number of double bonds in TG, **(b)** the carbon number in TG, **(c)** the number of double bonds in PC, and **(d)** the carbon number in PC.

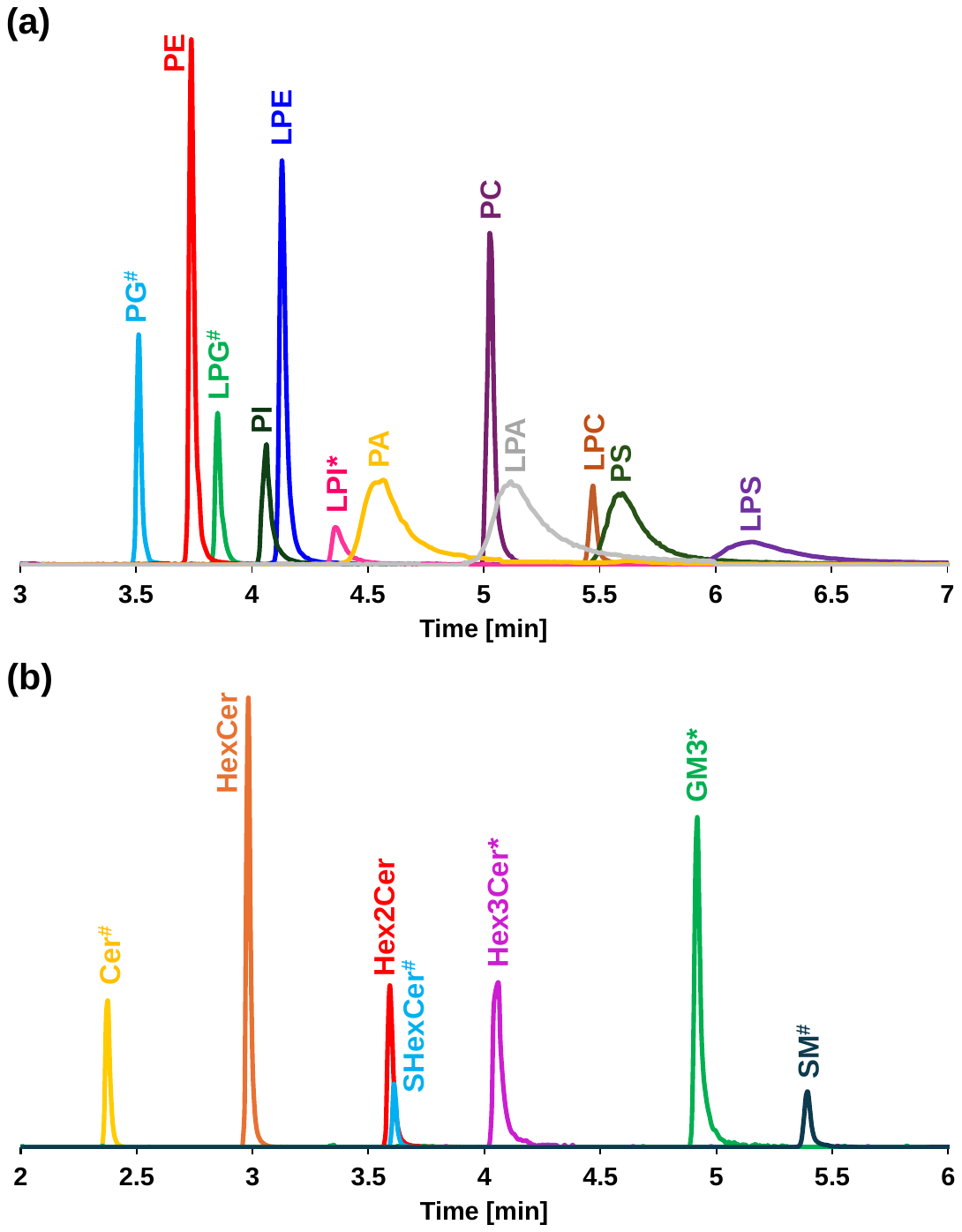

**Figure S7**: Separation of **(a)** phospholipids and **(b)** sphingolipids in negative ion mode. Signal of individual lipid species was modified for visualization as follows: ^#^ indicates signal decreased 10-fold and * indicates signal increased 10-fold.

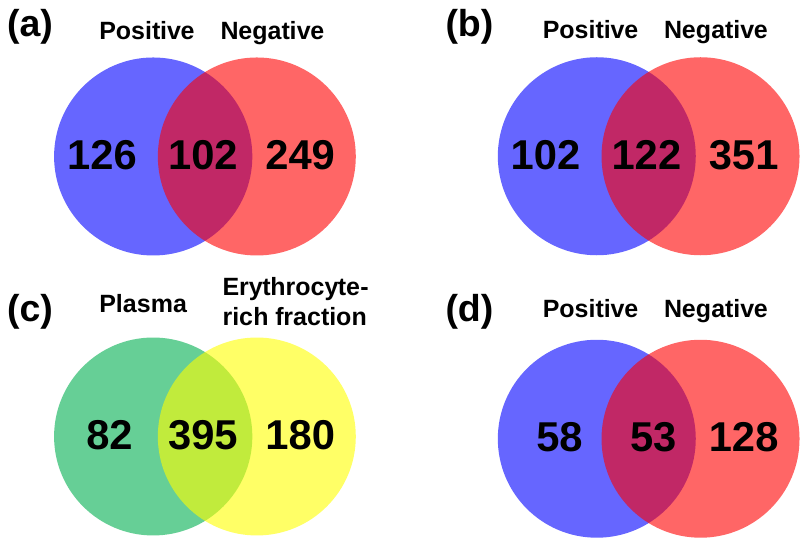

**Figure S8:** Venn diagrams comparing the numbers of lipid species: **(a)** identified in positive and negative ion modes in the plasma matrix, **(b)** identified in positive and negative ion modes in the erythrocyte-rich fraction, **(c)** identified in the plasma and erythrocyte-rich matrices in both polarities, and **(d)** quantified in positive and negative ion modes in the plasma matrix.
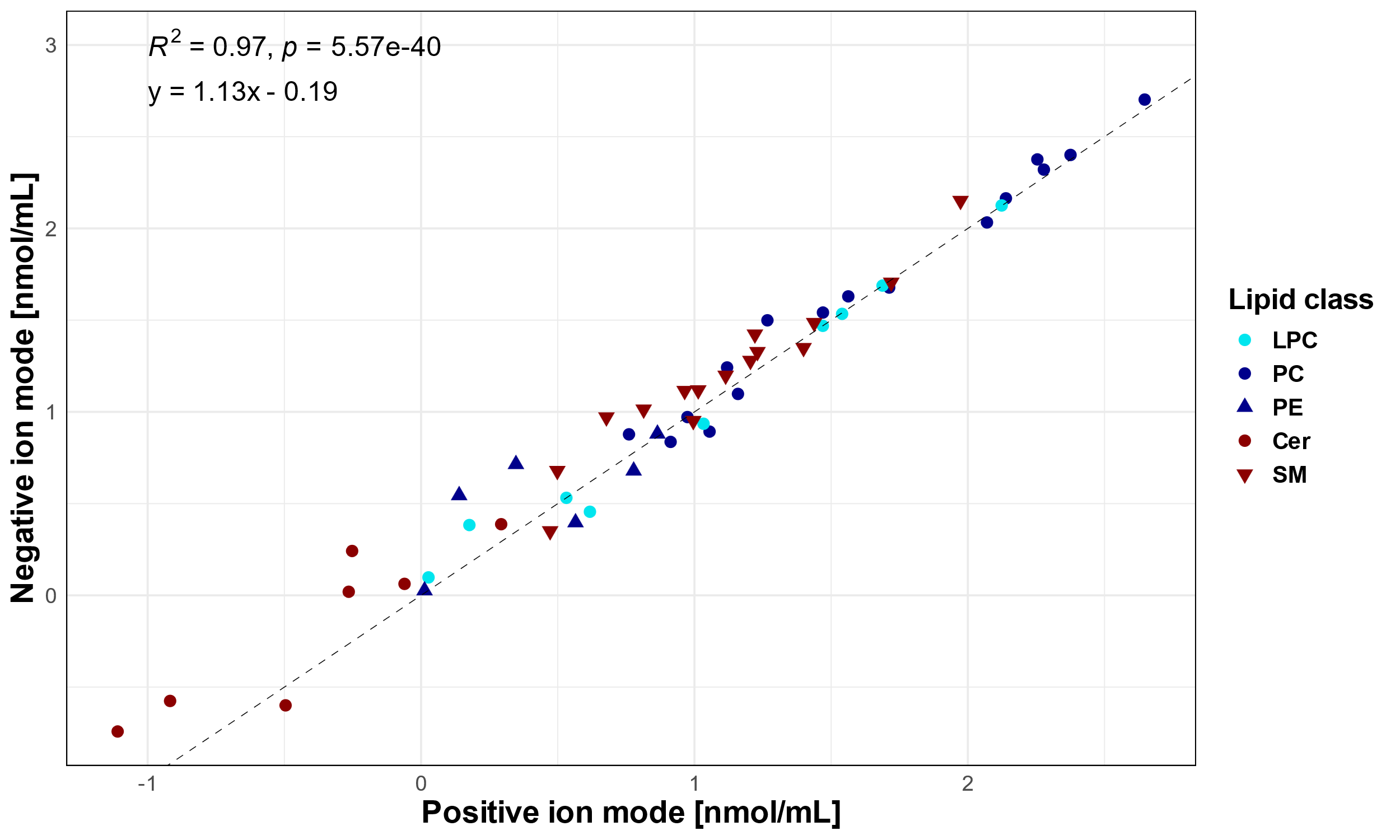

**Figure S9**: Correlation graph comparing the determined concentrations in positive and negative ion modes using the current method.

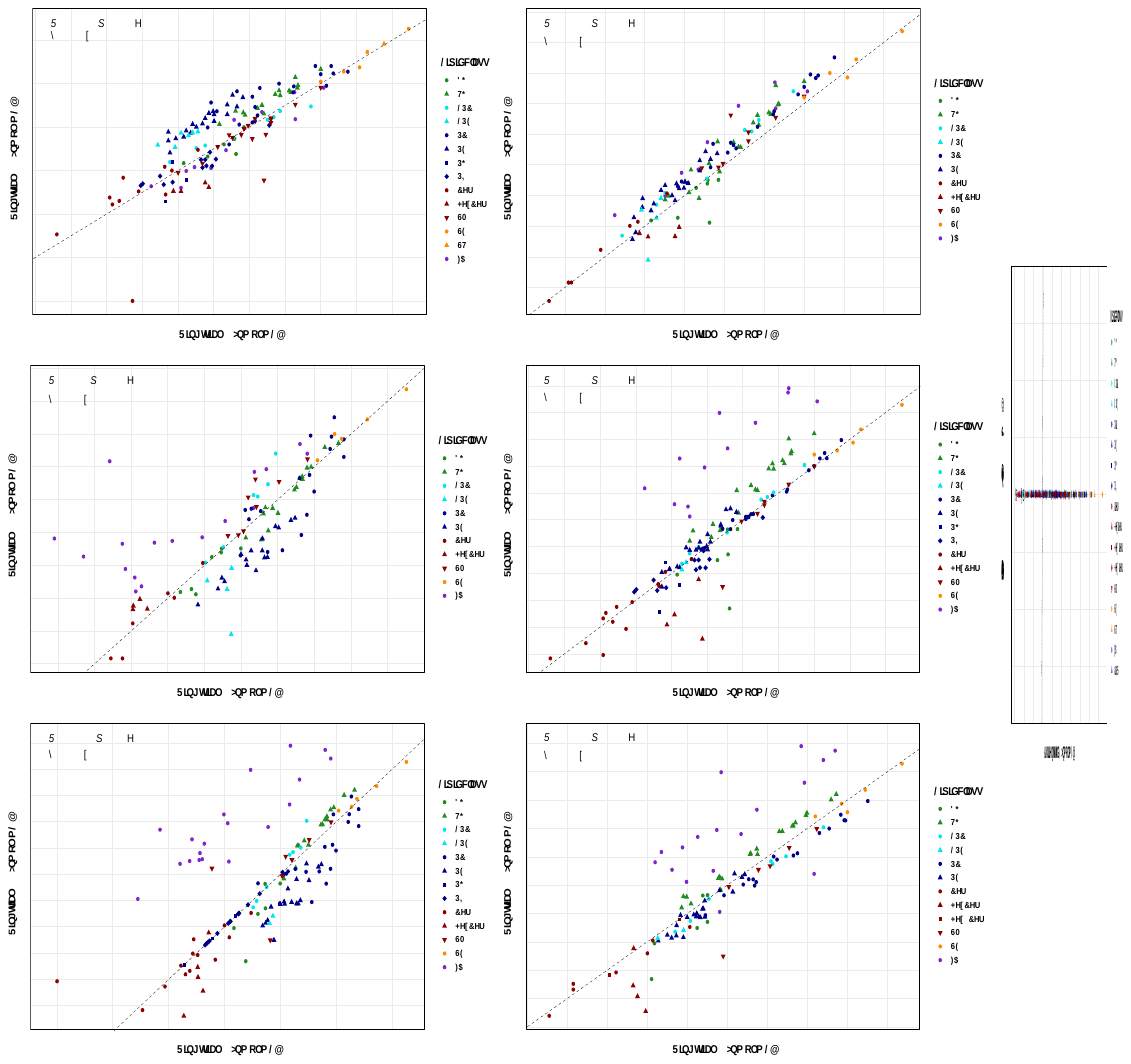

**Figure S10:** Correlation graphs for NIST SRM 1950 human plasma comparing results from individual ring trials: Bowden et al. (Ring trial 1) [4], Quehenberger et al. (Ring trial 2) [5], Ghorasaini et al. (Ring trial 3) [6], and Mandal et al. (Ring trial 4) [7].
