## Supplementary material for "Comprehensive Lipidomic Characterization of Human Blood by High-Throughput UHPSFC/MS using Bioinert Column and Modified MTBE Extraction": Reporting Checklist

### Contents of Report

Created by https://lipidomicstandards.org, version v2.5.0

**Separation Workflow** 2
- Overall study design 2
- Lipid extraction 2
- Analytical platform 2
- Quality control 3
- Method qualification and validation 3
- Reporting 3

**Sample Descriptions** 3
- Pooled plasma / Human / Plasma 3
- Pooled erythrocyte-rich fraction / Human / Cells 3

**Lipid Class Descriptions** 3
- 1) TG[M+NH4]+ / Lipid identification 3
- 1) TG[M+NH4]+ / Lipid quantification 4
- 2) PC[M+H]+ / Lipid identification 4
- 2) PC[M+H]+ / Lipid quantification 4
- 3) DG[M-H2O+H]+ / Lipid identification 5
- 3) DG[M-H2O+H]+ / Lipid quantification 5
- 4) PC[M+CH3COO]- / Lipid identification 5
- 4) PC[M+CH3COO]- / Lipid quantification 6
- 5) MG[M+H]+ / Lipid identification 6
- 5) MG[M+H]+ / Lipid quantification 6
- 6) CAR[M+H]+ / Lipid identification 7
- 6) CAR[M+H]+ / Lipid quantification 7
- 7) NAE[M-H]- / Lipid identification 7
- 7) NAE[M-H]- / Lipid quantification 8
- 8) PC P[M+H]+ / Lipid identification 8
- 8) PC P[M+H]+ / Lipid quantification 8
- 9) PC P[M+CH3COO]- / Lipid identification 8
- 9) PC P[M+CH3COO]- / Lipid quantification 9
- 10) FA[M-H]- / Lipid identification 9
- 10) FA[M-H]- / Lipid quantification 9
- 11) LPC[M+H]+ / Lipid identification 10
- 11) LPC[M+H]+ / Lipid quantification 10
- 12) LPC[M+CH3COO]- / Lipid identification 10
- 12) LPC[M+CH3COO]- / Lipid quantification 11
- 13) PE[M+H]+ / Lipid identification 11
- 13) PE[M+H]+ / Lipid quantification 11
- 14) PE[M-H]- / Lipid identification 12
- 14) PE[M-H]- / Lipid quantification 12
- 15) PE P[M+H]+ / Lipid identification 12
- 15) PE P[M+H]+ / Lipid quantification 13
- 16) PE P[M-H]- / Lipid identification 13
- 16) PE P[M-H]- / Lipid quantification 13
- 17) LPE[M+H]+ / Lipid identification 14
- 17) LPE[M+H]+ / Lipid quantification 14
- 18) LPE[M-H]- / Lipid identification 14
- 18) LPE[M-H]- / Lipid quantification 15
- 19) PS[M-H]- / Lipid identification 15
- 19) PS[M-H]- / Lipid quantification 15
- 20) LPC O-/P-[M+H]+ / Lipid identification 15
- 20) LPC O-/P-[M+H]+ / Lipid quantification 16
- 21) LPE P-/O-[M-H]- / Lipid identification 16
- 21) LPE P-/O-[M-H]- / Lipid quantification 16
- 22) PG[M-H]- / Lipid identification 17
- 22) PG[M-H]- / Lipid quantification 17
- 23) LPG[M-H]- / Lipid identification 17
- 23) LPG[M-H]- / Lipid quantification 18
- 24) PI[M-H]- / Lipid identification 18
- 24) PI[M-H]- / Lipid quantification 18
- 25) LPI[M-H]- / Lipid identification 19
- 25) LPI[M-H]- / Lipid quantification 19
- 26) PA[M-H]- / Lipid identification 19
- 26) PA[M-H]- / Lipid quantification 20
- 27) LPS[M-H]- / Lipid identification 20
- 27) LPS[M-H]- / Lipid quantification 20
- 28) CL[M-H]- / Lipid identification 20

### Separation Workflow

#### Overall study design

| | | | |
|---|---|---|---|
| Title of the study | UHPSFC/MS and bioinert column | | |
| Document creation date | 12/15/2025 | Principal investigator | Petra Peroutková |
| Institution | University of Pardubice | Corresponding Email | |
| Is the workflow targeted or untargeted? | Untargeted | Clinical | Yes |

#### Lipid extraction

| | | | |
|---|---|---|---|
| Extraction method | 2-phase system | pH adjustment | Acetic acid |
| 2-phase system | MTBE | Were internal standards added prior extraction? | Yes |

#### Analytical platform

| | | | |
|---|---|---|---|
| Ionizaton additives | Ammonium acetate | Number of separation dimensions | One dimension |
| Separation type 1 | SFC | Separation mode 1 (liquid) | HILIC |
| Detector | Mass spectrometer | MS type | QTOF |
| MS vendor | Waters | Ion source | ESI |

| | | | |
|---|---|---|---|
| MS Level | $MS^1$ | Mass resolution for detected ion at $MS^1$ | High resolution |
| Resolution at m/z 200 at $MS^1$ | 25000 | Mass accuracy in ppm at $MS^1$ | 5 |
| Recording mode of raw data at $MS^1$ | Profile mode | Was/Were additional dimension/techniques used | No |

#### Quality control

| | | | |
|---|---|---|---|
| Blanks | Yes | Type of Blanks | Extraction blank, Solvent blank, Internal standard blank |
| Quality control | Yes | Type of QC sample | Sample pool |

#### Method qualification and validation

| | | | |
|---|---|---|---|
| Method validation | Yes | Lipid recovery | Yes |
| Dynamic quantification range | Yes | Limit of quantitation (LOQ)/Limit of detection (LOD) | Yes |
| Precision | Yes | Accuracy | Yes |
| Guidelines followed | Combination of FDA and EMA | | |

#### Reporting

| | | | |
|---|---|---|---|
| Are reported raw data uploaded into repository? | Available on request | Are metadata available? | Available on request |
| Raw data upload | Available on request | | |

### Sample Descriptions

#### Pooled plasma / Human / Plasma

| | | | |
|---|---|---|---|
| Storage and collection conditions | Available | Temperature handling original sample | Unknown |
| Instant sample preparation | No | Storage temperature | -80 °C |
| Additives | None | | |

#### Pooled erythrocyte-rich fraction / Human / Cells

| | | | |
|---|---|---|---|
| Storage and collection conditions | Available | Temperature handling original sample | Unknown |
| Instant sample preparation | No | Storage temperature | -80 °C |
| Additives | None | | |

### Lipid Class Descriptions

#### 1) TG[M+NH4]+ / Lipid identification

| | | | |
|---|---|---|---|
| Lipid class | TG | MS Level for identification | $MS^1$ |
| Identification level | Species level | $MS^1$ adduct | [M+NH4]+ |

| | | | |
|---|---|---|---|
| Isotope correction at $MS^1$ | Type 2 | $MS^1$ verified by standard | Yes |
| Background check at $MS^1$ | Yes | Did you presume assumptions for identification? | Yes |
| Which assumptions were presumed? | Lipid class separation: coelution of lipids belonging to the same lipid class | Check on: | Isomeric overlap, Isobaric overlap, In-source fragmentation |
| Limit of detection | Signal threshold | RT verified by standard | Yes |
| Separation of isobaric/isomeric interferece confirmed | Yes | Model for separation prediction | No |
| Lipid Identification Software | Homemade | Data manipulation | Centroiding, Lock mass correction, Background subtraction |
| Nomenclature for intact lipid molecule | Yes | | |

#### 1) TG[M+NH4]+ / Lipid quantification

| | | | |
|---|---|---|---|
| Quantitative | Yes | MS Level for quantification | $MS^1$ |

Internal lipid standard(s) $MS^1$

| Internal standard | Endogenous subclass |
|---|---|
| 15:0/18:1-D7/15:0 TG | TG |
| 20:0/20:0/20:0 TG | TG |

| | | | |
|---|---|---|---|
| Type of quantification | Internal standard amount | Response correction | No |
| Type I isotope correction | Yes | Limit of quantification | 2.85 nmol/mL |
| Normalization to reference | No | Lipid Quantification Software | Homemade |
| Batch correction | No | | |

#### 2) PC[M+H]+ / Lipid identification

| | | | |
|---|---|---|---|
| Lipid class | PC | MS Level for identification | $MS^1$ |
| Identification level | Species level | $MS^1$ adduct | [M+H]+ |
| Isotope correction at $MS^1$ | Type 2 | $MS^1$ verified by standard | Yes |
| Background check at $MS^1$ | Yes | Did you presume assumptions for identification? | Yes |
| Which assumptions were presumed? | Lipid class separation: coelution of lipids belonging to the same lipid class | Check on: | Isomeric overlap, Isobaric overlap |
| Limit of detection | Signal threshold | RT verified by standard | Yes |
| Separation of isobaric/isomeric interferece confirmed | Yes | Model for separation prediction | No |
| Lipid Identification Software | Homemade | Data manipulation | Centroiding, Lock mass correction, Background subtraction |
| Nomenclature for intact lipid molecule | Yes | | |

#### 2) PC[M+H]+ / Lipid quantification

| | | | |
|---|---|---|---|
| Quantitative | Yes | MS Level for quantification | $MS^1$ |

Internal lipid standard(s) $MS^1$

| Internal standard | Endogenous subclass |
|---|---|
| 15:0/18:1-D7 PC | PC |

| | |
|---|---|
| 14:0/14:0 PC | PC |

| | | | |
|---|---|---|---|
| Type of quantification | Internal standard amount | Response correction | No |
| Type I isotope correction | Yes | Limit of quantification | 3.75 nmol/mL |
| Normalization to reference | No | Lipid Quantification Software | Homemade |
| Batch correction | No | | |

#### 3) DG[M-H2O+H]+ / Lipid identification

| | | | |
|---|---|---|---|
| Lipid class | DG | MS Level for identification | $MS^1$ |
| Identification level | Species level | $MS^1$ adduct | [M-H2O+H]+ |
| Isotope correction at $MS^1$ | Type 2 | $MS^1$ verified by standard | Yes |
| Background check at $MS^1$ | Yes | Did you presume assumptions for identification? | Yes |
| Which assumptions were presumed? | Lipid class separation: coelution of lipids belonging to the same lipid class | Check on: | Isomeric overlap, Isobaric overlap, In-source fragmentation |
| Limit of detection | Signal threshold | RT verified by standard | Yes |
| Separation of isobaric/isomeric interferece confirmed | Yes | Model for separation prediction | No |
| Lipid Identification Software | Homemade | Data manipulation | Centroiding, Lock mass correction, Background subtraction |
| Nomenclature for intact lipid molecule | Yes | | |

#### 3) DG[M-H2O+H]+ / Lipid quantification

| | | | |
|---|---|---|---|
| Quantitative | Yes | MS Level for quantification | $MS^1$ |

Internal lipid standard(s) $MS^1$

| Internal standard | Endogenous subclass |
|---|---|
| 15:0/18:1-D7 DG | DG |
| 18:1/18:1-D5 DG | DG |

| | | | |
|---|---|---|---|
| Type of quantification | Internal standard amount | Response correction | No |
| Type I isotope correction | Yes | Limit of quantification | 0.75 nmol/mL |
| Normalization to reference | No | Lipid Quantification Software | Homemade |
| Batch correction | No | | |

#### 4) PC[M+CH3COO]- / Lipid identification

| | | | |
|---|---|---|---|
| Lipid class | PC | MS Level for identification | $MS^1$ |
| Identification level | Species level | $MS^1$ adduct | [M+CH3COO]- |
| Isotope correction at $MS^1$ | Type 2 | $MS^1$ verified by standard | Yes |
| Background check at $MS^1$ | Yes | Did you presume assumptions for identification? | Yes |
| Which assumptions were presumed? | Lipid class separation: coelution of lipids belonging to the same lipid class | Check on: | Isomeric overlap, Isobaric overlap |

| | | | |
|---|---|---|---|
| Limit of detection | Signal threshold | RT verified by standard | Yes |
| Separation of isobaric/isomeric interferece confirmed | Yes | Model for separation prediction | No |
| Lipid Identification Software | Homemade | Data manipulation | Centroiding, Lock mass correction, Background subtraction |
| Nomenclature for intact lipid molecule | Yes | | |

#### 4) PC[M+CH3COO]- / Lipid quantification

| | | | |
|---|---|---|---|
| Quantitative | Yes | MS Level for quantification | $MS^1$ |

Internal lipid standard(s) $MS^1$

| Internal standard | Endogenous subclass |
|---|---|
| 15:0/18:1-D7 PC | PC |
| 14:0/14:0 PC | PC |

| | | | |
|---|---|---|---|
| Type of quantification | Internal standard amount | Response correction | No |
| Type I isotope correction | Yes | Limit of quantification | 0.938 nmol/mL |
| Normalization to reference | No | Lipid Quantification Software | Homemade |
| Batch correction | No | | |

#### 5) MG[M+H]+ / Lipid identification

| | | | |
|---|---|---|---|
| Lipid class | MG | MS Level for identification | $MS^1$ |
| Identification level | Species level | $MS^1$ adduct | [M+H]+ |
| Isotope correction at $MS^1$ | Type 2 | $MS^1$ verified by standard | Yes |
| Background check at $MS^1$ | Yes | Did you presume assumptions for identification? | Yes |
| Which assumptions were presumed? | Lipid class separation: coelution of lipids belonging to the same lipid class | Check on: | Isomeric overlap, Isobaric overlap, In-source fragmentation |
| Limit of detection | Signal threshold | RT verified by standard | Yes |
| Separation of isobaric/isomeric interference confirmed | Yes | Model for separation prediction | No |
| Lipid Identification Software | Homemade | Data manipulation | Centroiding, Lock mass correction, Background subtraction |
| Nomenclature for intact lipid molecule | Yes | | |

#### 5) MG[M+H]+ / Lipid quantification

| | | | |
|---|---|---|---|
| Quantitative | Yes | MS Level for quantification | $MS^1$ |

Internal lipid standard(s) $MS^1$

| Internal standard | Endogenous subclass |
|---|---|
| 18:1-D7 MG | MG |
| 19:1 MG | MG |

| | | | |
|---|---|---|---|
| Type of quantification | Internal standard amount | Response correction | No |
| Type I isotope correction | Yes | Limit of quantification | 0.625 nmol/mL |
| Normalization to reference | No | Lipid Quantification Software | Homemade |

| | |
|---|---|
| Batch correction | No |

#### 6) CAR[M+H]+ / Lipid identification

| | | | |
|---|---|---|---|
| Lipid class | CAR | MS Level for identification | $MS^1$ |
| Identification level | Species level | $MS^1$ adduct | [M+H]+ |
| Isotope correction at $MS^1$ | Type 2 | $MS^1$ verified by standard | Yes |
| Background check at $MS^1$ | Yes | Did you presume assumptions for identification? | Yes |
| Which assumptions were presumed? | Lipid class separation: coelution of lipids belonging to the same lipid class | Check on: | Isomeric overlap, Isobaric overlap |
| Limit of detection | Signal threshold | RT verified by standard | Yes |
| Separation of isobaric/isomeric interference confirmed | Yes | Model for separation prediction | No |
| Lipid Identification Software | Homemade | Data manipulation | Centroiding, Lock mass correction, Background subtraction |
| Nomenclature for intact lipid molecule | Yes | | |

#### 6) CAR[M+H]+ / Lipid quantification

| | | | |
|---|---|---|---|
| Quantitative | Yes | MS Level for quantification | $MS^1$ |

Internal lipid standard(s) $MS^1$

| Internal standard | Endogenous subclass |
|---|---|
| 14:0-D9 CAR | CAR |
| 24:0-D4 CAR | CAR |

| | | | |
|---|---|---|---|
| Type of quantification | Internal standard amount | Response correction | No |
| Type I isotope correction | Yes | Limit of quantification | 0.015 nmol/mL |
| Normalization to reference | No | Lipid Quantification Software | Homemade |
| Batch correction | No | | |

#### 7) NAE[M-H]- / Lipid identification

| | | | |
|---|---|---|---|
| Lipid class | NAE | MS Level for identification | $MS^1$ |
| Identification level | Species level | $MS^1$ adduct | [M-H]- |
| Isotope correction at $MS^1$ | Type 2 | $MS^1$ verified by standard | Yes |
| Background check at $MS^1$ | Yes | Did you presume assumptions for identification? | Yes |
| Which assumptions were presumed? | Lipid class separation: coelution of lipids belonging to the same lipid class | Check on: | Isomeric overlap, Isobaric overlap |
| Limit of detection | Signal threshold | RT verified by standard | Yes |
| Separation of isobaric/isomeric interference confirmed | Yes | Model for separation prediction | No |
| Lipid Identification Software | Homemade | Data manipulation | Centroiding, Lock mass correction, Background subtraction |

| | |
|---|---|
| Nomenclature for intact lipid molecule | Yes |

#### 7) NAE[M-H]- / Lipid quantification

| | | | |
|---|---|---|---|
| Quantitative | Yes | MS Level for quantification | $MS^1$ |

Internal lipid standard(s) $MS^1$

| Internal standard | Endogenous subclass |
|---|---|
| 17:1 NAE | NAE |

| | | | |
|---|---|---|---|
| Type of quantification | Internal standard amount | Response correction | No |
| Type I isotope correction | Yes | Limit of quantification | 2.5 nmol/mL |
| Normalization to reference | No | Lipid Quantification Software | Homemade |
| Batch correction | No | | |

#### 8) PC P[M+H]+ / Lipid identification

| | | | |
|---|---|---|---|
| Lipid class | PC P | MS Level for identification | $MS^1$ |
| Identification level | Species level | $MS^1$ adduct | [M+H]+ |
| Isotope correction at $MS^1$ | Type 2 | $MS^1$ verified by standard | Yes |
| Background check at $MS^1$ | Yes | Did you presume assumptions for identification? | Yes |
| Which assumptions were presumed? | Lipid class separation: coelution of lipids belonging to the same lipid class | Check on: | Isomeric overlap, Isobaric overlap |
| Limit of detection | Signal threshold | RT verified by standard | Yes |
| Separation of isobaric/isomeric interferece confirmed | Yes | Model for separation prediction | No |
| Lipid Identification Software | Homemade | Data manipulation | Centroiding, Lock mass correction, Background subtraction |
| Nomenclature for intact lipid molecule | Yes | | |

#### 8) PC P[M+H]+ / Lipid quantification

| | | | |
|---|---|---|---|
| Quantitative | Yes | MS Level for quantification | $MS^1$ |

Internal lipid standard(s) $MS^1$

| Internal standard | Endogenous subclass |
|---|---|
| 18:0/18:1-D9 PC P- | PC |

| | | | |
|---|---|---|---|
| Type of quantification | Internal standard amount | Response correction | No |
| Type I isotope correction | Yes | Limit of quantification | 3.75 nmol/mL |
| Normalization to reference | No | Lipid Quantification Software | Homemade |
| Batch correction | No | | |

#### 9) PC P[M+CH3COO]- / Lipid identification

| | | | |
|---|---|---|---|
| Lipid class | PC P | MS Level for identification | $MS^1$ |
| Identification level | Species level | $MS^1$ adduct | [M+CH3COO]- |
| Isotope correction at $MS^1$ | Type 2 | $MS^1$ verified by standard | Yes |
| Background check at $MS^1$ | Yes | Did you presume assumptions for identification? | Yes |
| Which assumptions were presumed? | Lipid class separation: coelution of lipids belonging to the same lipid class | Check on: | Isomeric overlap, Isobaric overlap |
| Limit of detection | Signal threshold | RT verified by standard | Yes |
| Separation of isobaric/isomeric interferece confirmed | Yes | Model for separation prediction | No |
| Lipid Identification Software | Homemade | Data manipulation | Centroiding, Lock mass correction, Background subtraction |
| Nomenclature for intact lipid molecule | Yes | | |

#### 9) PC P[M+CH3COO]- / Lipid quantification

| | | | |
|---|---|---|---|
| Quantitative | Yes | MS Level for quantification | $MS^1$ |

Internal lipid standard(s) $MS^1$

| Internal standard | Endogenous subclass |
|---|---|
| 18:0/18:1-D9 PC P- | PC |

| | | | |
|---|---|---|---|
| Type of quantification | Internal standard amount | Response correction | No |
| Type I isotope correction | Yes | Limit of quantification | 1.00 nmol/mL |
| Normalization to reference | No | Lipid Quantification Software | Homemade |
| Batch correction | No | | |

#### 10) FA[M-H]- / Lipid identification

| | | | |
|---|---|---|---|
| Lipid class | FA | MS Level for identification | $MS^1$ |
| Identification level | Species level | $MS^1$ adduct | [M-H]- |
| Isotope correction at $MS^1$ | Type 2 | $MS^1$ verified by standard | Yes |
| Background check at $MS^1$ | Yes | Did you presume assumptions for identification? | Yes |
| Which assumptions were presumed? | Lipid class separation: coelution of lipids belonging to the same lipid class | Check on: | Isomeric overlap, Isobaric overlap, In-source fragmentation |
| Limit of detection | Signal threshold | RT verified by standard | Yes |
| Separation of isobaric/isomeric interferece confirmed | Yes | Model for separation prediction | No |
| Lipid Identification Software | Homemade | Data manipulation | Centroiding, Lock mass correction, Background subtraction |
| Nomenclature for intact lipid molecule | Yes | | |

#### 10) FA[M-H]- / Lipid quantification

| | | | |
|---|---|---|---|
| Quantitative | Yes | MS Level for quantification | $MS^1$ |

Internal lipid standard(s) $MS^1$

| Internal standard | Endogenous subclass |
|---|---|

| | |
|---|---|
| 18:1-D9 FA | FA |
| 13:0 FA | FA |

| | | | |
|---|---|---|---|
| Type of quantification | Internal standard amount | Response correction | No |
| Type I isotope correction | Yes | Limit of quantification | 0.25 nmol/mL |
| Normalization to reference | No | Lipid Quantification Software | Homemade |
| Batch correction | No | | |

#### 11) LPC[M+H]+ / Lipid identification

| | | | |
|---|---|---|---|
| Lipid class | LPC | MS Level for identification | $MS^1$ |
| Identification level | Species level | $MS^1$ adduct | [M+H]+ |
| Isotope correction at $MS^1$ | Type 2 | $MS^1$ verified by standard | Yes |
| Background check at $MS^1$ | Yes | Did you presume assumptions for identification? | Yes |
| Which assumptions were presumed? | Lipid class separation: coelution of lipids belonging to the same lipid class | Check on: | Isomeric overlap, Isobaric overlap, In-source fragmentation |
| Limit of detection | Signal threshold | RT verified by standard | Yes |
| Separation of isobaric/isomeric interferece confirmed | Yes | Model for separation prediction | No |
| Lipid Identification Software | Homemade | Data manipulation | Centroiding, Lock mass correction, Background subtraction |
| Nomenclature for intact lipid molecule | Yes | | |

#### 11) LPC[M+H]+ / Lipid quantification

| | | | |
|---|---|---|---|
| Quantitative | Yes | MS Level for quantification | $MS^1$ |

Internal lipid standard(s) $MS^1$

| Internal standard | Endogenous subclass |
|---|---|
| 18:1-D7 LPC | LPC |
| 13:0 LPC | LPC |

| | | | |
|---|---|---|---|
| Type of quantification | Internal standard amount | Response correction | No |
| Type I isotope correction | Yes | Limit of quantification | 0.875 nmol/mL |
| Normalization to reference | No | Lipid Quantification Software | Homemade |
| Batch correction | No | | |

#### 12) LPC[M+CH3COO]- / Lipid identification

| | | | |
|---|---|---|---|
| Lipid class | LPC | MS Level for identification | $MS^1$ |
| Identification level | Species level | $MS^1$ adduct | [M+CH3COO]- |
| Isotope correction at $MS^1$ | Type 2 | $MS^1$ verified by standard | Yes |
| Background check at $MS^1$ | Yes | Did you presume assumptions for identification? | Yes |

| | | | |
|---|---|---|---|
| Which assumptions were presumed? | Lipid class separation: coelution of lipids belonging to the same lipid class | Check on: | Isomeric overlap, Isobaric overlap, In-source fragmentation |
| Limit of detection | Signal threshold | RT verified by standard | Yes |
| Separation of isobaric/isomeric interferece confirmed | Yes | Model for separation prediction | No |
| Lipid Identification Software | Homemade | Data manipulation | Centroiding, Lock mass correction, Background subtraction |
| Nomenclature for intact lipid molecule | Yes | | |

#### 12) LPC[M+CH3COO]- / Lipid quantification

| | | | |
|---|---|---|---|
| Quantitative | Yes | MS Level for quantification | $MS^1$ |

Internal lipid standard(s) $MS^1$

| Internal standard | Endogenous subclass |
|---|---|
| 18:1-D7 LPC | LPC |
| 13:0 LPC | LPC |

| | | | |
|---|---|---|---|
| Type of quantification | Internal standard amount | Response correction | No |
| Type I isotope correction | Yes | Limit of quantification | 0.875 nmol/mL |
| Normalization to reference | No | Lipid Quantification Software | Homemade |
| Batch correction | No | | |

#### 13) PE[M+H]+ / Lipid identification

| | | | |
|---|---|---|---|
| Lipid class | PE | MS Level for identification | $MS^1$ |
| Identification level | Species level | $MS^1$ adduct | [M+H]+ |
| Isotope correction at $MS^1$ | Type 2 | $MS^1$ verified by standard | Yes |
| Background check at $MS^1$ | Yes | Did you presume assumptions for identification? | Yes |
| Which assumptions were presumed? | Lipid class separation: coelution of lipids belonging to the same lipid class | Check on: | Isomeric overlap, Isobaric overlap |
| Limit of detection | Signal threshold | RT verified by standard | Yes |
| Separation of isobaric/isomeric interferece confirmed | Yes | Model for separation prediction | No |
| Lipid Identification Software | Homemade | Data manipulation | Centroiding, Lock mass correction, Background subtraction |
| Nomenclature for intact lipid molecule | Yes | | |

#### 13) PE[M+H]+ / Lipid quantification

| | | | |
|---|---|---|---|
| Quantitative | Yes | MS Level for quantification | $MS^1$ |

Internal lipid standard(s) $MS^1$

| Internal standard | Endogenous subclass |
|---|---|
| 15:0/18:1-D7 PE | PE |
| 14:0/14:0 PE | PE |

| | | | |
|---|---|---|---|
| Type of quantification | Internal standard amount | Response correction | No |

| | | | |
|---|---|---|---|
| Type I isotope correction | Yes | Limit of quantification | 1.5 nmol/mL |
| Normalization to reference | No | Lipid Quantification Software | Homemade |
| Batch correction | No | | |

#### 14) PE[M-H]- / Lipid identification

| | | | |
|---|---|---|---|
| Lipid class | PE | MS Level for identification | $MS^1$ |
| Identification level | Species level | $MS^1$ adduct | [M-H]- |
| Isotope correction at $MS^1$ | Type 2 | $MS^1$ verified by standard | Yes |
| Background check at $MS^1$ | Yes | Did you presume assumptions for identification? | Yes |
| Which assumptions were presumed? | Lipid class separation: coelution of lipids belonging to the same lipid class | Check on: | Isomeric overlap, Isobaric overlap |
| Limit of detection | Signal threshold | RT verified by standard | Yes |
| Separation of isobaric/isomeric interferece confirmed | Yes | Model for separation prediction | No |
| Lipid Identification Software | Homemade | Data manipulation | Centroiding, Lock mass correction, Background subtraction |
| Nomenclature for intact lipid molecule | Yes | | |

#### 14) PE[M-H]- / Lipid quantification

| | | | |
|---|---|---|---|
| Quantitative | Yes | MS Level for quantification | $MS^1$ |

Internal lipid standard(s) $MS^1$

| Internal standard | Endogenous subclass |
|---|---|
| 15:0/18:1-D7 PE | PE |
| 14:0/14:0 PE | PE |

| | | | |
|---|---|---|---|
| Type of quantification | Internal standard amount | Response correction | No |
| Type I isotope correction | Yes | Limit of quantification | 0.15 nmol/mL |
| Normalization to reference | No | Lipid Quantification Software | Homemade |
| Batch correction | No | | |

#### 15) PE P[M+H]+ / Lipid identification

| | | | |
|---|---|---|---|
| Lipid class | PE P | MS Level for identification | $MS^1$ |
| Identification level | Species level | $MS^1$ adduct | [M+H]+ |
| Isotope correction at $MS^1$ | Type 2 | $MS^1$ verified by standard | Yes |
| Background check at $MS^1$ | Yes | Did you presume assumptions for identification? | Yes |
| Which assumptions were presumed? | Lipid class separation: coelution of lipids belonging to the same lipid class | Check on: | Isomeric overlap, Isobaric overlap |
| Limit of detection | Signal threshold | RT verified by standard | Yes |
| Separation of isobaric/isomeric interferece confirmed | Yes | Model for separation prediction | No |

| | | | |
|---|---|---|---|
| Lipid Identification Software | Homemade | Data manipulation | Centroiding, Lock mass correction, Background subtraction |
| Nomenclature for intact lipid molecule | Yes | | |

#### 15) PE P[M+H]+ / Lipid quantification

| | | | |
|---|---|---|---|
| Quantitative | Yes | MS Level for quantification | $MS^1$ |

Internal lipid standard(s) $MS^1$

| Internal standard | Endogenous subclass |
|---|---|
| 18:0/18:1-D9 PE P- | PE P- |

| | | | |
|---|---|---|---|
| Type of quantification | Internal standard amount | Response correction | No |
| Type I isotope correction | Yes | Limit of quantification | 0.5 nmol/mL |
| Normalization to reference | No | Lipid Quantification Software | Homemade |
| Batch correction | No | | |

#### 16) PE P[M-H]- / Lipid identification

| | | | |
|---|---|---|---|
| Lipid class | PE P | MS Level for identification | $MS^1$ |
| Identification level | Species level | $MS^1$ adduct | [M-H]- |
| Isotope correction at $MS^1$ | Type 2 | $MS^1$ verified by standard | Yes |
| Background check at $MS^1$ | Yes | Did you presume assumptions for identification? | Yes |
| Which assumptions were presumed? | Lipid class separation: coelution of lipids belonging to the same lipid class | Check on: | Isomeric overlap, Isobaric overlap |
| Limit of detection | Signal threshold | RT verified by standard | Yes |
| Separation of isobaric/isomeric interferece confirmed | Yes | Model for separation prediction | No |
| Lipid Identification Software | Homemade | Data manipulation | Centroiding, Lock mass correction, Background subtraction |
| Nomenclature for intact lipid molecule | Yes | | |

#### 16) PE P[M-H]- / Lipid quantification

| | | | |
|---|---|---|---|
| Quantitative | Yes | MS Level for quantification | $MS^1$ |

Internal lipid standard(s) $MS^1$

| Internal standard | Endogenous subclass |
|---|---|
| 18:0/18:1-D9 PE P- | PE P- |

| | | | |
|---|---|---|---|
| Type of quantification | Internal standard amount | Response correction | No |
| Type I isotope correction | Yes | Limit of quantification | 0.05 nmol/mL |
| Normalization to reference | No | Lipid Quantification Software | Homemade |
| Batch correction | No | | |

#### 17) LPE[M+H]+ / Lipid identification

| | | | |
|---|---|---|---|
| Lipid class | LPE | MS Level for identification | $MS^1$ |
| Identification level | Species level | $MS^1$ adduct | [M+H]+ |
| Isotope correction at $MS^1$ | Type 2 | $MS^1$ verified by standard | Yes |
| Background check at $MS^1$ | Yes | Did you presume assumptions for identification? | Yes |
| Which assumptions were presumed? | Lipid class separation: coelution of lipids belonging to the same lipid class | Check on: | Isomeric overlap, Isobaric overlap, In-source fragmentation |
| Limit of detection | Signal threshold | RT verified by standard | Yes |
| Separation of isobaric/isomeric interferece confirmed | Yes | Model for separation prediction | No |
| Lipid Identification Software | Homemade | Data manipulation | Centroiding, Lock mass correction, Background subtraction |
| Nomenclature for intact lipid molecule | Yes | | |

#### 17) LPE[M+H]+ / Lipid quantification

| | | | |
|---|---|---|---|
| Quantitative | Yes | MS Level for quantification | $MS^1$ |

Internal lipid standard(s) $MS^1$

| Internal standard | Endogenous subclass |
|---|---|
| 18:1-D7 LPE | LPE |
| 14:0 LPE | LPE |

| | | | |
|---|---|---|---|
| Type of quantification | Internal standard amount | Response correction | No |
| Type I isotope correction | Yes | Limit of quantification | 1.5 nmol/mL |
| Normalization to reference | No | Lipid Quantification Software | Homemade |
| Batch correction | No | | |

#### 18) LPE[M-H]- / Lipid identification

| | | | |
|---|---|---|---|
| Lipid class | LPE | MS Level for identification | $MS^1$ |
| Identification level | Species level | $MS^1$ adduct | [M-H]- |
| Isotope correction at $MS^1$ | Type 2 | $MS^1$ verified by standard | Yes |
| Background check at $MS^1$ | Yes | Did you presume assumptions for identification? | Yes |
| Which assumptions were presumed? | Lipid class separation: coelution of lipids belonging to the same lipid class | Check on: | Isomeric overlap, Isobaric overlap, In-source fragmentation |
| Limit of detection | Signal threshold | RT verified by standard | Yes |
| Separation of isobaric/isomeric interferece confirmed | Yes | Model for separation prediction | No |
| Lipid Identification Software | Homemade | Data manipulation | Centroiding, Lock mass correction, Background subtraction |
| Nomenclature for intact lipid molecule | Yes | | |

#### 18) LPE[M-H]- / Lipid quantification

| | | | |
|---|---|---|---|
| Quantitative | Yes | MS Level for quantification | $MS^1$ |

Internal lipid standard(s) $MS^1$

| Internal standard | Endogenous subclass |
|---|---|
| 18:1-D7 LPE | LPE |
| 14:0 LPE | LPE |

| | | | |
|---|---|---|---|
| Type of quantification | Internal standard amount | Response correction | No |
| Type I isotope correction | Yes | Limit of quantification | 0.075 nmol/mL |
| Normalization to reference | No | Lipid Quantification Software | Homemade |
| Batch correction | No | | |

#### 19) PS[M-H]- / Lipid identification

| | | | |
|---|---|---|---|
| Lipid class | PS | MS Level for identification | $MS^1$ |
| Identification level | Species level | $MS^1$ adduct | [M-H]- |
| Isotope correction at $MS^1$ | Type 2 | $MS^1$ verified by standard | Yes |
| Background check at $MS^1$ | Yes | Did you presume assumptions for identification? | Yes |
| Which assumptions were presumed? | Lipid class separation: coelution of lipids belonging to the same lipid class | Check on: | Isomeric overlap, Isobaric overlap |
| Limit of detection | Signal threshold | RT verified by standard | Yes |
| Separation of isobaric/isomeric interference confirmed | Yes | Model for separation prediction | No |
| Lipid Identification Software | Homemade | Data manipulation | Centroiding, Lock mass correction, Background subtraction |
| Nomenclature for intact lipid molecule | Yes | | |

#### 19) PS[M-H]- / Lipid quantification

| | | | |
|---|---|---|---|
| Quantitative | Yes | MS Level for quantification | $MS^1$ |

Internal lipid standard(s) $MS^1$

| Internal standard | Endogenous subclass |
|---|---|
| 15:0/18:1-D7 PS | PS |
| 14:0/14:0 PS | PS |

| | | | |
|---|---|---|---|
| Type of quantification | Internal standard amount | Response correction | No |
| Type I isotope correction | Yes | Limit of quantification | 12.5 nmol/mL |
| Normalization to reference | No | Lipid Quantification Software | Homemade |
| Batch correction | No | | |

#### 20) LPC O-/P-[M+H]+ / Lipid identification

| | | | |
|---|---|---|---|
| Lipid class | LPC O-/P- | MS Level for identification | $MS^1$ |

| | | | |
|---|---|---|---|
| Identification level | Species level | $MS^1$ adduct | [M+H]+ |
| Isotope correction at $MS^1$ | Type 2 | $MS^1$ verified by standard | No |
| Background check at $MS^1$ | Yes | Did you presume assumptions for identification? | Yes |
| Which assumptions were presumed? | Lipid class separation: coelution of lipids belonging to the same lipid class | Check on: | Isomeric overlap, Isobaric overlap, In-source fragmentation |
| Limit of detection | Signal threshold | RT verified by standard | No |
| Separation of isobaric/isomeric interferece confirmed | No | Model for separation prediction | No |
| Lipid Identification Software | Homemade | Data manipulation | Centroiding, Lock mass correction, Background subtraction |
| Nomenclature for intact lipid molecule | Yes | | |

#### 20) LPC O-/P-[M+H]+ / Lipid quantification

| | | | |
|---|---|---|---|
| Quantitative | Yes | MS Level for quantification | $MS^1$ |

Internal lipid standard(s) $MS^1$

| Internal standard | Endogenous subclass |
|---|---|
| 18:1-D7 LPC | LPC |
| 13:0 LPC | LPC |

| | | | |
|---|---|---|---|
| Type of quantification | Internal standard amount | Response correction | No |
| Type I isotope correction | Yes | Limit of quantification | 0.075 nmol/mL |
| Normalization to reference | No | Lipid Quantification Software | Homemade |
| Batch correction | No | | |

#### 21) LPE P-/O-[M-H]- / Lipid identification

| | | | |
|---|---|---|---|
| Lipid class | LPE P-/O- | MS Level for identification | $MS^1$ |
| Identification level | Species level | $MS^1$ adduct | [M-H]- |
| Isotope correction at $MS^1$ | Type 2 | $MS^1$ verified by standard | No |
| Background check at $MS^1$ | Yes | Did you presume assumptions for identification? | Yes |
| Which assumptions were presumed? | Lipid class separation: coelution of lipids belonging to the same lipid class | Check on: | Isomeric overlap, Isobaric overlap, In-source fragmentation |
| Limit of detection | Signal threshold | RT verified by standard | No |
| Separation of isobaric/isomeric interferece confirmed | No | Model for separation prediction | No |
| Lipid Identification Software | Homemade | Data manipulation | Centroiding, Lock mass correction, Background subtraction |
| Nomenclature for intact lipid molecule | Yes | | |

#### 21) LPE P-/O-[M-H]- / Lipid quantification

| | | | |
|---|---|---|---|
| Quantitative | Yes | MS Level for quantification | $MS^1$ |

Internal lipid standard(s) $MS^1$

| Internal standard | Endogenous subclass |
|---|---|

| | |
|---|---|
| 18:1-D7 LPE | LPE |
| 13:0 LPE | LPE |

| | | | |
|---|---|---|---|
| Type of quantification | Internal standard amount | Response correction | No |
| Type I isotope correction | Yes | Limit of quantification | 0.075 nmol/mL |
| Normalization to reference | No | Lipid Quantification Software | Homemade |
| Batch correction | No | | |

#### 22) PG[M-H]- / Lipid identification

| | | | |
|---|---|---|---|
| Lipid class | PG | MS Level for identification | $MS^1$ |
| Identification level | Species level | $MS^1$ adduct | [M-H]- |
| Isotope correction at $MS^1$ | Type 2 | $MS^1$ verified by standard | Yes |
| Background check at $MS^1$ | Yes | Did you presume assumptions for identification? | Yes |
| Which assumptions were presumed? | Lipid class separation: coelution of lipids belonging to the same lipid class | Check on: | Isomeric overlap, Isobaric overlap |
| Limit of detection | Signal threshold | RT verified by standard | Yes |
| Separation of isobaric/isomeric interferece confirmed | Yes | Model for separation prediction | No |
| Lipid Identification Software | Homemade | Data manipulation | Centroiding, Lock mass correction, Background subtraction |
| Nomenclature for intact lipid molecule | Yes | | |

#### 22) PG[M-H]- / Lipid quantification

| | | | |
|---|---|---|---|
| Quantitative | Yes | MS Level for quantification | $MS^1$ |

Internal lipid standard(s) $MS^1$

| Internal standard | Endogenous subclass |
|---|---|
| 15:0/18:1-D7 PG | PG |
| 14:0/14:0 PG | PG |

| | | | |
|---|---|---|---|
| Type of quantification | Internal standard amount | Response correction | No |
| Type I isotope correction | Yes | Limit of quantification | 1.25 nmol/mL |
| Normalization to reference | No | Lipid Quantification Software | Homemade |
| Batch correction | No | | |

#### 23) LPG[M-H]- / Lipid identification

| | | | |
|---|---|---|---|
| Lipid class | LPG | MS Level for identification | $MS^1$ |
| Identification level | Species level | $MS^1$ adduct | [M-H]- |
| Isotope correction at $MS^1$ | Type 2 | $MS^1$ verified by standard | Yes |
| Background check at $MS^1$ | Yes | Did you presume assumptions for identification? | Yes |

| | | | |
|---|---|---|---|
| Which assumptions were presumed? | Lipid class separation: coelution of lipids belonging to the same lipid class | Check on: | Isomeric overlap, Isobaric overlap, In-source fragmentation |
| Limit of detection | Signal threshold | RT verified by standard | Yes |
| Separation of isobaric/isomeric interferece confirmed | Yes | Model for separation prediction | No |
| Lipid Identification Software | Homemade | Data manipulation | Centroiding, Lock mass correction, Background subtraction |
| Nomenclature for intact lipid molecule | Yes | | |

#### 23) LPG[M-H]- / Lipid quantification

| | | | |
|---|---|---|---|
| Quantitative | Yes | MS Level for quantification | $MS^1$ |

Internal lipid standard(s) $MS^1$

| Internal standard | Endogenous subclass |
|---|---|
| 17:1 LPG | LPG |
| 14:0 LPG | LPG |

| | | | |
|---|---|---|---|
| Type of quantification | Internal standard amount | Response correction | No |
| Type I isotope correction | Yes | Limit of quantification | 6.25 nmol/mL |
| Normalization to reference | No | Lipid Quantification Software | Homemade |
| Batch correction | No | | |

#### 24) PI[M-H]- / Lipid identification

| | | | |
|---|---|---|---|
| Lipid class | PI | MS Level for identification | $MS^1$ |
| Identification level | Species level | $MS^1$ adduct | [M-H]- |
| Isotope correction at $MS^1$ | Type 2 | $MS^1$ verified by standard | Yes |
| Background check at $MS^1$ | Yes | Did you presume assumptions for identification? | Yes |
| Which assumptions were presumed? | Lipid class separation: coelution of lipids belonging to the same lipid class | Check on: | Isomeric overlap, Isobaric overlap |
| Limit of detection | Signal threshold | RT verified by standard | Yes |
| Separation of isobaric/isomeric interferece confirmed | Yes | Model for separation prediction | No |
| Lipid Identification Software | Homemade | Data manipulation | Centroiding, Lock mass correction, Background subtraction |
| Nomenclature for intact lipid molecule | Yes | | |

#### 24) PI[M-H]- / Lipid quantification

| | | | |
|---|---|---|---|
| Quantitative | Yes | MS Level for quantification | $MS^1$ |

Internal lipid standard(s) $MS^1$

| Internal standard | Endogenous subclass |
|---|---|
| 15:0/18:1-D7 PI | PI |
| 15:0/18:1 PI | PI |

| | | | |
|---|---|---|---|
| Type of quantification | Internal standard amount | Response correction | No |

| | | | |
|---|---|---|---|
| Type I isotope correction | Yes | Limit of quantification | 0.3 nmol/mL |
| Normalization to reference | No | Lipid Quantification Software | Homemade |
| Batch correction | No | | |

#### 25) LPI[M-H]- / Lipid identification

| | | | |
|---|---|---|---|
| Lipid class | LPI | MS Level for identification | $MS^1$ |
| Identification level | Species level | $MS^1$ adduct | [M-H]- |
| Isotope correction at $MS^1$ | Type 2 | $MS^1$ verified by standard | Yes |
| Background check at $MS^1$ | Yes | Did you presume assumptions for identification? | Yes |
| Which assumptions were presumed? | Lipid class separation: coelution of lipids belonging to the same lipid class | Check on: | Isomeric overlap, Isobaric overlap, In-source fragmentation |
| Limit of detection | Signal threshold | RT verified by standard | Yes |
| Separation of isobaric/isomeric interferece confirmed | Yes | Model for separation prediction | No |
| Lipid Identification Software | Homemade | Data manipulation | Centroiding, Lock mass correction, Background subtraction |
| Nomenclature for intact lipid molecule | Yes | | |

#### 25) LPI[M-H]- / Lipid quantification

| | | | |
|---|---|---|---|
| Quantitative | Yes | MS Level for quantification | $MS^1$ |

Internal lipid standard(s) $MS^1$

| Internal standard | Endogenous subclass |
|---|---|
| 17:0-D5 LPI | LPI |
| 15:0-D5 LPI | LPI |

| | | | |
|---|---|---|---|
| Type of quantification | Internal standard amount | Response correction | No |
| Type I isotope correction | Yes | Limit of quantification | 0.15 nmol/mL |
| Normalization to reference | No | Lipid Quantification Software | Homemade |
| Batch correction | No | | |

#### 26) PA[M-H]- / Lipid identification

| | | | |
|---|---|---|---|
| Lipid class | PA | MS Level for identification | $MS^1$ |
| Identification level | Species level | $MS^1$ adduct | [M-H]- |
| Isotope correction at $MS^1$ | Type 2 | $MS^1$ verified by standard | Yes |
| Background check at $MS^1$ | Yes | Did you presume assumptions for identification? | Yes |
| Which assumptions were presumed? | Lipid class separation: coelution of lipids belonging to the same lipid class | Check on: | Isomeric overlap, Isobaric overlap, In-source fragmentation |
| Limit of detection | Signal threshold | RT verified by standard | Yes |
| Separation of isobaric/isomeric interferece confirmed | Yes | Model for separation prediction | No |

| | | | |
|---|---|---|---|
| Lipid Identification Software | Homemade | Data manipulation | Centroiding, Lock mass correction, Background subtraction |
| Nomenclature for intact lipid molecule | Yes | | |

#### 26) PA[M-H]- / Lipid quantification

| | | | |
|---|---|---|---|
| Quantitative | No | Normalization to reference | No |
| Batch correction | No | | |

#### 27) LPS[M-H]- / Lipid identification

| | | | |
|---|---|---|---|
| Lipid class | LPS | MS Level for identification | $MS^1$ |
| Identification level | Species level | $MS^1$ adduct | [M-H]- |
| Isotope correction at $MS^1$ | Type 2 | $MS^1$ verified by standard | Yes |
| Background check at $MS^1$ | Yes | Did you presume assumptions for identification? | Yes |
| Which assumptions were presumed? | Lipid class separation: coelution of lipids belonging to the same lipid class | Check on: | Isomeric overlap, Isobaric overlap, In-source fragmentation |
| Limit of detection | Signal threshold | RT verified by standard | Yes |
| Separation of isobaric/isomeric interferece confirmed | Yes | Model for separation prediction | No |
| Lipid Identification Software | Homemade | Data manipulation | Centroiding, Lock mass correction, Background subtraction |
| Nomenclature for intact lipid molecule | Yes | | |

#### 27) LPS[M-H]- / Lipid quantification

| | | | |
|---|---|---|---|
| Quantitative | No | Normalization to reference | No |
| Batch correction | No | | |

#### 28) CL[M-H]- / Lipid identification

| | | | |
|---|---|---|---|
| Lipid class | CL | MS Level for identification | $MS^1$ |
| Identification level | Species level | $MS^1$ adduct | [M-H]- |
| Isotope correction at $MS^1$ | Type 2 | $MS^1$ verified by standard | Yes |
| Background check at $MS^1$ | Yes | Did you presume assumptions for identification? | Yes |
| Which assumptions were presumed? | Lipid class separation: coelution of lipids belonging to the same lipid class | Check on: | Isomeric overlap, Isobaric overlap |
| Limit of detection | Signal threshold | RT verified by standard | Yes |
| Separation of isobaric/isomeric interferece confirmed | Yes | Model for separation prediction | No |

| | | | |
|---|---|---|---|
| Lipid Identification Software | Homemade | Data manipulation | Centroiding, Lock mass correction, Background subtraction |
| Nomenclature for intact lipid molecule | Yes | | |

#### 28) CL[M-H]- / Lipid quantification

| | | | |
|---|---|---|---|
| Quantitative | Yes | MS Level for quantification | $MS^1$ |

Internal lipid standard(s) $MS^1$

| Internal standard | Endogenous subclass |
|---|---|
| 14:0/14:0/14:0/14:0 CL | CL |

| | | | |
|---|---|---|---|
| Type of quantification | Internal standard amount | Response correction | No |
| Type I isotope correction | Yes | Limit of quantification | 3.125 nmol/mL |
| Normalization to reference | No | Lipid Quantification Software | Homemade |
| Batch correction | No | | |

#### 29) BMP[M-H]- / Lipid identification

| | | | |
|---|---|---|---|
| Lipid class | BMP | MS Level for identification | $MS^1$ |
| Identification level | Species level | $MS^1$ adduct | [M-H]- |
| Isotope correction at $MS^1$ | Type 2 | $MS^1$ verified by standard | Yes |
| Background check at $MS^1$ | Yes | Did you presume assumptions for identification? | Yes |
| Which assumptions were presumed? | Lipid class separation: coelution of lipids belonging to the same lipid class | Check on: | Isomeric overlap, Isobaric overlap, In-source fragmentation |
| Limit of detection | Signal threshold | RT verified by standard | Yes |
| Separation of isobaric/isomeric interferece confirmed | Yes | Model for separation prediction | No |
| Lipid Identification Software | Homemade | Data manipulation | Centroiding, Lock mass correction, Background subtraction |
| Nomenclature for intact lipid molecule | Yes | | |

#### 29) BMP[M-H]- / Lipid quantification

| | | | |
|---|---|---|---|
| Quantitative | Yes | MS Level for quantification | $MS^1$ |

Internal lipid standard(s) $MS^1$

| Internal standard | Endogenous subclass |
|---|---|
| 14:0/14:0 BMP | BMP |

| | | | |
|---|---|---|---|
| Type of quantification | Internal standard amount | Response correction | No |
| Type I isotope correction | Yes | Limit of quantification | 12.5 nmol/mL |
| Normalization to reference | No | Lipid Quantification Software | Homemade |
| Batch correction | No | | |

#### 30) Cer[M+H-H2O]+ / Lipid identification

| | | | |
|---|---|---|---|
| Lipid class | Cer | MS Level for identification | $MS^1$ |
| Identification level | Species level | $MS^1$ adduct | [M+H-H2O]+ |
| Isotope correction at $MS^1$ | Type 2 | $MS^1$ verified by standard | Yes |
| Background check at $MS^1$ | Yes | Did you presume assumptions for identification? | Yes |
| Which assumptions were presumed? | Lipid class separation: coelution of lipids belonging to the same lipid class | Check on: | Isomeric overlap, Isobaric overlap, In-source fragmentation |
| Limit of detection | Signal threshold | RT verified by standard | Yes |
| Separation of isobaric/isomeric interferece confirmed | Yes | Model for separation prediction | No |
| Lipid Identification Software | Homemade | Data manipulation | Centroiding, Lock mass correction, Background subtraction |
| Nomenclature for intact lipid molecule | Yes | | |

#### 30) Cer[M+H-H2O]+ / Lipid quantification

| | | | |
|---|---|---|---|
| Quantitative | Yes | MS Level for quantification | $MS^1$ |

Internal lipid standard(s) $MS^1$

| Internal standard | Endogenous subclass |
|---|---|
| 12:0/18:1;O2 Cer | Cer |
| 18:0/18:1-D7;O2 Cer | Cer |

| | | | |
|---|---|---|---|
| Type of quantification | Internal standard amount | Response correction | No |
| Type I isotope correction | Yes | Limit of quantification | 0.05 nmol/mL |
| Normalization to reference | No | Lipid Quantification Software | Homemade |
| Batch correction | No | | |

#### 31) Cer[M+CH3COO]- / Lipid identification

| | | | |
|---|---|---|---|
| Lipid class | Cer | MS Level for identification | $MS^1$ |
| Identification level | Species level | $MS^1$ adduct | [M+CH3COO]- |
| Isotope correction at $MS^1$ | Type 2 | $MS^1$ verified by standard | Yes |
| Background check at $MS^1$ | Yes | Did you presume assumptions for identification? | Yes |
| Which assumptions were presumed? | Lipid class separation: coelution of lipids belonging to the same lipid class | Check on: | Isomeric overlap, Isobaric overlap, In-source fragmentation |
| Limit of detection | Signal threshold | RT verified by standard | Yes |
| Separation of isobaric/isomeric interferece confirmed | Yes | Model for separation prediction | No |
| Lipid Identification Software | Homemade | Data manipulation | Centroiding, Lock mass correction, Background subtraction |
| Nomenclature for intact lipid molecule | Yes | | |

#### 31) Cer[M+CH3COO]- / Lipid quantification

| | | | |
|---|---|---|---|
| Quantitative | Yes | MS Level for quantification | $MS^1$ |

Internal lipid standard(s) $MS^1$

| Internal standard | Endogenous subclass |
|---|---|
| 12:0/18:1;O2 Cer | Cer |
| 18:0/18:1-D7;O2 Cer | Cer |

| | | | |
|---|---|---|---|
| Type of quantification | Internal standard amount | Response correction | No |
| Type I isotope correction | Yes | Limit of quantification | 0.05 nmol/mL |
| Normalization to reference | No | Lipid Quantification Software | Homemade |
| Batch correction | No | | |

#### 32) CerPE[M-H]- / Lipid identification

| | | | |
|---|---|---|---|
| Lipid class | CerPE | MS Level for identification | $MS^1$ |
| Identification level | Species level | $MS^1$ adduct | [M-H]- |
| Isotope correction at $MS^1$ | Type 2 | $MS^1$ verified by standard | Yes |
| Background check at $MS^1$ | Yes | Did you presume assumptions for identification? | Yes |
| Which assumptions were presumed? | Lipid class separation: coelution of lipids belonging to the same lipid class | Check on: | Isomeric overlap, Isobaric overlap |
| Limit of detection | Signal threshold | RT verified by standard | Yes |
| Separation of isobaric/isomeric interfeence confirmed | Yes | Model for separation prediction | No |
| Lipid Identification Software | Homemade | Data manipulation | Centroiding, Lock mass correction, Background subtraction |
| Nomenclature for intact lipid molecule | Yes | | |

#### 32) CerPE[M-H]- / Lipid quantification

| | | | |
|---|---|---|---|
| Quantitative | Yes | MS Level for quantification | $MS^1$ |

Internal lipid standard(s) $MS^1$

| Internal standard | Endogenous subclass |
|---|---|
| 12:0/17:1;O2 CerPE | CerPE |

| | | | |
|---|---|---|---|
| Type of quantification | Internal standard amount | Response correction | No |
| Type I isotope correction | Yes | Limit of quantification | 6.25 nmol/mL |
| Normalization to reference | No | Lipid Quantification Software | Homemade |
| Batch correction | No | | |

#### 33) HexCer[M+H]+ / Lipid identification

| | | | |
|---|---|---|---|
| Lipid class | HexCer | MS Level for identification | $MS^1$ |
| Identification level | Species level | $MS^1$ adduct | [M+H]+ |

| | | | |
|---|---|---|---|
| Isotope correction at $MS^1$ | Type 2 | $MS^1$ verified by standard | Yes |
| Background check at $MS^1$ | Yes | Did you presume assumptions for identification? | Yes |
| Which assumptions were presumed? | Lipid class separation: coelution of lipids belonging to the same lipid class | Check on: | Isomeric overlap, Isobaric overlap, In-source fragmentation |
| Limit of detection | Signal threshold | RT verified by standard | Yes |
| Separation of isobaric/isomeric interferece confirmed | Yes | Model for separation prediction | No |
| Lipid Identification Software | Homemade | Data manipulation | Centroiding, Lock mass correction, Background subtraction |
| Nomenclature for intact lipid molecule | Yes | | |

#### 33) HexCer[M+H]+ / Lipid quantification

| | | | |
|---|---|---|---|
| Quantitative | Yes | MS Level for quantification | $MS^1$ |

Internal lipid standard(s) $MS^1$

| Internal standard | Endogenous subclass |
|---|---|
| 18:1;O2/12:0 GalCer | HexCer |
| 18:1-D5;O2/18:0 GlucCer | HexCer |

| | | | |
|---|---|---|---|
| Type of quantification | Internal standard amount | Response correction | No |
| Type I isotope correction | Yes | Limit of quantification | 0.75 nmol/mL |
| Normalization to reference | No | Lipid Quantification Software | Homemade |
| Batch correction | No | | |

#### 34) HexCer[M+CH3COO]- / Lipid identification

| | | | |
|---|---|---|---|
| Lipid class | HexCer | MS Level for identification | $MS^1$ |
| Identification level | Species level | $MS^1$ adduct | [M+CH3COO]- |
| Isotope correction at $MS^1$ | Type 2 | $MS^1$ verified by standard | Yes |
| Background check at $MS^1$ | Yes | Did you presume assumptions for identification? | Yes |
| Which assumptions were presumed? | Lipid class separation: coelution of lipids belonging to the same lipid class | Check on: | Isomeric overlap, Isobaric overlap, In-source fragmentation |
| Limit of detection | Signal threshold | RT verified by standard | Yes |
| Separation of isobaric/isomeric interferece confirmed | Yes | Model for separation prediction | No |
| Lipid Identification Software | Homemade | Data manipulation | Centroiding, Lock mass correction, Background subtraction |
| Nomenclature for intact lipid molecule | Yes | | |

#### 34) HexCer[M+CH3COO]- / Lipid quantification

| | | | |
|---|---|---|---|
| Quantitative | Yes | MS Level for quantification | $MS^1$ |

Internal lipid standard(s) $MS^1$

| Internal standard | Endogenous subclass |
|---|---|
| 18:1;O2/12:0 GalCer | HexCer |

| | |
|---|---|
| 18:1-D5;O2/18:0 GlucCer | HexCer |

| | | | |
|---|---|---|---|
| Type of quantification | Internal standard amount | Response correction | No |
| Type I isotope correction | Yes | Limit of quantification | 0.25 nmol/mL |
| Normalization to reference | No | Lipid Quantification Software | Homemade |
| Batch correction | No | | |

#### 35) Hex2Cer[M+H]+ / Lipid identification

| | | | |
|---|---|---|---|
| Lipid class | Hex2Cer | MS Level for identification | $MS^1$ |
| Identification level | Species level | $MS^1$ adduct | [M+H]+ |
| Isotope correction at $MS^1$ | Type 2 | $MS^1$ verified by standard | Yes |
| Background check at $MS^1$ | Yes | Did you presume assumptions for identification? | Yes |
| Which assumptions were presumed? | Lipid class separation: coelution of lipids belonging to the same lipid class | Check on: | Isomeric overlap, Isobaric overlap, In-source fragmentation |
| Limit of detection | Signal threshold | RT verified by standard | Yes |
| Separation of isobaric/isomeric interferece confirmed | Yes | Model for separation prediction | No |
| Lipid Identification Software | Homemade | Data manipulation | Centroiding, Lock mass correction, Background subtraction |
| Nomenclature for intact lipid molecule | Yes | | |

#### 35) Hex2Cer[M+H]+ / Lipid quantification

| | | | |
|---|---|---|---|
| Quantitative | Yes | MS Level for quantification | $MS^1$ |

Internal lipid standard(s) $MS^1$

| Internal standard | Endogenous subclass |
|---|---|
| 18:1;O2/12:0 LacCer | Hex2Cer |
| 18:1-D7;O2/15:0 LacCer | Hex2Cer |

| | | | |
|---|---|---|---|
| Type of quantification | Internal standard amount | Response correction | No |
| Type I isotope correction | Yes | Limit of quantification | 0.6 nmol/mL |
| Normalization to reference | No | Lipid Quantification Software | Homemade |
| Batch correction | No | | |

#### 36) Hex2Cer[M+CH3COO]- / Lipid identification

| | | | |
|---|---|---|---|
| Lipid class | Hex2Cer | MS Level for identification | $MS^1$ |
| Identification level | Species level | $MS^1$ adduct | [M+CH3COO]- |
| Isotope correction at $MS^1$ | Type 2 | $MS^1$ verified by standard | Yes |
| Background check at $MS^1$ | Yes | Did you presume assumptions for identification? | Yes |
| Which assumptions were presumed? | Lipid class separation: coelution of lipids belonging to the same lipid class | Check on: | Isomeric overlap, Isobaric overlap, In-source fragmentation |

| | | | |
|---|---|---|---|
| Limit of detection | Signal threshold | RT verified by standard | Yes |
| Separation of isobaric/isomeric interferece confirmed | Yes | Model for separation prediction | No |
| Lipid Identification Software | Homemade | Data manipulation | Centroiding, Lock mass correction, Background subtraction |
| Nomenclature for intact lipid molecule | Yes | | |

#### 36) Hex2Cer[M+CH3COO]- / Lipid quantification

| | | | |
|---|---|---|---|
| Quantitative | Yes | MS Level for quantification | MS$^1$ |

Internal lipid standard(s) MS$^1$

| Internal standard | Endogenous subclass |
|---|---|
| 18:1;O2/12:0 LacCer | Hex2Cer |
| 18:1-D7;O2/15:0 LacCer | Hex2Cer |

| | | | |
|---|---|---|---|
| Type of quantification | Internal standard amount | Response correction | No |
| Type I isotope correction | Yes | Limit of quantification | 0.3 nmol/mL |
| Normalization to reference | No | Lipid Quantification Software | Homemade |
| Batch correction | No | | |

#### 37) Hex3Cer[M+CH3COO]- / Lipid identification

| | | | |
|---|---|---|---|
| Lipid class | Hex3Cer | MS Level for identification | MS$^1$ |
| Identification level | Species level | MS$^1$ adduct | [M+CH3COO]- |
| Isotope correction at MS$^1$ | Type 2 | MS$^1$ verified by standard | Yes |
| Background check at MS$^1$ | Yes | Did you presume assumptions for identification? | Yes |
| Which assumptions were presumed? | Lipid class separation: coelution of lipids belonging to the same lipid class | Check on: | Isomeric overlap, Isobaric overlap, In-source fragmentation |
| Limit of detection | Signal threshold | RT verified by standard | Yes |
| Separation of isobaric/isomeric interference confirmed | Yes | Model for separation prediction | No |
| Lipid Identification Software | Homemade | Data manipulation | Centroiding, Lock mass correction, Background subtraction |
| Nomenclature for intact lipid molecule | Yes | | |

#### 37) Hex3Cer[M+CH3COO]- / Lipid quantification

| | | | |
|---|---|---|---|
| Quantitative | Yes | MS Level for quantification | MS$^1$ |

Internal lipid standard(s) MS$^1$

| Internal standard | Endogenous subclass |
|---|---|
| 18:1;O2/17:0 Gb3 | Hex3Cer |
| 18:1-D9;O2/16:0 Gb3 | Hex3Cer |

| | | | |
|---|---|---|---|
| Type of quantification | Internal standard amount | Response correction | No |
| Type I isotope correction | Yes | Limit of quantification | 0.1 nmol/mL |
| Normalization to reference | No | Lipid Quantification Software | Homemade |

| | |
|---|---|
| Batch correction | No |

#### 38) Hex4Cer[M+CH3COO]- / Lipid identification

| | | | |
|---|---|---|---|
| Lipid class | Hex4Cer | MS Level for identification | $MS^1$ |
| Identification level | Species level | $MS^1$ adduct | [M+CH3COO]- |
| Isotope correction at $MS^1$ | Type 2 | $MS^1$ verified by standard | No |
| Background check at $MS^1$ | Yes | Did you presume assumptions for identification? | Yes |
| Which assumptions were presumed? | Lipid class separation: coelution of lipids belonging to the same lipid class | Check on: | Isomeric overlap, Isobaric overlap, In-source fragmentation |
| Limit of detection | Signal threshold | RT verified by standard | No |
| Separation of isobaric/isomeric interferece confirmed | No | Model for separation prediction | No |
| Lipid Identification Software | Homemade | Data manipulation | Centroiding, Lock mass correction, Background subtraction |
| Nomenclature for intact lipid molecule | Yes | | |

#### 38) Hex4Cer[M+CH3COO]- / Lipid quantification

| | | | |
|---|---|---|---|
| Quantitative | No | Normalization to reference | No |
| Batch correction | No | | |

#### 39) GM3[M-H]- / Lipid identification

| | | | |
|---|---|---|---|
| Lipid class | GM3 | MS Level for identification | $MS^1$ |
| Identification level | Species level | $MS^1$ adduct | [M-H]- |
| Isotope correction at $MS^1$ | Type 2 | $MS^1$ verified by standard | Yes |
| Background check at $MS^1$ | Yes | Did you presume assumptions for identification? | Yes |
| Which assumptions were presumed? | Lipid class separation: coelution of lipids belonging to the same lipid class | Check on: | Isomeric overlap, Isobaric overlap, In-source fragmentation |
| Limit of detection | Signal threshold | RT verified by standard | Yes |
| Separation of isobaric/isomeric interferece confirmed | Yes | Model for separation prediction | No |
| Lipid Identification Software | Homemade | Data manipulation | Centroiding, Lock mass correction, Background subtraction |
| Nomenclature for intact lipid molecule | Yes | | |

#### 39) GM3[M-H]- / Lipid quantification

| | | | |
|---|---|---|---|
| Quantitative | Yes | MS Level for quantification | $MS^1$ |

Internal lipid standard(s) $MS^1$

| Internal standard | Endogenous subclass |
|---|---|

| | |
|---|---|
| 18:1-D5;O2/18:0 GM3 | GM3 |
| 18:1-D9;O2/16:0 GM3 | GM3 |

| | | | |
|---|---|---|---|
| Type of quantification | Internal standard amount | Response correction | No |
| Type I isotope correction | Yes | Limit of quantification | 0.125 nmol/mL |
| Normalization to reference | No | Lipid Quantification Software | Homemade |
| Batch correction | No | | |

#### 40) SHexCer[M-H]- / Lipid identification

| | | | |
|---|---|---|---|
| Lipid class | SHexCer | MS Level for identification | $MS^1$ |
| Identification level | Species level | $MS^1$ adduct | [M-H]- |
| Isotope correction at $MS^1$ | Type 2 | $MS^1$ verified by standard | Yes |
| Background check at $MS^1$ | Yes | Did you presume assumptions for identification? | Yes |
| Which assumptions were presumed? | Lipid class separation: coelution of lipids belonging to the same lipid class | Check on: | Isomeric overlap, Isobaric overlap, In-source fragmentation |
| Limit of detection | Signal threshold | RT verified by standard | Yes |
| Separation of isobaric/isomeric interference confirmed | Yes | Model for separation prediction | No |
| Lipid Identification Software | Homemade | Data manipulation | Centroiding, Lock mass correction, Background subtraction |
| Nomenclature for intact lipid molecule | Yes | | |

#### 40) SHexCer[M-H]- / Lipid quantification

| | | | |
|---|---|---|---|
| Quantitative | Yes | MS Level for quantification | $MS^1$ |

Internal lipid standard(s) $MS^1$

| Internal standard | Endogenous subclass |
|---|---|
| 18:1;O2/12:0 SHexCer | SHexCer |
| 18:1-D7;O2/13:0 SHexCer | SHexCer |

| | | | |
|---|---|---|---|
| Type of quantification | Internal standard amount | Response correction | No |
| Type I isotope correction | Yes | Limit of quantification | 0.02 nmol/mL |
| Normalization to reference | No | Lipid Quantification Software | Homemade |
| Batch correction | No | | |

#### 41) SM[M+CH3COO]- / Lipid identification

| | | | |
|---|---|---|---|
| Lipid class | SM | MS Level for identification | $MS^1$ |
| Identification level | Species level | $MS^1$ adduct | [M+CH3COO]- |
| Isotope correction at $MS^1$ | Type 2 | $MS^1$ verified by standard | Yes |
| Background check at $MS^1$ | Yes | Did you presume assumptions for identification? | Yes |

| | | | |
|---|---|---|---|
| Which assumptions were presumed? | Lipid class separation: coelution of lipids belonging to the same lipid class | Check on: | Isomeric overlap, Isobaric overlap, In-source fragmentation |
| Limit of detection | Signal threshold | RT verified by standard | Yes |
| Separation of isobaric/isomeric interference confirmed | Yes | Model for separation prediction | No |
| Lipid Identification Software | Homemade | Data manipulation | Centroiding, Lock mass correction, Background subtraction |
| Nomenclature for intact lipid molecule | Yes | | |

#### 41) SM[M+CH3COO]- / Lipid quantification

| | | | |
|---|---|---|---|
| Quantitative | Yes | MS Level for quantification | $MS^1$ |

Internal lipid standard(s) $MS^1$

| Internal standard | Endogenous subclass |
|---|---|
| 18:1;O2/12:0 SM | SM |
| 18:1-D9;O2/18:1 SM | SM |

| | | | |
|---|---|---|---|
| Type of quantification | Internal standard amount | Response correction | No |
| Type I isotope correction | Yes | Limit of quantification | 0.87 nmol/mL |
| Normalization to reference | No | Lipid Quantification Software | Homemade |
| Batch correction | No | | |

#### 42) SHex2Cer[M-H]- / Lipid identification

| | | | |
|---|---|---|---|
| Lipid class | SHex2Cer | MS Level for identification | $MS^1$ |
| Identification level | Species level | $MS^1$ adduct | [M-H]- |
| Isotope correction at $MS^1$ | Type 2 | $MS^1$ verified by standard | No |
| Background check at $MS^1$ | Yes | Did you presume assumptions for identification? | Yes |
| Which assumptions were presumed? | Lipid class separation: coelution of lipids belonging to the same lipid class | Check on: | Isomeric overlap, Isobaric overlap, In-source fragmentation |
| Limit of detection | Signal threshold | RT verified by standard | No |
| Separation of isobaric/isomeric interference confirmed | No | Model for separation prediction | No |
| Lipid Identification Software | Homemade | Data manipulation | Centroiding, Lock mass correction, Background subtraction |
| Nomenclature for intact lipid molecule | Yes | | |

#### 42) SHex2Cer[M-H]- / Lipid quantification

| | | | |
|---|---|---|---|
| Quantitative | No | Normalization to reference | No |
| Batch correction | No | | |

#### 43) ST[M+H-H2O]+ / Lipid identification

| | | | |
|---|---|---|---|
| Lipid class | ST | MS Level for identification | $MS^1$ |
| Identification level | Species level | $MS^1$ adduct | [M+H-H2O]+ |
| Isotope correction at $MS^1$ | Type 2 | $MS^1$ verified by standard | Yes |
| Background check at $MS^1$ | Yes | Did you presume assumptions for identification? | Yes |
| Which assumptions were presumed? | Lipid class separation: coelution of lipids belonging to the same lipid class | Check on: | Isomeric overlap, Isobaric overlap, In-source fragmentation |
| Limit of detection | Signal threshold | RT verified by standard | Yes |
| Separation of isobaric/isomeric interferece confirmed | Yes | Model for separation prediction | No |
| Lipid Identification Software | Homemade | Data manipulation | Centroiding, Lock mass correction, Background subtraction |
| Nomenclature for intact lipid molecule | Yes | | |

#### 43) ST[M+H-H2O]+ / Lipid quantification

| | | | |
|---|---|---|---|
| Quantitative | Yes | MS Level for quantification | $MS^1$ |

Internal lipid standard(s) $MS^1$

| Internal standard | Endogenous subclass |
|---|---|
| Cholesterol D7 | ST |

| | | | |
|---|---|---|---|
| Type of quantification | Internal standard amount | Response correction | No |
| Type I isotope correction | Yes | Limit of quantification | 4.375 nmol/mL |
| Normalization to reference | No | Lipid Quantification Software | Homemade |
| Batch correction | No | | |

#### 44) SE[M+Na]+ / Lipid identification

| | | | |
|---|---|---|---|
| Lipid class | SE | MS Level for identification | $MS^1$ |
| Identification level | Species level | $MS^1$ adduct | [M+Na]+ |
| Isotope correction at $MS^1$ | Type 2 | $MS^1$ verified by standard | Yes |
| Background check at $MS^1$ | Yes | Did you presume assumptions for identification? | Yes |
| Which assumptions were presumed? | Lipid class separation: coelution of lipids belonging to the same lipid class | Check on: | Isomeric overlap, Isobaric overlap, In-source fragmentation |
| Limit of detection | Signal threshold | RT verified by standard | Yes |
| Separation of isobaric/isomeric interferece confirmed | Yes | Model for separation prediction | No |
| Lipid Identification Software | Homemade | Data manipulation | Centroiding, Lock mass correction, Background subtraction |
| Nomenclature for intact lipid molecule | Yes | | |

#### 44) SE[M+Na]+ / Lipid quantification

| | | | |
|---|---|---|---|
| Quantitative | Yes | MS Level for quantification | $MS^1$ |

Internal lipid standard(s) $MS^1$

| Internal standard | Endogenous subclass |
|---|---|
| 16:0 CE-D7 | SE |
| 20:0 CE | SE |

| | | | |
|---|---|---|---|
| Type of quantification | Internal standard amount | Response correction | Response model |
| Type I isotope correction | Yes | Limit of quantification | 50 nmol/mL |
| Normalization to reference | No | Lipid Quantification Software | Homemade |
| Batch correction | No | | |

#### 45) PR[M+H]+ / Lipid identification

| | | | |
|---|---|---|---|
| Lipid class | PR | MS Level for identification | $MS^1$ |
| Identification level | Species level | $MS^1$ adduct | [M+H]+ |
| Isotope correction at $MS^1$ | Type 2 | $MS^1$ verified by standard | Yes |
| Background check at $MS^1$ | Yes | Did you presume assumptions for identification? | Yes |
| Which assumptions were presumed? | Lipid class separation: coelution of lipids belonging to the same lipid class | Check on: | Isomeric overlap, Isobaric overlap, In-source fragmentation |
| Limit of detection | Signal threshold | RT verified by standard | Yes |
| Separation of isobaric/isomeric interferece confirmed | Yes | Model for separation prediction | No |
| Lipid Identification Software | Homemade | Data manipulation | Centroiding, Lock mass correction, Background subtraction |
| Nomenclature for intact lipid molecule | Yes | | |

#### 45) PR[M+H]+ / Lipid quantification

| | | | |
|---|---|---|---|
| Quantitative | Yes | MS Level for quantification | $MS^1$ |

Internal lipid standard(s) $MS^1$

| Internal standard | Endogenous subclass |
|---|---|
| α-tocopherol D6 | PR |

| | | | |
|---|---|---|---|
| Type of quantification | Internal standard amount | Response correction | No |
| Type I isotope correction | Yes | Limit of quantification | 6.25 nmol/mL |
| Normalization to reference | No | Lipid Quantification Software | Homemade |
| Batch correction | No | | |

#### 46) PR[M-H]- / Lipid identification

| | | | |
|---|---|---|---|
| Lipid class | PR | MS Level for identification | $MS^1$ |
| Identification level | Species level | $MS^1$ adduct | [M-H]- |
| Isotope correction at $MS^1$ | Type 2 | $MS^1$ verified by standard | Yes |
| Background check at $MS^1$ | Yes | Did you presume assumptions for identification? | Yes |

| | | | |
|---|---|---|---|
| Which assumptions were presumed? | Lipid class separation: coelution of lipids belonging to the same lipid class | Check on: | Isomeric overlap, Isobaric overlap, In-source fragmentation |
| Limit of detection | Signal threshold | RT verified by standard | Yes |
| Separation of isobaric/isomeric interferece confirmed | Yes | Model for separation prediction | No |
| Lipid Identification Software | Homemade | Data manipulation | Centroiding, Lock mass correction, Background subtraction |
| Nomenclature for intact lipid molecule | Yes | | |

#### 46) PR[M-H]- / Lipid quantification

| | | | |
|---|---|---|---|
| Quantitative | Yes | MS Level for quantification | $MS^1$ |

Internal lipid standard(s) $MS^1$

| Internal standard | Endogenous subclass |
|---|---|
| α-tocopherol D6 | PR |

| | | | |
|---|---|---|---|
| Type of quantification | Internal standard amount | Response correction | No |
| Type I isotope correction | Yes | Limit of quantification | 18.75 nmol/mL |
| Normalization to reference | No | Lipid Quantification Software | Homemade |
| Batch correction | No | | |

#### 47) NA[M-H]- / Lipid identification

| | | | |
|---|---|---|---|
| Lipid class | NA | MS Level for identification | $MS^1$ |
| Identification level | Species level | $MS^1$ adduct | [M-H]- |
| Isotope correction at $MS^1$ | Type 2 | $MS^1$ verified by standard | Yes |
| Background check at $MS^1$ | Yes | Did you presume assumptions for identification? | Yes |
| Which assumptions were presumed? | Lipid class separation: coelution of lipids belonging to the same lipid class | Check on: | Isomeric overlap, Isobaric overlap |
| Limit of detection | Signal threshold | RT verified by standard | Yes |
| Separation of isobaric/isomeric interferece confirmed | No | Model for separation prediction | No |
| Lipid Identification Software | Homemade | Data manipulation | Centroiding, Lock mass correction, Background subtraction |
| Nomenclature for intact lipid molecule | Yes | | |

#### 47) NA[M-H]- / Lipid quantification

| | | | |
|---|---|---|---|
| Quantitative | No | Normalization to reference | No |
| Batch correction | No | | |
